# The CRL7^FBXW8^ Complex Controls the Mammary Stem Cell Compartment Through Regulation of NUMB Levels

**DOI:** 10.1101/2025.02.27.640530

**Authors:** Simone Sabbioni, Maria Grazia Filippone, Letizia Amadori, Stefano Confalonieri, Roberta Bonfanti, Stefano Capoano, Ivan Nicola Colaluca, Stefano Freddi, Giovanni Bertalot, Giovanni Fagà, Elisa Zagarrí, Mario Varasi, Rosalind Helen Gunby, Ciro Mercurio, Salvatore Pece, Pier Paolo Di Fiore, Daniela Tosoni

## Abstract

NUMB is a tumor suppressor gene that functions by inhibiting the action of the NOTCH proto-oncogene and enhancing the levels and activity of the tumor suppressor protein p53. In breast cancer (BC), NUMB loss-of-function (LOF), mediated by various molecular mechanisms, is a frequent and causal event. Herein, we establish that loss of NUMB protein, resulting from protein hyper-degradation, is the prevalent mechanism of NUMB LOF in BC. Through a RNAi-based screening, we identified the CRL7^FBXW8^ complex as the E3 ligase complex responsible for NUMB hyper-degradation in BC. Genetic and pharmacological inhibition of CRL7^FBXW8^ rescued the transformation-related phenotypes induced by NUMB LOF in BC cell lines and in patient-derived xenografts. These effects were directly dependent on the restoration of NUMB protein levels. Thus, enhanced CRL7^FBXW8^ activity, through its interference with the tumor suppressor activity of NUMB, is a causal alteration in BC, suggesting it as a potential therapeutic target for precision medicine.

## 1. Introduction

NUMB is a tumor suppressor gene implicated in various cancers, including breast, prostate, lung, stomach, liver and others.^[1]^ It was originally identified in the fruit fly as a cell fate determinant which, through its asymmetric partitioning during cell mitosis, determines the fate of the daughter cells in a cell-autonomous fashion.^[2]^ In the fruit fly, the role of NUMB is to counteract the action of the membrane signaling receptor NOTCH,^[3]^ through a mechanism involving its canonical function as an endocytic regulator.^[4]^ While this function is conserved throughout evolution,^[5]^ in chordates – including mammals – the molecular mechanisms governing cell fate regulation by NUMB are more diverse owing to the incorporation of an alternatively spliced exon (exon 3) in the *NUMB* transcript. This exon encodes an 11-amino acid sequence that enables NUMB to bind and inhibit MDM2, the major E3 ligase responsible for the ubiquitination and degradation of p53.^[1a, 1b, 6]^ Consequently, the downregulation of NUMB results in reduced levels of p53.^[1a, 1b]^ This specific function of NUMB is crucial for maintaining homeostasis of the adult stem cell (SC) compartment in various organs, as has been well-established in the mammary gland.^[1e, 1h, 1m]^ NUMB loss-of-function (LOF) causes profound alterations in SC division, shifting it from a predominantly asymmetric mode (1 SC - > 1 SC + 1 committed progenitor) to a symmetric mode (1 SC -> 2 SCs). The upshot of these events is an expansion of the SC population, a hallmark of the cancer SC (CSC) compartment.

In summary, NUMB plays an important physiological role, keeping the action of NOTCH (a proto-oncogene) in check while sustaining the activity of p53 (a tumor suppressor). It follows that NUMB LOF is expected to have a significant impact on oncogenic transformation, as clearly demonstrated in breast cancer (BC) where this alteration determines an aggressive disease course.^[1a, 1b, 1e]^

Various mechanisms lead to NUMB LOF in BC: i) reduction in NUMB protein levels due its hyper-degradation, as evidenced in a limited number of clinical cases,^[1r]^ ii) alterations in *NUMB* splicing resulting in downregulation of exon 3-containing isoforms that regulate p53,^[1b]^ iii) aberrant NUMB phosphorylation leading to loss of its asymmetric partitioning and functional inactivation.^[1e]^ The first mechanism, involving NUMB hyper-degradation, has been associated with heightened ubiquitination,^[1r]^ catalyzed by as yet uncharacterized molecular players.

Protein ubiquitination proceeds in a step-wise fashion,^[7]^ beginning with an E1 enzyme that binds to and activates ubiquitin (Ub). Subsequently, the Ub moiety is transferred to an E2 enzyme and the ubiquitination reaction is then completed by an E3 Ub-ligase. There are at least four classes of E3 ligase: HECT type, U-box type, RING-finger type, and RBR type.^[8]^ The HECT and RING-finger type E3s have been most extensively studied. HECT-E3s function as *bona fide* enzymes, receiving Ub from E2 and transferring it to the substrate.^[7]^ In contrast, RING-finger E3s serve as a scaffold for an E2∼Ub intermediate and a protein substrate, facilitating the direct ubiquitination of the latter by the E2 enzyme. In this latter case, the coordinated activity of the E2 and E3 enzymes is necessary to promote the elongation of the Ub chain with high processivity.^[8]^

Cullin-RING Ligases (CRLs) are a subfamily of RING-finger ligases,^[9]^ consisting of two core subunits: a cullin, of which there are nine different types in human (CUL1, CUL2, CUL3, CUL4a, CUL4b, CUL5, CUL7, CUL9, AP2), and a RING-domain partner (RBX1, RBX2, APC11) that binds to the E2∼Ub intermediate.^[10]^ Cullins also interact with substrate-binding receptors, often in the form of multiprotein complexes containing a cullin-binding protein and a substrate-binding protein, frequently from the F-box protein family.^[9, 11]^ Of interest to this study is the atypical CRL complex, CRL7^FBXW8^. The recent structural resolution of CRL7^FBXW8^ revealed the unexpected finding that the complex contains both CUL7 and CUL1 subunits, which together contribute to the formation of a fully catalytically competent complex (see also Figure S1a, Supporting Information).^[10]^ The distinctive feature of CRL7^FBXW8^ is its exceptional specificity for the F-box protein FBXW8, at variance with other CRLs that utilize various F-box proteins or other substrate receptors.^[12]^

Herein, we report that the CRL7^FBXW8^ E3 ligase complex is responsible for aberrant NUMB degradation in BC. Furthermore, we demonstrate that the inhibition of this complex in BC cell lines and patient-derived xenografts (PDXs) selectively affects those exhibiting loss of NUMB protein expression. In these BC models, CRL7^FBXW8^/proteasome inhibition is sufficient to rescue NUMB expression, inhibit tumor growth, and revert the defects in the SC compartment, ultimately leading to a reduction in the number of tumor-initiating cells (TICs). Thus, the CRL7^FBXW8^ complex controls the mammary SC compartment through its regulation of NUMB levels, making it a promising therapeutic target in NUMB-deficient BCs.

## 2. Results

### 2.1 Downregulation of NUMB protein expression is a common alteration in BC associated with poor prognosis

We aimed to establish the significance of NUMB protein expression loss in the natural history of BC. To accomplish this goal and to compare the frequency of this alteration with other mechanisms affecting NUMB,^[1b, 1e]^ we developed a platform for the orthogonal analysis of clinical data, protein expression, and mRNA expression in BC patients. We utilized a BC case-cohort (N=890), previously employed to establish the role of exon 3 loss and NUMB hyperphosphorylation as causal events in BC (henceforth, the IEO Cohort).^[1b, 1e]^ Within this cohort, we conducted a comprehensive analysis of NUMB expression on whole formalin-fixed paraffin-embedded (FFPE) slices using immunohistochemistry (IHC) and RNAseq.

By IHC, we consistently observed high levels of NUMB expression in the normal mammary gland. On a semiquantitative scale ranging from 0 to 3 (in 0.5-point increments), this corresponded to a value of ≥ 2.0 in at least 70% of normal mammary cells (see Figure S1b, Supporting Information, and ^[1r]^). In contrast, BCs displayed variable NUMB protein expression, ranging from completely negative to normal-like staining (Figure S1b, Supporting Information). We classified BCs with an IHC score ≥ 2.0 (as observed in the normal mammary gland) in at least 70% of tumor cells as NUMB-proficient, while the remainder were labeled as NUMB-deficient. According to this classification system, 48.8% of all BCs were identified as NUMB-deficient. This NUMB-deficient status correlated with increased mortality due to BC in both univariate and multivariable analyses across the entire IEO Cohort and in the subgroup of Luminal BCs (Figure 1a,b).

**Figure 1.**
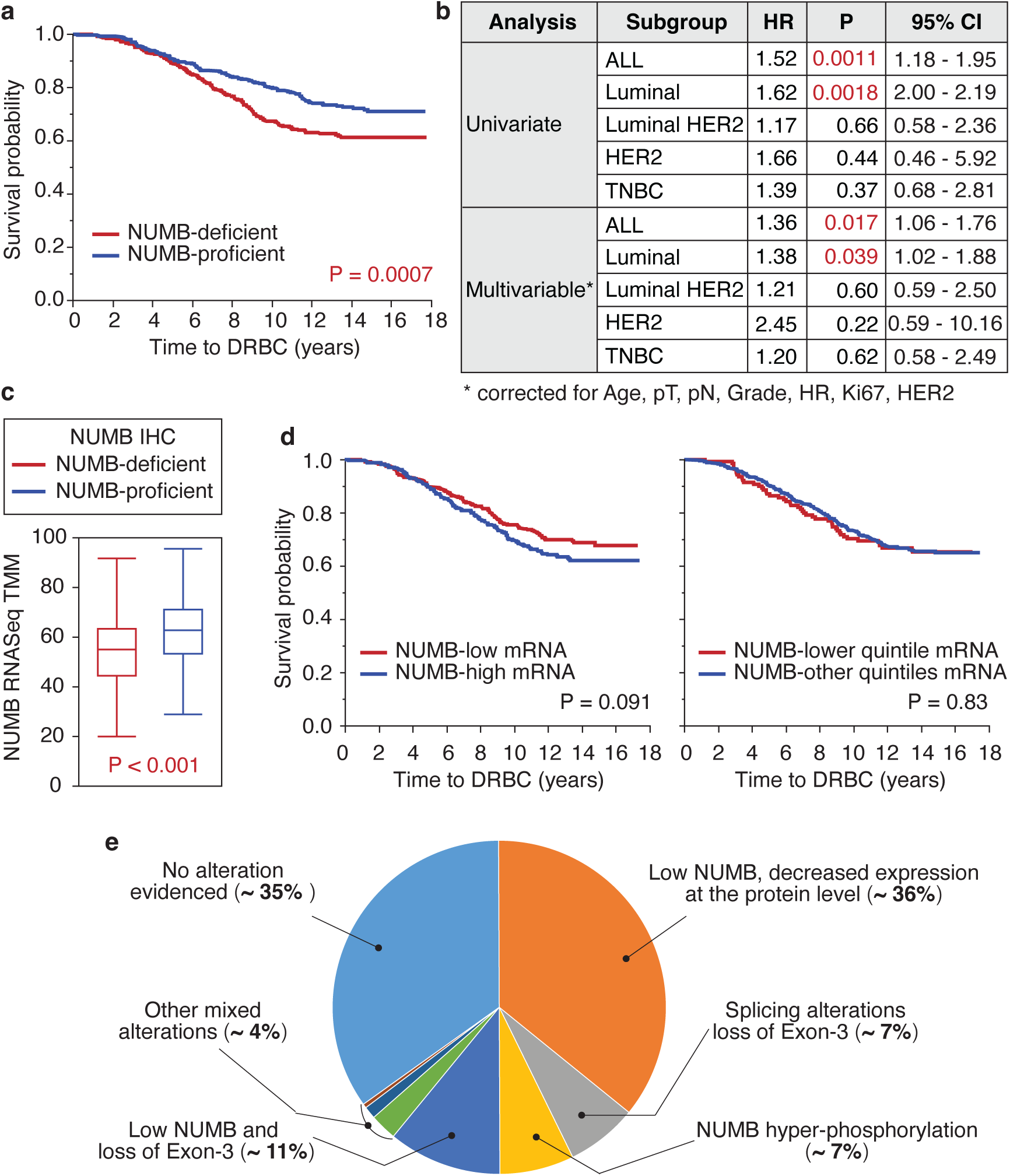
Loss of NUMB expression at the protein level in BCs correlates with prognosis. **a.** Kaplan-Meier (KM) survival analysis of the IEO Cohort. Time to death related to BC (DRBC) was determined in patients stratified by NUMB IHC status (deficient or proficient). Reliable IHC data was obtained for 808 patients. **b.** Univariate and multivariable analysis of the BCs analyzed in ‘a’. HR, hazard ratio. **c.** Relative *NUMB* mRNA levels, determined by TMM (trimmed mean of M values) normalization of RNAseq data, in Numb-deficient and NUMB-proficient BCs, defined by IHC, as described in ‘a’. The analysis was performed on 716 patients for whom reliable IHC and RNAseq data were available. P was calculated by the non-parametric Wilcoxon test using JMP. **d.** KM survival analysis of the 716 patients defined in ‘c’, stratified by levels of *NUMB* mRNA expression determined by RNAseq. Left, BCs were classified as *NUMB*-low or *NUMB*-high relative to the average *NUMB* mRNA expression across the entire cohort. Right, BCs were stratified based on the quintiles of *NUMB* mRNA levels. The Cox proportional hazards model was used to calculate the Hazard Ratio and corresponding p-values. **e.** Pie chart showing the relative contribution of different NUMB LOF mechanisms in BC. Data on NUMB protein levels, expression of Ex3-containing NUMB isoforms, and NUMB hyper-phosphorylation in the 890-case cohort were extracted from this paper and our previous publications.^[1b, 1e]^ In 683 cases, data were retrievable for all three categories (missing samples were mainly due to loss of tissue-microarray cores in the “NUMB hyper-phosphorylation” category^[1e]^). Low mRNA expression of Ex3-containing isoforms occurred in the presence (11% of cases) or absence (7% of cases) of decreased NUMB protein expression. The cumulative frequency of NUMB alterations was ∼ 65%. Other mixed alterations: low NUMB + NUMB hyperphosphorylation (2.6%), low Ex3-containing isoforms + NUMB hyperphosphorylation (1.2%), all three alterations together (0.4%).

Integration of the IHC data with *NUMB* mRNA data obtained by RNAseq revealed that NUMB-deficient BCs (as defined by IHC) exhibited a statistically significant, albeit minor, reduction in the levels of *NUMB* mRNA: average *NUMB* TMM was 56 ± 17 and 63 ± 16 in NUMB-deficient *vs*. NUMB-proficient BCs, respectively (Figure 1c). This small difference at the mRNA level appears insufficient to account for the substantial variations in NUMB expression observed at the protein level by IHC. In support of this notion, *NUMB* mRNA levels were not predictive of disease outcome in the IEO Cohort (Figure 1d) and the METABRIC BC cohort (Figure S1c, Supporting Information).^[13]^

We concluded that loss of NUMB protein in BC is primarily driven by alterations occurring at the post-transcriptional/post-translational levels. This finding is consistent with our previous study in a limited number of BC cases, which demonstrated that NUMB hyper-degradation following excessive ubiquitination is responsible for its loss of expression.^[1r]^ Furthermore, our work highlights loss of NUMB protein as the most frequent NUMB alteration correlating with prognostic outcomes in BC (Figure 1e). We, therefore, sought to uncover the molecular mechanisms governing heightened NUMB protein degradation in BCs.

### 2.2 The CRL7^FBXW8^ complex is responsible for NUMB hyper-degradation in BC cell lines

To investigate NUMB hyper-degradation, we selected suitable *in vitro* models from a screening of established BC cell lines: NUMB-deficient MDA-MB-361 cells expressing low NUMB levels rescuable by treatment with the proteasome inhibitor Bortezomib (BTZ), and NUMB-proficient MDA-MB-231 cells exhibiting high basal NUMB levels insensitive to BTZ (Figure 2a). NUMB levels in MDA-MB-361 cells could also be rescued by treatment with MLN4924 (MLN) (Figure 2a), an inhibitor of the NEDD8-activating enzyme (NAE), which prevents the activation of CRL complexes by inhibiting cullin NEDDylation.^[14]^ We verified comparable *NUMB* mRNA levels in both cell lines (Figure 2b, left) that were unaffected by BTZ or MLN treatment (Figure 2b, right). The restoration of NUMB by MLN in MDA-MB-361 cells suggests the involvement of a CRL ligase, as HECT-type ligases are insensitive to this inhibitor. BTZ and MLN treatments were also efficacious in 3D organoid cultures of the selected cell lines (Figure 2c).

**Figure 2.**
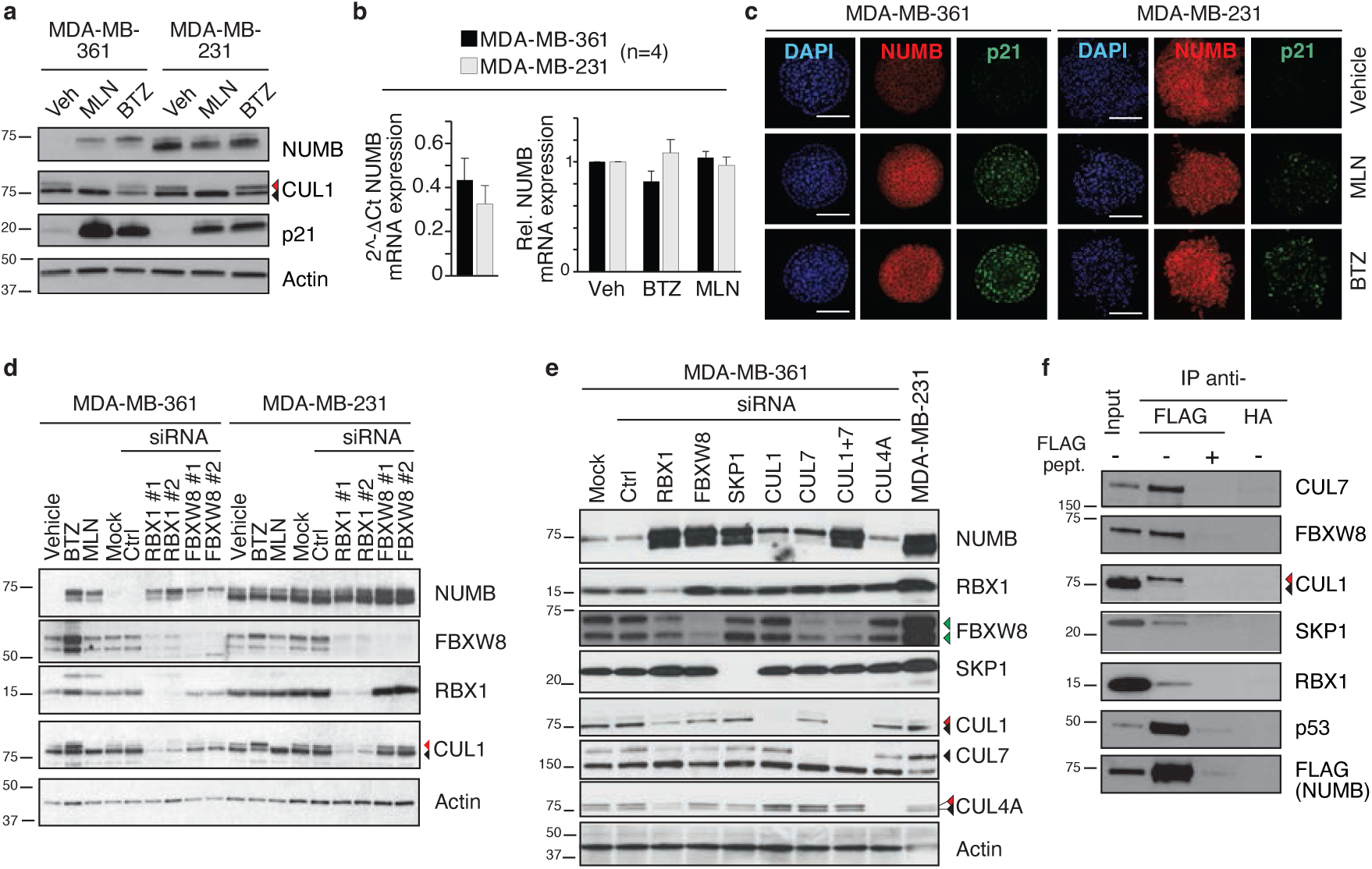
The CRL7^FBXW8^ complex is responsible for NUMB degradation in a NUMB-deficient BC cell line. **a.** MDA-MB-361 and MDA-MB-231 BC cell lines were treated for 24 h with Bortezomib (BTZ, 20 nM), MLN4924 (MLN, 0.5 μM), or vehicle (Veh) and effects on NUMB levels was determined by immunoblotting (IB). p21 was used as a control for proteasome inhibition by BTZ. MLN efficacy was controlled by assessing the levels of NEDDylated CUL1 (upper band, red arrowhead) compared with non-NEDDylated CUL1 (lower band, black arrowhead). Actin, loading control. Results are representative of three independent experiments. In this and all subsequent figures: MW markers for IB are shown in KDa. **b.** *NUMB* mRNA levels in MDA-MB-361 or MDA-MB-231 cells determined by RT-qPCR. Left, basal expression (expressed as 2^^-ΔCt^); right, mRNA levels upon treatment with BTZ or MLN as in ‘a’. In the right panel, the 2^^-ΔCt^ method was used to assess relative NUMB expression, using Veh-treated cells as the reference sample. Data are expressed as mean ± SE (normalized to Veh in the right panel) from 4 independent experiments. No statistically significant differences were observed in the unpaired t-test. **c.** Organotypic cultures of MDA-MB-361 or MDAMB-231 cells were established in Matrigel and treated for ∼24 h with 20 nM BTZ, 0.5 μM MLN or Veh when organoids reached ∼40 μm in diameter. Organoids were fixed and analyzed for NUMB and p21 expression by immunofluorescence (IF). Nuclei were counterstained with DAPI. Results are representative of two independent experiments. Bar = 100 μm. **d.** Transient knockdown (KD) of RBX1 or FBXW8 in MDA-MB-361 and MDA-MB-231 cells using different siRNA oligos (#1, #2). As negative controls cells were transfected with non-targeting control (Ctrl) siRNA, or mock transfected. BTZ and MLN treatments (as in panel a) were used as positive controls for NUMB restoration. IB was performed with the indicated Ab. The arrows in the CUL1 blot are as in ‘a’. For FBXW8, the two bands represent different isoforms. Actin, loading control. **e.** Individual CRL7^FBXW8^ complex components were transiently silenced in MDA-MB-361 cells. For CUL1 and CUL4A, the arrowheads point to NEDDylated (red) and non-NEDDylated (black) forms. For FBXW8, green arrowheads point to the two isoforms. For CUL7, the black arrowhead points to the protein (the lower band visible in the blot is non-specific). MDA-MB-231 cells were used as a reference for “normal” NUMB expression levels. **f.** FLAG-tagged NUMB was transfected into HEK293 cells. Lysates were immunoprecipitated (IP) with anti-FLAG, with or without FLAG peptide to displace the specific interactions, or with anti-HA as an additional negative control. Interacting proteins were detected by IB. p53, a known NUMB interactor, served as positive control. For CUL1, arrowheads point to NEDDylated (red) and non-NEDDylated (black) forms. Note that in HEK293 cells, FBXW8 migrates as a single band. Input, 15 μg; IP, 8 mg. Results are representative of 3 independent experiments.

To identify the components of the human ubiquitin conjugation machinery involved in NUMB hyper-degradation, we performed a high-throughput siRNA screening coupled with a NUMB capture ELISA assay in NUMB-deficient MDA-MB-361 cells (see Supporting Methods, Supporting Information; Table S1, Supporting Information, and Figure S2a, Supporting Information). By adopting stringent filters for significance and reproducibility, we found that RBX1 and FBXW8 represented the top two hits (Figure 2b, Supporting Information). These proteins are part of the CRL7^FBXW8^ complex which contains CUL7, CUL1, SKP1, RBX1, and FBXW8 (Figure S1a, Supporting Information).^[10]^ This complex is inhibitable by MLN, in line with our results (Figure 2a-c).

To validate these hits, we silenced their expression in MDA-MB-361 cells using two different siRNAs and observed the efficient restoration of NUMB levels (Figure 2d). In contrast, no effects of RBX1 knockdown (KD) or FBXW8 KD were evidenced in MDA-MB-231 cells (Figure 2d). Notably, RBX1 KD also reduced the levels of FBXW8 and CUL1 (Figure 2d, see also Figures 2e, 4a, 6e), suggesting a role of RBX1 in promoting the stability of these proteins likely through the formation of a CRL complex.

To obtain further evidence of the involvement of the CRL7^FBXW8^ complex in NUMB degradation, we examined the effects of silencing each of its components individually. In MDA-MB-361 cells, we found that NUMB degradation was inhibited by each of the single KDs: CUL7, CUL1, SKP1, RBX1, and FBXW8 (Figure 2e). In contrast, the KD of CUL4A, not involved in the CRL7^FBXW8^ complex, had no effect on NUMB degradation (Figure 2e). Of note, single KD of CUL1 and CUL7 only partially rescued NUMB levels, while their double KD fully restored NUMB (Figure 2e). These results suggest some level of redundant, yet synergic, action between these cullins in the CRL7^FBXW8^ complex, as discussed below. Furthermore, the silencing of the CRL7^FBXW8^ complex, in MDA-MB-361 cells, had functional consequences on biochemical pathways known to be regulated by NUMB, congruent with the expected results of the restoration of NUMB levels (Figure S3, Supporting Information).

To establish the physical interaction between NUMB and the CRL7^FBXW8^ complex components, we performed co-immunoprecipitation experiments. Given the low levels of NUMB in MDA-MB-361 cells, we used HEK293 cells transiently transfected with a FLAG-tagged NUMB construct. NUMB-FLAG co-immunoprecipitated with all the components of the CRL7^FBXW8^ complex indicating the physical interaction of these proteins (Figure 2f). Moreover, results suggest that within the complex, CUL1 is mainly in its NEDDylated active form.

We next performed NUMB ubiquitination assays *in vivo*. In anti-NUMB immunoprecipitates, ubiquitinated NUMB (detected in IB with anti-Ub) was readily detectable in MDA-MB-361 cells, but scarcely visible in MDA-MB-231 cells (Figure 3a), compatible with all previous results. NUMB ubiquitination in MDA-MB-361 cells was significantly reduced by treatment with MLN (Figure 3a), confirming that the responsible ligase is a NEDD8-dependent one. Importantly, the silencing of RBX1 or FBXW8 reduced significantly NUMB ubiquitination *in vivo* (Figure 3b).

**Figure 3.**
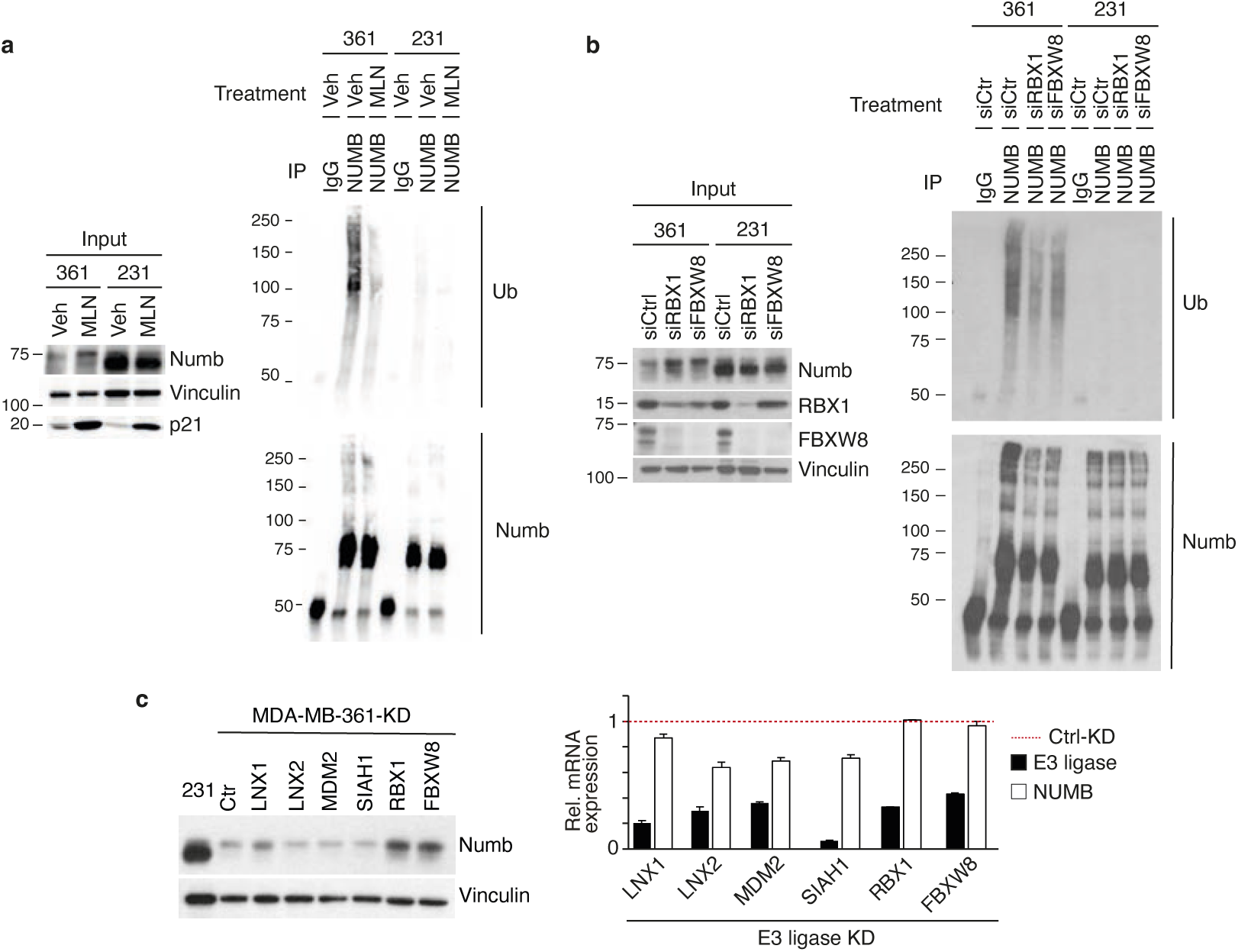
NUMB polyubiquitination in BC cell lines requires the CRL7^FBXW8^ complex. **a.** MDA-MB-361 (361) and MDA-MB-231 (231) cells were treated for 24 h with MLN4924 (MLN, 0.5 μM), or vehicle (Veh). Lysates were then immunoprecipitated (IP, Abs are shown on the top) and immunoblotted (IB, as shown on the right). Input, 10 μg; IP, 6 and 3 mg for MDA-MB-361 and MDA-MB-231, respectively. **b.** MDA-MB-361 and MDA-MB-231 BC cells were transiently silenced for RBX1 or FBXW8 or with a non-targeting control (Ctrl) siRNA. Lysates were IP (as shown on the top) and IB (as shown on the right). Input, 10 μg; IP, 6 and 3 mg for MDA-MB-361 and MDA-MB-231, respectively. **c.** MDA-MB-361 cells were stably transduced with the doxycycline (DOX)-inducible shRNA constructs (pTRIPZ-based vectors) targeting the indicated E3 ligases, and then treated with DOX. Left, IB was performed with the indicated Abs. MDA-MB-231 cells (231) were used as a Numb-proficient control. Vinculin, loading control. Results are representative of 2 independent experiments. Right, RT-qPCR analysis of the expression levels of KD target genes (black bars) and NUMB (white bars), relative to the Ctrl KD (dashed red line = 1). Data are the mean ± SD of technical triplicates.

Finally, several E3 ligases, such as LNX1/2, MDM2, and SIAH1, have been linked in the literature to NUMB ubiquitination and degradation.^[15]^ While none of these enzymes emerged as a significant hit in our high-throughput siRNA screening, we directly excluded their involvement in the ubiquitination/degradation of NUMB in MDA-MB-361 cells since, upon their silencing, there was no significant increase in NUMB levels (Figure 3c).

Together, these findings suggest that the CRL7^FBXW8^ complex binds to NUMB and is responsible for its hyper-degradation in NUMB-deficient MDA-MB-361 BC cells.

### 2.3 CRL7^FBXW8^-mediated NUMB degradation is essential for maintaining CSC phenotypes *in vitro*

To investigate the biological relevance of NUMB degradation by the CRL7^FBXW8^ complex, we employed 3D culture assays that represent *in vitro* readouts of SC properties and tumorigenicity: i) the organoid Matrigel assay, which measures the ability of cells to undergo self-renewal and give rise to differentiated progeny that generate organotypic structures,^[16]^ and ii) the methylcellulose colony assay, which measures the ability of single cells to undergo clonal expansion, i.e., with self-renewal and clonogenic potential, to form cell spheroids.^[17]^

We first analyzed the effects of RBX1 or FBXW8 silencing on MDA-MB-361 and MDA-MB-231 cells. To achieve long-lasting protein ablation, we engineered cell lines stably expressing doxycycline (DOX)-inducible shRNAs against RBX1 and FBXW8 (Figure S4a, Supporting Information). In MDA-MB-361 transduced cells, DOX treatment resulted in the stable KD of RBX1 or FBXW8, accompanied by an increase in NUMB protein levels (Figure 4a,b and Figure S4b, Supporting Information). These molecular changes were accompanied by a significant reduction in the growth potential of MDA-MB-361 cells in 3D cultures, as evidenced by a decrease in organoid-forming efficiency (OFE) in Matrigel and spheroid-forming efficiency (SFE) in methylcellulose (Figure 4c). In contrast, no effects were observed in transduced MDA-MB-231 cells (Figure 4d). Notably, despite RBX1 being involved in multiple CRL complexes, the growth inhibitory effects upon its KD were similar in amplitude to those of FBXW8 KD. Since it has been shown that FBXW8 participates exclusively in the CRL7^FBXW8^ complex,^[12]^ we reasoned that the observed biological consequences of RBX1 KD can be attributed largely to CRL7^FBXW8^ inhibition rather than the inhibition of other CRL complexes. Furthermore, we demonstrated that the selective inhibition of MDA-MB-361 cell growth in 3D cultures was directly linked to NUMB restoration resulting from RBX1 or FBXW8 KD, since it was abolished by the simultaneous KD of NUMB (Figure 4e,f and Figure S4b,c, Supporting Information). These results support a scenario where CRL7^FBXW8^-mediated NUMB degradation is causally linked to the growth potential of the CSC population of NUMB-deficient BC cells.

**Figure 4.**
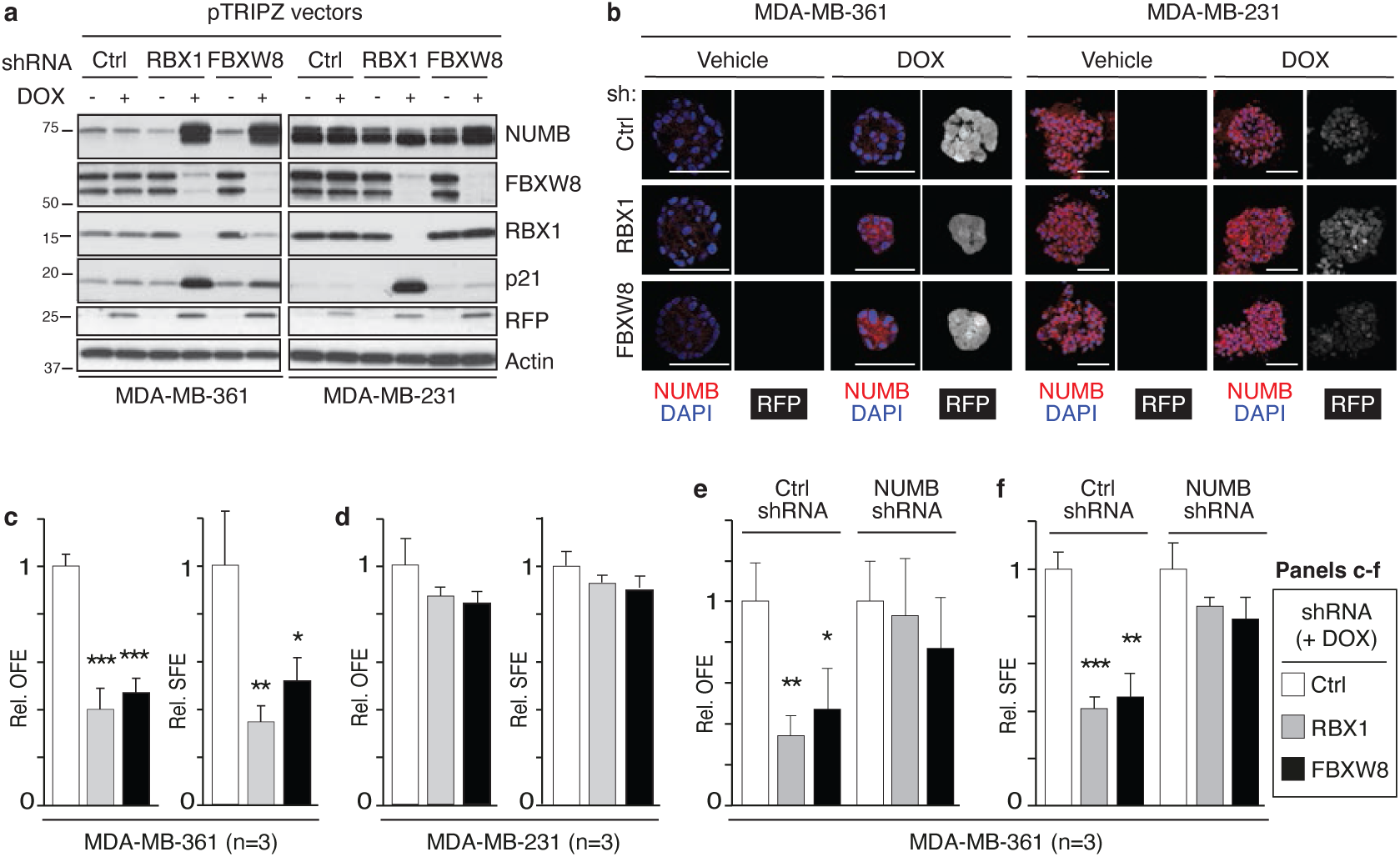
The CRL7^FBXW8^ complex modulates NUMB-dependent biological activities. **a.** MDA-MB-361 and MDA-MB-231 cells were stably transduced with the indicated doxycycline (DOX)-inducible shRNA constructs (pTRIPZ-based vectors) and treated or not with DOX. IB was performed with the indicated Ab. RFP is a reporter expressed upon DOX stimulation, used as a control for shRNA induction efficiency. p21, positive control for RBX1 KD. Actin, loading control. The MDA-MB-361 and MDA-MB-231 blots come from a single larger blot in which irrelevant lanes were spliced out. The NUMB blots represent different exposures (shorter exposure for MDA-MB-231 because of high endogenous NUMB levels). **b.** Representative images of a 3D Matrigel assay performed with MDA-MB-361 (left) and MDA-MB-231 (right) cells, silenced with the indicated shRNAs. After fixation, organoids were stained with anti-NUMB (pseudo-colored in red) and DAPI (blue). RFP (pseudo-colored in gray), reporter of DOX-induced shRNA expression. Bar, 50 μm. In MDA-MB-361 cells, the reduction in organoid size following RBX1 or FBXW8 KD was consistent with the increased NUMB levels. Data are representative of two independent biological replicates. **c,d.** MDA-MB-361 (c) or MDA-MB-231 (d) cells stably transduced with the DOX-inducible shRNA pTRIPZ constructs as in ‘a’, were analyzed for organoid-forming efficiency (OFE) in Matrigel or spheroid-forming efficiency (SFE) in methylcellulose, in the presence of DOX. **e,f.** MDA-MB-361 cells were stably silenced for NUMB with a DOX-inducible shRNA construct (pTRIPZ-vector) and further silenced for RBX1 or FBXW8 (SMARTvector particles) and analyzed for OFE (e) and SFE (f). Protein and mRNA levels are shown in Figure S4b,c, Supporting Information. In C-F, data are normalized to Ctrl in each experiment and expressed as mean ± SD. The number of biological replicates is shown (n=3). Statistical analysis was performed with the unpaired t-test, and significant differences are indicated by asterisks. In this and all subsequent figures, *, p <0.05, **, p < 0.01, *** p < 0.001.

### 2.4 *In vivo* tumorigenicity of NUMB-deficient BC cells is dependent on CRL7^FBXW^-mediated NUMB degradation

We investigated the relevance of CRL7^FBXW^–mediated NUMB degradation to tumorigenesis in BC using an integrated pharmacological and molecular genetics approach. Initially, we examined the effects of BTZ or MLN on the tumorigenicity of MDA-MB-361 or MDA-MB-231 cells. Xenografts were established by transplanting cells into the inguinal mammary fat pads of NOD/SCID/IL2Rγ^−/−^ (NSG) mice. Once tumors were palpable, BTZ and MLN were administered according to the scheme in Figure 5a. Both inhibitors significantly and comparably reduced the growth of MDA-MB-361 xenografts (Figure 5b), accompanied by NUMB restoration (Figure 5c). In contrast, no effects were observed in MDA-MB-231 xenografts (Figure 5b,c). The efficacy of the *in vivo* inhibitor treatments was confirmed by p21 restoration (BTZ, MLN) and the reduction in CUL1 NEDDylation (for MLN), in both model systems (Figure 5c). These *in vivo* results were paralleled in *in vitro* assays. Similarly to RBX1 and FBXW8 silencing (Figure 4c,d), BTZ and MLN treatment reduced both the OFE and SFE of MDA-MB-361 cells, while having no effects on MDA-MB-231 cells (Figure 5d). These results indicate that BTZ and MLN can selectively reduce the tumorigenic potential and size of the CSC compartment in Numb-deficient BC cells.

**Figure 5.**
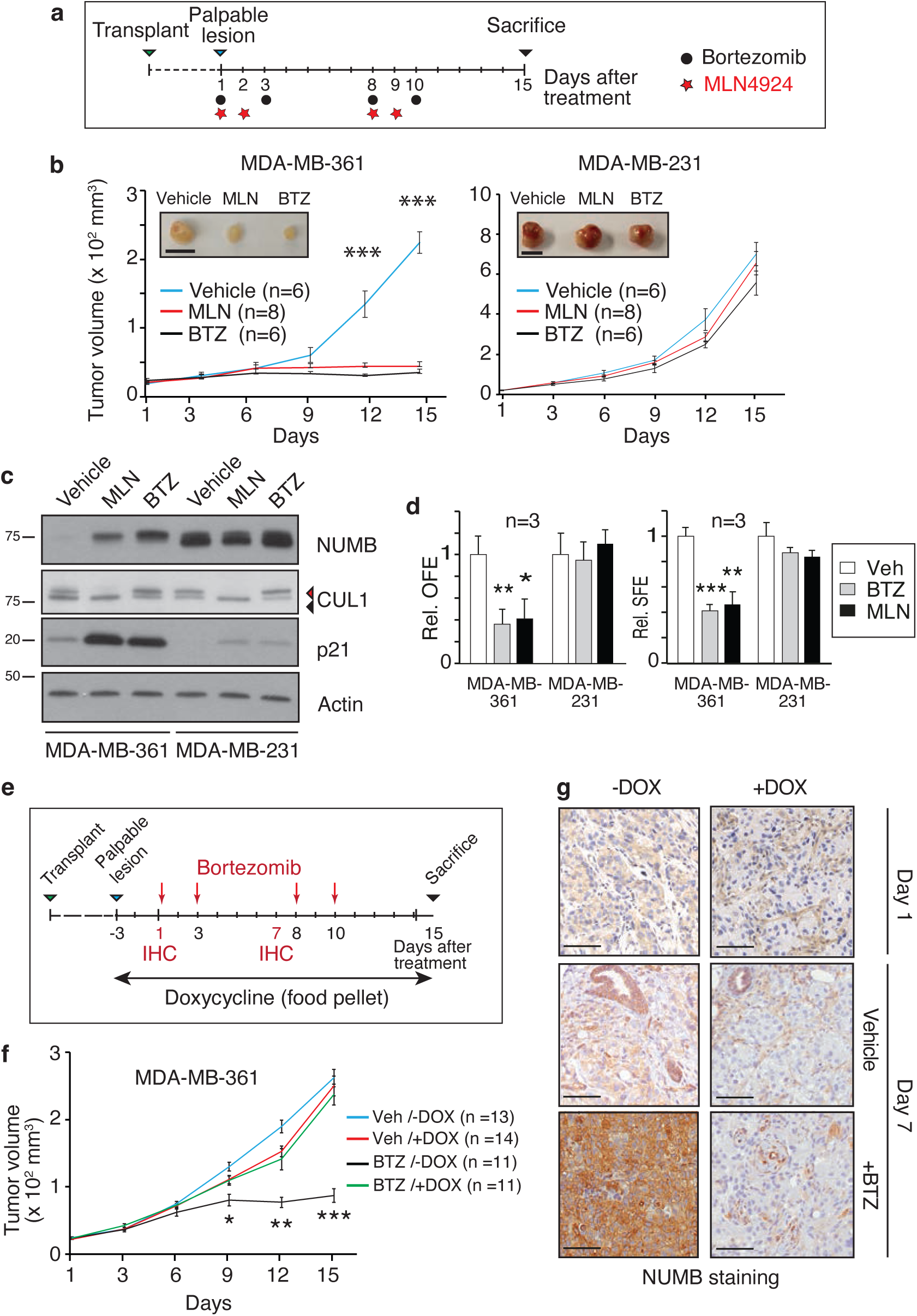
Tumorigenic potential of NUMB-deficient BC cells is dependent on CRL7^FBXW8^–mediated NUMB degradation. **a.** Scheme of the BTZ (black circles) and MLN (red stars) *in vivo* treatment protocol. Cells were transplanted into the inguinal mammary fat pads of NSG mice. When xenografts reached palpable size (∼20 mm^3^) drugs were administered as indicated. **b.** Growth curves of MDA-MB-361 or MDAMB-231 xenografts in NSG mice, mock-treated (Vehicle) or treated with BTZ (350 μg/Kg, once a day) or MLN (30 mg/Kg, twice daily). Data are expressed as mean ± SE. Statistical analysis was performed with the unpaired t-test and significant differences between treated *vs.* vehicle control are indicated; n, number of tumors per experimental group (in this and all subsequent figures). Inset images show typical final tumor sizes. Bar, 1 cm. **c.** Tumors, excised at the end of the experiments shown in ‘b’, were IB as indicated. For CUL1, the arrowheads point to NEDDylated (red) and non-NEDDylated (black) forms. **d.** Effects of *in vitro* BTZ (20 nM) and MLN (0.5 μM) treatments on the OFE (Matrigel) and SFE (methylcellulose) of MDA-MB-361 or MDA-MB-231 cells. Data are normalized to respective control and expressed as mean ± SD. Statistical analysis was performed with the unpaired t-test. **e.** Scheme of the BTZ and DOX treatment protocol used for xenografts of MDA-MB-361 cells stably expressing the DOX-inducible shRNA-NUMB construct. BTZ was administered as indicated by red arrows. DOX was administered via food pellets over the period indicated by the black arrow. Representative animals were sacrificed at days 1 and 7 to assess NUMB levels in xenografts by IHC. **f.** Growth curves of MDA-MB-361-inducible-shNUMB xenografts treated with BTZ or vehicle in the presence and absence of DOX. Data are expressed as mean ± SE. Statistical analysis was performed with unpaired t-test, and significant differences between BTZ/+DOX *vs*. BTZ/-DOX samples are indicated. **g.** Analysis of NUMB expression by IHC on FFPE sections of xenografts as in ‘f’, at days 1 and 7 of the experiment (see also panel e). Bar = 100 μm.

To assess the dependency of growth inhibition caused by BTZ on the restoration of NUMB expression, we used the above-described MDA-MB-361 cells stably expressing a DOX-inducible shRNA-NUMB construct (Figure 4e,f and Figure S4b,c, Supporting Information). These cells were transplanted into the inguinal mammary fat pads of NSG mice. When xenografts reached a palpable size, mice were treated with BTZ, with or without concomitant DOX treatment to induce Numb shRNA expression (see scheme in Figure 5e). NUMB KD *in vivo* completely abrogated the growth inhibitory effect of BTZ, arguing that the relevant target of proteasome inhibition *in vivo* was NUMB itself (Figure 5f). The efficacy of BTZ treatment and NUMB silencing in xenografts was confirmed by anti-NUMB IHC staining of FFPE sections (Figure 5g).

We next asked whether the tumorigenicity of MDA-MB-361 cells is specifically due to the exaggerated action of the CRL7^FBXW8^ complex on NUMB. To investigate this, we assessed the impact of RBX1 and FBXW8 silencing on xenograft growth, exploiting the above-described cell lines stably transduced with DOX-inducible vectors expressing the appropriate shRNAs. Xenografts were established from MDA-MB-361 and MDA-MB-231 cells transduced with inducible shRBX1, shFBXW8 or control shRNA. When xenografts reached palpable size, mice were administered DOX or treated with BTZ as a positive control for NUMB restoration (Figure 6a). The silencing of RBX1 or FBXW8 significantly impaired MDA-MB-361 xenograft growth, similarly to BTZ, but had no effect on MDA-MB-231 xenografts (Figure 6b-d). At the end of the experiment, tumors were excised and cellular lysates analyzed by IB to confirm RBX1/FBXW8 KD efficiency and NUMB restoration (Figure 6e). These results strongly indicate that the tumorigenic properties of MDA-MB-361 cells are dependent on the exaggerated activity of the CRL7^FBXW8^ complex towards NUMB.

**Figure 6.**
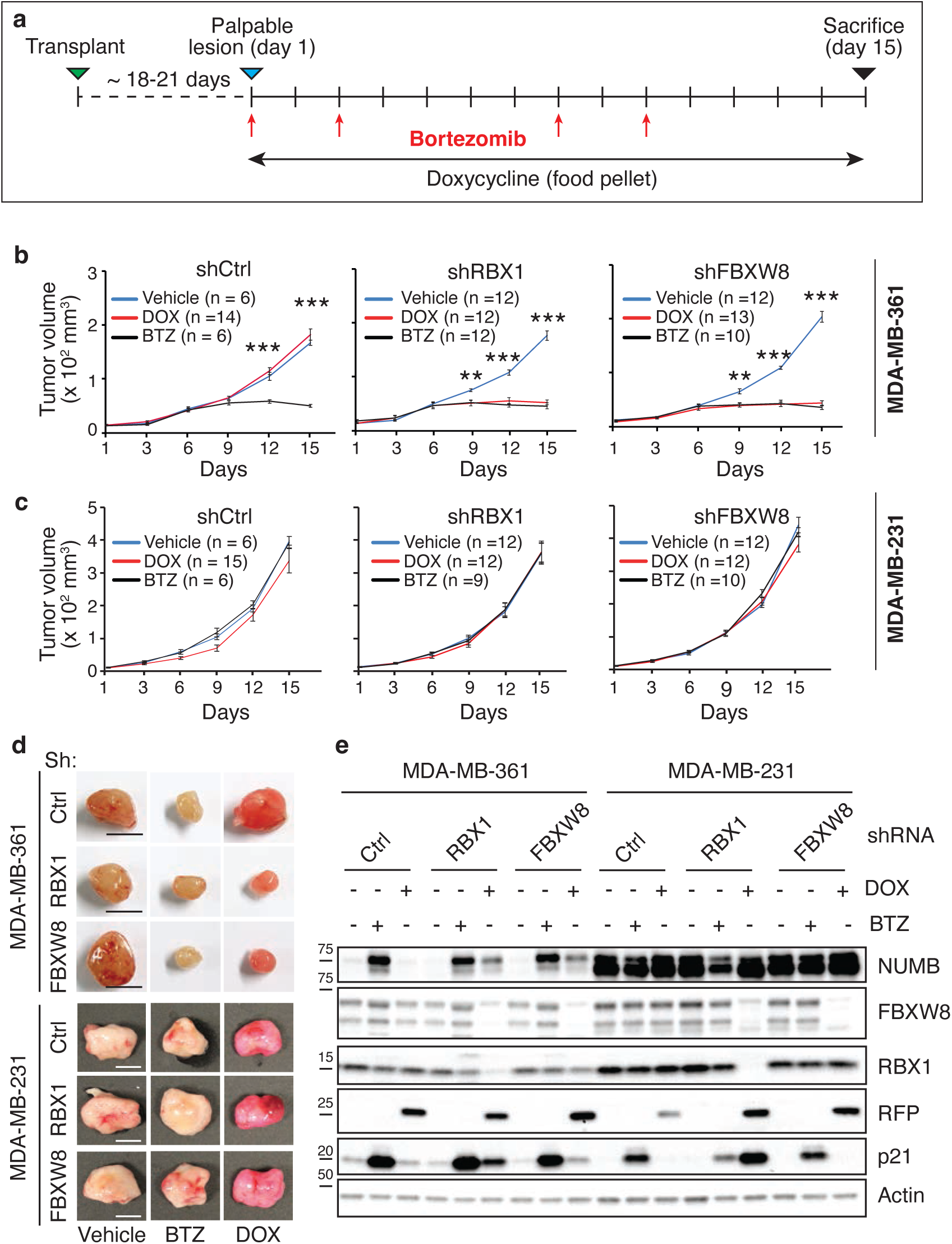
The CRL7^FBXW8^ complex is responsible for the tumorigenicity of NUMB-deficient BC cells. **a.** Scheme of the treatment protocol of xenografts derived from MDA-MB-361 or MDA-MB-231 cells transduced with DOX-inducible constructs expressing shRBX1, shFBXW8 or control shRNA (shCtrl). DOX was administered via food pellets over the indicated period. Two groups of xenografted mice were treated in parallel with BTZ or vehicle (red arrows) as controls. **b, c.** Growth of MDA-MB-361 (b) or MDAMB-MB-231 (c) xenografts treated as outlined in ‘a’. Data are expressed as mean ± SE. Statistical analysis was performed with unpaired t-test. Significant differences between Dox-/BTZ-treated *vs*. control samples are indicated. n, number of tumors per experimental group. **d.** Representative final xenograft sizes from experiments in ‘b’ and ‘c’. Bar, 1 cm. **E.** Xenografts excised at the end of the experiments in ‘b’ and ‘c’ were analyzed by IB as indicated. RFP is a reporter of DOX-induced shRNA expression used to assess the efficacy of DOX treatment.

### 2.5 CRL7^FBXW8^-mediated NUMB hyper-degradation is a therapeutic vulnerability in clinical BC samples

The data obtained in established BC cell lines suggest that CRL7^FBXW8^-mediated NUMB hyper-degradation could represent a driver alteration in clinical cases of NUMB-deficient BC characterized by low/undetectable levels of NUMB protein despite normal *NUMB* mRNA levels. To test this hypothesis, we established PDXs of BC specimens displaying different characteristics. These included a triple-negative and a luminal tumor with a NUMB-deficient status and normal *NUMB* mRNA levels (T1 and T2, respectively), and a luminal and a triple-negative tumor with a NUMB-proficient status at both the protein and mRNA levels (TA and TB, respectively) (Figure 7a-c).

**Figure 7.**
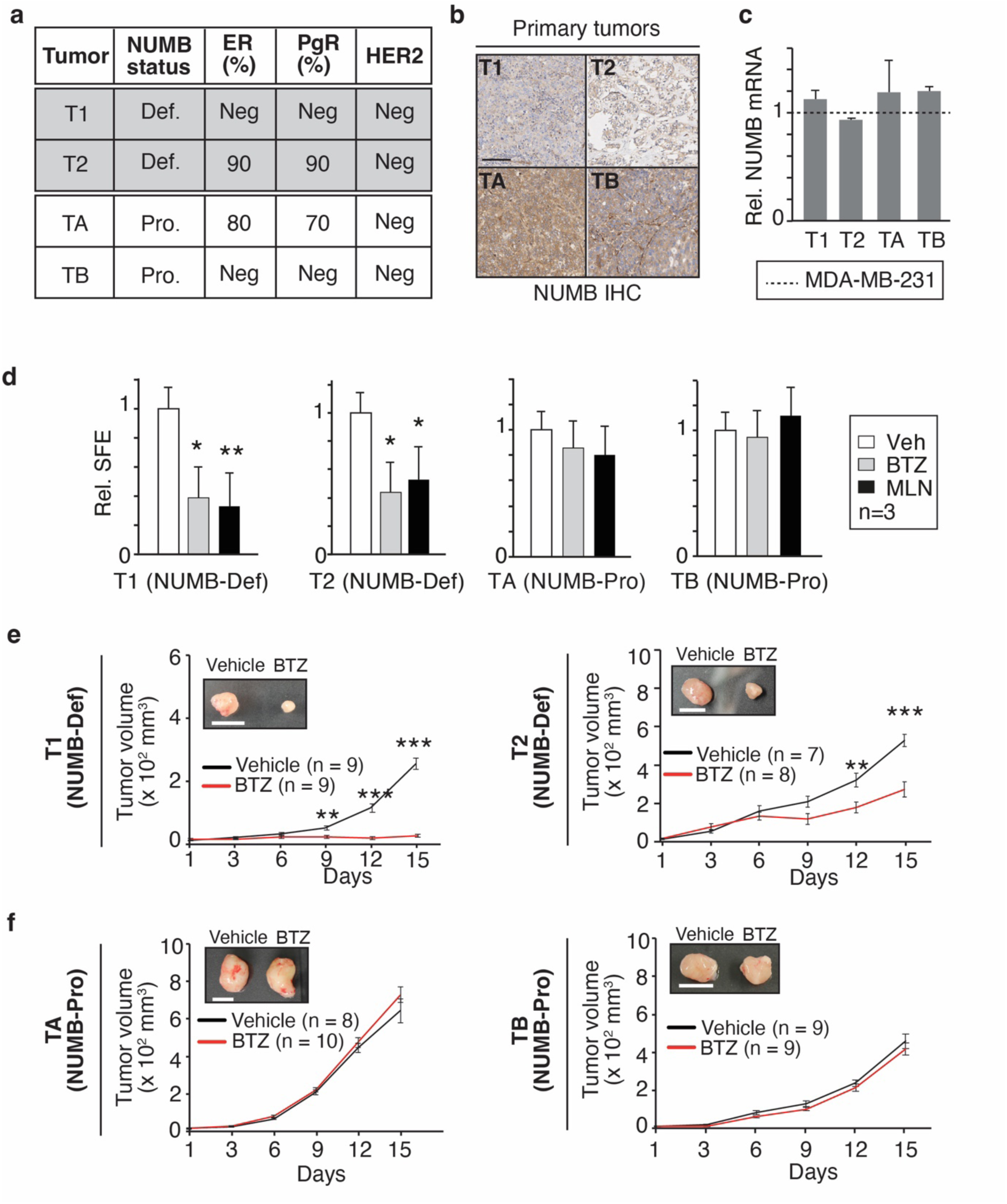
Targeting of the ubiquitin-proteasome system has anti-tumoral effects in NUMB-deficient PDXs. **a.** NUMB, hormone receptor (ER, estrogen receptor; PgR, progesterone receptor) and HER2 status in the selected PDXs is shown. Def, deficient; Pro, proficient, Neg, negative. **b.** IHC staining of NUMB expression in the primary tumors from which the PDXs were established. Bar, 100 μm. **c.** *NUMB* mRNA levels in the indicated PDXs relative to the reference cell line MDA-MB-231 (dashed line). Results are expressed as mean ± SD, normalized to MDA-MB-231 (2^^-ΔCt^ method). **d.** SFE in methylcellulose of PDX-derived primary cells treated or not at plating with BTZ (20 nM) or MLN (0.5 μM). Data are normalized to corresponding vehicle (Veh) control and expressed as mean ± SD (n=3). Statistical analysis was performed with the unpaired t-test. Significant differences are indicated. **e, f.** Growth of PDXs in NSG mice treated with BTZ or vehicle alone following the treatment protocol in Figure 5a. Data are expressed as mean ± SE. Statistical analysis was performed with unpaired t-test. Significant differences are indicated. The insets show typical final tumor sizes. Bars, 1 cm.

Initially, we tested the sensitivity of primary cells, isolated from the different PDXs, to BTZ and MLN in the *in vitro* 3D culture assays. NUMB-deficient primary cell lines (T1 and T2) displayed selective sensitivity to BTZ and MLN, evidenced by a reduction in SFE (Figure 7d) and OFE (Figure S5a, Supporting Information). These effects were mirrored *in vivo*, where the growth of NUMB-deficient BC xenografts was severely impaired by BTZ treatment (Figure 7e). In contrast, no effects of the treatments were observed on the NUMB-proficient primary cells *in vitro* or *in vivo* (Figure 7d,f and Figure S5a, Supporting Information). The restoration of NUMB following *in vivo* BTZ treatment was confirmed by IB and IHC analysis of the tumor outgrowths at the end of the experiment (Figure S5b,c, Supporting Information).

To provide further support for the idea that the effects of BTZ and MLN are mediated through the CRL7^FBXW8^ complex, we examined the effects of silencing RBX1 and FBXW8 on NUMB-deficient (T1 and T2) and NUMB-proficient (TA and TB) PDXs (Figure S5d,e, Supporting Information). Primary cells isolated from PDXs were stably transduced with the appropriate pTRIPZ DOX-inducible shRNA lentiviral vectors as described above for the established cell lines. *In vitro*, both RBX1 and FBXW8 KD resulted in a significant decrease in the SFE and OFE of NUMB-deficient T1 and T2 BC cells, while no effects were observed in NUMB-proficient TA and TB BC cells (Figure 8a,b). Of note, the effects of RBX1 and FBXW8 KD were not additive to the effects of BTZ or MLN on the SFE of NUMB-deficient (or NUMB-proficient) BCs, compatible with the idea that the biological effects of BTZ and MLN are exerted on the same pathway controlled by the CRL7^FBXW8^ complex (Figure S6, Supporting Information).

**Figure 8.**
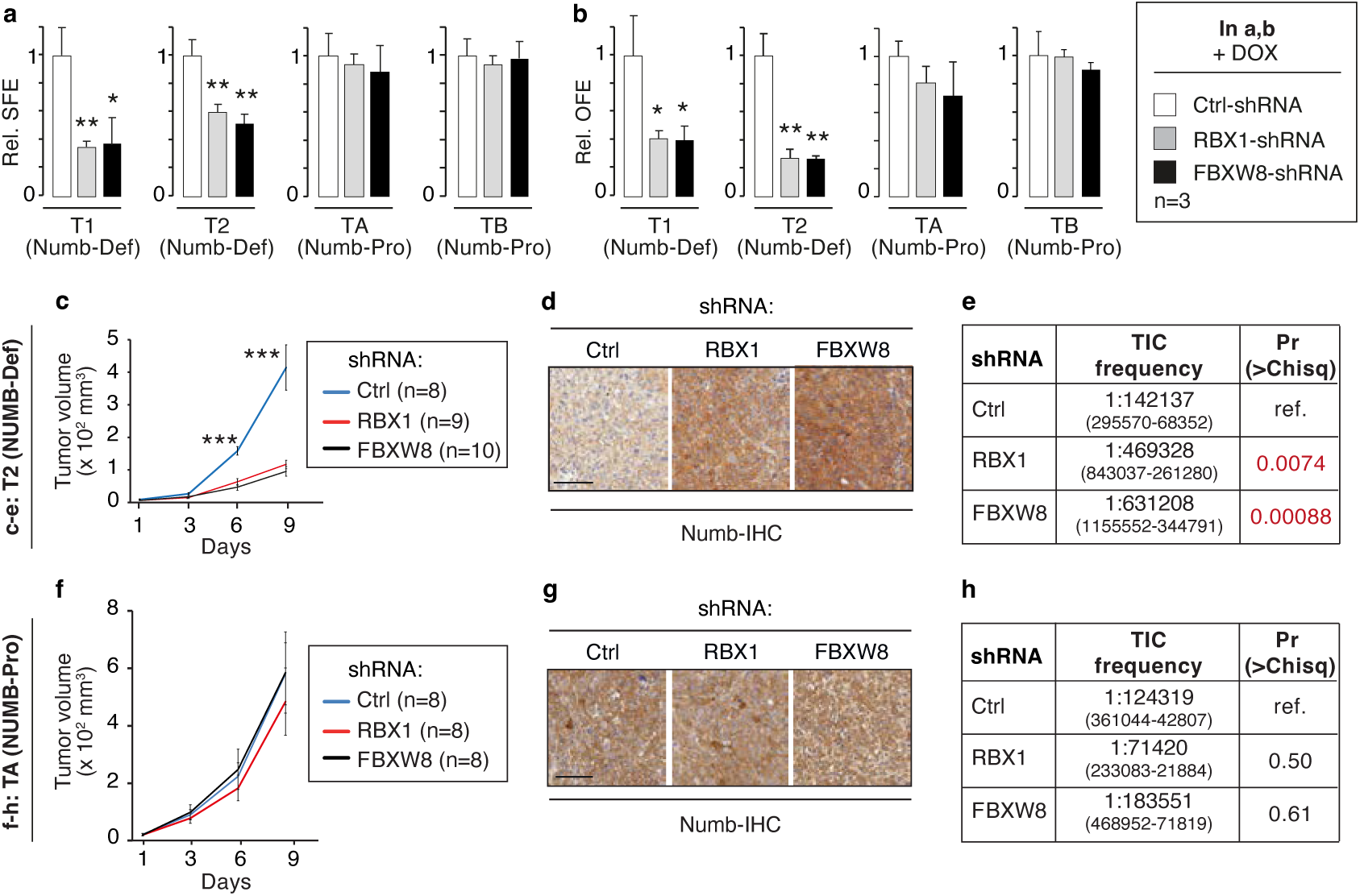
CRL7^FBXW8^-mediated NUMB degradation is essential for maintenance of the CSC compartment in clinical NUMB-deficient BCs. **a,b.** SFE (a) and OFE (b) of PDX-derived primary cells transduced with the indicated pTRIPZ lentiviral vectors encoding DOX-inducible shRNA constructs, in the presence of DOX. Data are normalized to the corresponding Ctrl-shRNA sample and expressed as mean ± SD (n=3). Statistical analysis was performed with the unpaired t-test. Significant differences are indicated. **c.** NUMB-deficient T2 BC primary cells transduced as in ‘a’ were transplanted in NSG mice. Once outgrowths were palpable, mice were administered DOX as shown in Figure 6a and tumor growth monitored. Data are expressed as mean ± SE. Statistical analysis was performed with the unpaired t-test. Significant differences between shRBX1 or shFBXW8 *vs*. shCtrl samples are indicated. **d.** Tumors at the end of the experiment in ‘c’ were stained by IHC to detect NUMB levels. Bar, 100 μm. **e.** NUMB-deficient T2 BC primary cells transduced as in ‘a’ were assessed for frequency of tumor-initiating cells (TICs) in a limiting-dilution xenograft assay. The ELDA software (https://bioinf.wehi.edu.au/software/elda/) was used for statistical analysis.^[32]^ The frequency of TICs, with 95% confidence intervals in parentheses, and the pairwise test for differences in SC frequency *vs*. control, are shown. **f-h**. The same set of experiments reported in c-e was replicated with the NUMB-proficient TA primary BC culture transduced as indicated in ‘a’. Bar in G, 100 μm.

Analogous results were obtained in *in vivo* tumorigenicity assays. Due to the limited availability of cells, we could only test the tumorigenic properties of one NUMB-deficient (T2) and one NUMB-proficient (TA) PDX in the different KD conditions. RBX1 or FBXW8 KD significantly inhibited the growth of NUMB-deficient T2 PDXs (Figure 8c) and efficiently restored NUMB expression as evidenced by IHC analysis of the final tumors (Figure 8d).

Finally, since NUMB LOF is needed to maintain the CSC compartment in NUMB-deficient BCs,^[1e, 1h, 1m]^ we investigated the effects of RBX1 and FBXW8 KD directly on this cell compartment *in vivo*. The frequency of tumor-initiating cells (TICs), which operationally define CSCs, was measured by the limiting dilution xenograft assay. Both RBX1 and FBXW8 KD reduced the number of TICs in T2 cells by 3-4-fold (Figure 8e). Conversely, NUMB-proficient TA cells were insensitive to RBX1/FBXW8 KD, both in the xenograft growth assay and the TIC assay (Figure 8f-h). Cells from the same tumors, were also analyzed for the expression of the SC and CSC marker ALDH.^[18]^ The silencing of RBX1 or FBXW8 significantly reduced the expression of ALDH in cells from the NUMB-deficient tumor, but not in cells from the NUMB-proficient one, further confirming that the degradation of NUMB, mediated by the CRL7^FBXW8^ complex, is needed for the maintenance of the CSC state (Figure S7, Supporting Information).

The sum of our results argues that the hyper-degradation of NUMB by the CRL7^FBXW8^ complex is causal in the natural history of the tumor and represents a therapeutic vulnerability, paving the way for the development of novel targeted therapies for BC.

## 3. Discussion

NUMB LOF in the mammary gland leads to tumorigenesis by subverting the physiological dynamics of the SC compartment and progenitor maturation. Under physiological conditions, the division of a mammary SC generates two daughter cells that adopt alternative fates: one retains the SC identity, withdrawing into quiescence yet retaining self-renewal ability; the other commits to a progenitor fate, undergoing several cycles of division followed by terminal differentiation.^[19]^ NUMB is central to this cell fate specification process, as it is asymmetrically partitioned between the two daughters, imposing a SC fate onto the daughter that inherits the majority of the protein.^[1m]^ Molecularly, this is partly due to the ability of NUMB to stabilize p53, through inhibition of MDM2, thereby determining quiescence, a hallmark of stemness,^[20]^ in the NUMB-inheriting cell.^[1a, 1b]^ In contrast, in the daughter cell that commits to progenitor fate, the initial loss of NUMB expression is progressively reversed in the transit-amplifying progenitor compartment, where the re-expression of NUMB is essential for correction differentiation.^[1m]^ NUMB LOF is therefore predicted to be catastrophic to the dynamics of the SC compartment, an issue compounded by the fact that NUMB also inhibits NOTCH, a potent proliferation-inducer in various cellular settings and developmental stages.^[3, 5a, 5c, 5d]^ Indeed, in the absence of NUMB, mammary SC division occurs predominantly symmetrically producing immature progenitors displaying phenotypic plasticity and self-renewal ability associated with EMT activation. This ultimately leads to the expansion of an immature progenitor/SC-like compartment functionally equivalent to CSCs.^[1m]^

In clinical BC samples, NUMB LOF, due to various molecular mechanisms, is a frequent event, occurring in the majority of BC cases (Figure 1e).^[1b, 1e, 1r]^ These mechanisms include aberrant splicing resulting in loss of exon 3-containing *NUMB* isoforms and subsequent p53 degradation.^[1b]^ Additionally, hyper-phosphorylation of NUMB by PKCs leads to loss of its asymmetric partitioning at SC mitosis and functional inactivation.^[1e]^ Finally, heightened NUMB ubiquitination leads to its excessive proteasomal degradation (^[1r]^ and this paper).

In this study, we investigated the mechanism underlying aberrant proteasomal degradation of NUMB in BC. By comparing NUMB-proficient and NUMB-deficient BC cell lines and clinical samples, we established that the CRL7^FBXW8^ E3 ligase complex is responsible for NUMB hyper-degradation. Silencing of the CRL7^FBXW8^ complex components restored NUMB levels in NUMB-deficient BC cells and reduced their tumorigenic potential both *in vitro* (SFE and OFE) and *in vivo* (xenograft growth and frequency of TICs). These data are consistent with the known role of NUMB LOF in the emergence and expansion of the CSC population and in BC tumorigenesis.^[1a, 1e, 1h, 1m, 1r]^ The role of the CRL7^FBXW8^ complex in NUMB-deficient BC was specifically due to its ability to degrade NUMB, since the effects of its silencing were reversed by the simultaneous ablation of NUMB.

Recent structural studies support a scenario where the CRL7^FBXW8^ complex functions as a substrate receptor for a NEDDylated CUL1-RBX1 catalytic module mediating ubiquitination in a multi-RING-E3-catalyzed reaction (Figure S1a, Supporting Information).^[10]^ A corollary of this model is that KD of either CUL7 or CUL1 should rescue the ubiquitination and degradation of target proteins. Our results largely agree with this model, in that the individual CUL7 or CUL1 KD increased NUMB levels in a NUMB-deficient cell line. The rescue, however, was only partial and the double CUL1-CUL7 KD was required to restore NUMB levels with an efficiency equivalent to RBX1/FBXW8 KD. Whether the partial rescue induced by the individual KDs was due to incomplete silencing or to more complex mechanistic explanations, relating to possible redundancy between CRL complexes, remains to be determined. Similarly, it remains to be established how NUMB interacts with the catalytic complex.

Previously, several E3 ligases, such as LNX1/2, MDM2, and SIAH1, have been linked to NUMB ubiquitination and degradation.^[15]^ Our results show that in NUMB-deficient BCs these enzymes do not appear to be involved in controlling NUMB levels, which instead are regulated by CRL7^FBXW8^. This raises the question of the molecular mechanism that directs NUMB to degradation via the CRL7^FBXW8^ complex in some BCs or, in other words, why is the CRL7^FBXW8^ complex active on NUMB in some BCs and not in others? One possibility is that components of the CRL7^FBXW8^ complex are altered genetically in some BCs. A screening of the METABRIC and TCGA databases,^[13, 21]^ however, revealed a very low frequency of mutations and/or amplifications of the *FBXW8, RBX1, SKP1, CUL1, CUL7* and *NUMB* genes in BC (Figure S8a, Supporting Information). Similarly, an extensive comparison of NUMB protein levels and mRNA levels of *FBXW8, RBX1, SKP1, CUL1* or *CUL7* in the IEO Cohort did not reveal any correlations worthy of note, arguing against overexpression as a probable cause (Figure S8b,c, Supporting Information).

Some characteristics of the modalities of target substrate recognition by the CRL7^FBXW8^ complex could offer viable working hypotheses to account for NUMB degradation. In contrast to other cullin-based complexes that ubiquitinate a wide array of substrates, CRL7^FBXW8^ appears to be much more selective. A handful of CRL7^FBXW8^ substrates are known, including IRS1, TBC1D3, Cyclin D1, HPK1, NANOG and OCT4.^[22]^ Interestingly, phosphorylation events seem to regulate the interaction of substrates with FBXW8 (or with the entire CRL7^FBXW8^ complex) and the ensuing substrate ubiquitination/degradation. In the case of IRS1, phosphorylation of both IRS1 and FBXW8 appear to be required for IRS1 degradation.^[22a-c]^ Phosphorylation of TBC1D3 is likely involved in its binding to FBXW8 and CRL7^FBXW8^-mediated degradation.^[22d]^ Similarly, Cyclin D1 phosphorylation is needed for its interaction with CRL7^FBXW8^ and degradation.^[22e]^ Autophosphorylation of the kinase HPK1 is required for its degradation by the CRL7^FBXW8^ complex.^[22f]^ Two transcription factors, involved in the homeostasis of embryonic SCs, NANOG and OCT4 (HUGO: POU5F1) have also been described as CRL7^FBXW8^ complex substrates, dependent on their phosphorylation which determines binding to FBXW8.^[22g, 22h]^

It is, therefore, tempting to speculate that aberrant phosphorylation events, affecting NUMB (or FBXW8) might be involved in CRL7^FBXW8^-dependent hyper-degradation of NUMB in BC. Notably, NUMB is a phosphoprotein whose activity is controlled by phosphorylation. Specifically, phosphorylation regulates NUMB asymmetrical partitioning at SC mitosis,^[1e, 23]^ and its ability to bind and regulate NOTCH and MDM2/p53.^[1e]^ In addition, NUMB phosphorylation disrupts interactions with β-catenin, p120 catenin and the clathrin adaptor α-adaptin.^[23d, 24]^ Three evolutionarily conserved serine phosphorylation sites in NUMB have been implicated in its phosphoregulation: Ser7, Ser276 and Ser295.^[23b, 23d, 23e, 24]^ These residues are phosphorylated by atypical PKCs, under physiological conditions or by aberrantly activated typical PKCs in BC.^[1e, 22e-h, 23a, 23c, 23e]^ However, these phosphosites do not appear to be involved in regulating the stability of NUMB.^[1e]^ Nevertheless, there are 25 Ser/Thr and one Tyr phosphosites in NUMB.^[25]^ Thus, the possibility that NUMB phosphorylation by an, as yet, unidentified kinase controls its degradation in some BCs, warrants further investigation.

Finally, the discovery that NUMB is degraded via CRL7^FBXW8^-mediated degradation has interesting therapeutic implications. Our previous work demonstrated, in pre-clinical models, that targeting NUMB/p53 LOF in NUMB-deficient BCs using the MDM2 inhibitor, Nutlin-3, to restore functional p53, is an efficient anti-CSC therapy.^[1m]^ However, despite intense research, clinical use of MDM2 inhibitors has been hampered by their limited clinical efficacy and associated toxicities.^[26]^ In this study, by elucidating the mechanism of aberrant NUMB degradation in BC, we have highlighted an additional point of therapeutic intervention in the NUMB/p53 pathway. Indeed, direct targeting of the enzymatic activity of the CRL7^FBXW8^ complex is a possibility. This can be achieved by NEDDylation inhibitors, such as MLN4924 (Pevonedistat),^[14d]^ which is in clinical trials for various malignancies. Another possibility is selective targeting of FBXW8. This adaptor protein exhibits selective interactions with CRL7^FBXW8^ substrates. Thus, its inhibition might prove efficacious in stabilizing these substrates, including NUMB, possibly with modest general toxicity. Strategies to design potential FBXW8 inhibitors, by structure-based drug design, have been reported.^[27]^ NUMB expression in BC might provide a useful stratification criterion for pre-clinical and eventually clinical studies with these compounds.

## 4. Experimental Section

### Clinical samples and clinical cohorts

Fresh, frozen, and archival FFPE normal and tumor mammary tissue specimens were collected at the European Institute of Oncology (IEO) following standard operating procedures approved by the Institutional Ethical Board (reference, UID 2931). Informed consent was obtained for all specimens linked to clinical data.

The IEO consecutive cohort of 2453 BC patients, who underwent surgery at IEO between 1997 – 2000 has been previously described.^[1b, 28]^ From this cohort, we randomly selected a sub-cohort of 672 patients (matched to the entire cohort), containing 585 patients free of distant metastasis and 87 patients with distant metastasis. To obtain the case-cohort used in this study (IEO-Cohort), we added the remaining patients who experienced distant metastasis within 10 yr (218 patients). The final case-cohort included 305 patients with distant metastasis and 585 patients who were free of distant metastasis at 10 yr follow-up. The primary endpoint of the analyses was death related to BC (DRBC).

We also employed two publicly available datasets. The METABRIC dataset (1904 samples) was obtained through the cBioPortal (2019 freeze, available at https://github.com/cBioPortal/datahub/tree/master/public/brca_metabric).^[13]^ The TCGA BC dataset was downloaded from the cBioPortal (http://www.cbioportal.org/).^[21]^

### Analyses on clinical cohorts

NUMB status was attributed to IEO cohort cases (or to BCs used in Figures 7, 8 and Figures S1b, S5, Supporting Information) by measuring the NUMB protein levels by IHC on whole FFPE slices (see Supporting Methods, Supporting Information). Normal mammary tissues displayed homogenous and intense NUMB staining in the luminal layer (IHC score 2-3).^[1a, 1r]^ Tumors were classified on a NUMB IHC scale from 0 to 3; 0 = undetectable expression; score ≥ 2, expression comparable to that of normal luminal mammary cells (see examples in Figure S1b, Supporting Information). Tumors were classified as NUMB-proficient if they exhibited staining comparable to normal breast (IHC score ≥ 2.0 in at least 70% of the cells), all other cases were labeled as NUMB-deficient.

Transcriptomic profiling of the IEO cohort was performed by RNAseq (see Supporting Methods, Supporting Information), and relevant data for genes-of-interest were extracted.

All Kaplan-Meier, univariate and multivariable survival analyses, were conducted using the JMP software version 14.3 (SAS Institute Inc., Cary, NC, 1989–2023). The Cox proportional hazards model was used to calculate the Hazard Ratio and corresponding p-values.

### Animal experiments and treatments with drugs

All *in vivo* experiments were approved by the Italian Ministry of Health (Protocol number 704/2018-PR) and performed in accordance with the Italian laws (D.L.vo 116/92 and following additions), which enforce the EU 86/609 directive, and under the control of the institutional organism for animal welfare and ethical approach to animals in experimental procedures (Cogentech OPBA). Mice were maintained in a controlled environment, at 18–23 °C, 40–60% humidity and with 12-h dark/12-h light cycles, in a certified animal facility. Animal weight was monitored throughout the experiments and variations never exceeded 20%; animals did not show evident signs of distress throughout the experiment. Potential neuro-toxicity was also evaluated (appearance of ataxia, numbness and loss of four limb grasping strength), with no noticeable signs of toxicity. At sacrifice, macroscopic examination of internal organs did not show evident signs of toxicity. The size of the animal cohorts in each group was empirically established, based on our previous experience, with the aim of minimizing the number of animals used.^[1e, 1h]^

The concentrations of BTZ and MLN4924 used in the animal experiments were established based on extant literature on preclinical studies with the two drugs.^[14d, 29]^ For the *in vitro* treatments with the drugs, we used a concentration of 20 nM and 0.5 μM, for BTZ and MLN4924, respectively. For BTZ, it was shown that, at the approved therapeutic dose in humans, the drug reaches a blood concentration of ∼ 50 nM.^[30]^ We de-escalated this dose to 20 nM, as the lowest dose at which clear effects on NUMB levels in MDA-MB-361 cells were visible (see Figure 2a) in the absence of toxic effects. By adopting a similar criterion, we used a dose of and 0.5 μM for MLN4924, de-escalating from a typical dose of 2-3 μM used in literature (see for instance ^[14d]^).

### Gene silencing by lentiviral transduction and transient transfection

DOX-inducible shRNA pTRIPZ lentiviral vectors (Dharmacon) or SMARTvector inducible shRNA lentiviral particles (Dharmacon) were used for stable silencing of NUMB, RBX1, FBXW8, MDM2, LNX1, LNX2 or SIAH1. pTRIPZ shRNA lentiviral vectors were: RBX1 (Clone V3THS_337317), FBXW8 (Clone V3THS_364865), NUMB (Clone V3THS_397259), MDM2 (Clone V3THS_379472), LNX1 (Clone V2THS_159554), LNX2 (Clone V3THS_393564), SIAH1 (Clone V3THS_391107), (Figures 4, 5f,g, 6, 8 and Figures S4, S5d,e, Supporting Information). The pTRIPZ empty vector served as a negative control. VSV-G pseudotyped lentiviral vector stocks were produced in HEK293T cells by transient transfection of target plasmid and packaging plasmids, DR8.2 and VSV-G (10, 5 and 4 μg, respectively, for 10 cm dishes). Established and primary cell lines were stably transduced with the pTRIPZ lentiviral vectors in the presence of 8 μg/mL polybrene (Sigma). After 48 h, the infected cells were placed in fresh medium with 1 μg/mL puromycin (Vinci Bio Chem) and selected for 72 h. Finally, stable transfectants were treated *in vitro* with 1 μg/mL doxycycline-hyclate (DOX, Sigma) for at least 50 h to test the efficiency of shRNA expression and ensuing KD.

For the double NUMB/RBX or NUMB/FBXW8 KD experiments (Figure 4e,f and Figure S4b,c, Supporting Information), MDA-MB-361 cells stably transduced with the pTRIPZ lentiviral vector expressing DOX-inducible shNUMB, were further transduced with SMARTvector inducible lentiviral particles (Dharmacon) expressing shRBX1 (Clone V3IHSMCG_7732361) or shFBXW8 (Clone V3IHSMCG_5355635). Double transduced MDA-MB-361 cells were isolated by FACS-sorting of cells expressing both RFP and GFP, fluorescent reporter genes of pTRIPZ vectors and SMARTvectors, respectively. Briefly, double transduced cells were treated with DOX (1 μg/mL, 12 h) to induce pTRIPZ shRNA expression. Single cell suspensions were FACS sorted using a BD FACSMelody equipped with 2 laser lines (*i.e.*, 488 and 561 nm) and band-pass optical filters (*i.e.*, 527/32 and 582/15, respectively). Not-induced cells were used as a negative control to assess the background fluorescence. A drop frequency of 34 kHz, a sorting pressure of 22 PSI and a 100 μm nozzle were maintained. A mean sorting rate of 1000-1500 events/s was chosen.

To achieve transient KD (Figure 2d,e, Figure S7, Supporting Information), MDA-MB-361, MDA-MB-231 and PDX-derived primary cells were transfected with 5 nM siRNA oligo using Lipofectamine RNAiMAX (Invitrogen) cationic reagent following the “reverse” approach, as per the manufacturer’s instructions. Since the various cells differ in growth rate, the optimal number of seeded cells was 3.2×10^4^ for MDA-MB-361 and PDX-derived primary cells, and 8×10^3^ cells/cm^2^ for MDA-MB-231. After 72 h, cells were harvested and processed for IB, or for ALDH assays. In each experiment, Horizon siGENOME non-targeting control siRNA (D-001210-03) was included as negative control. Targeting siGENOME siRNA oligo duplexes were from Horizon and directed against: RBX1 (D-004087-1 and D-004087-5, corresponding to siRBX1 #1 and #2, respectively), FBXW8 (D-012431-3 and D-012431-4, corresponding to siFBXW8 #1 and #2, respectively), CUL1 (D-004086-01-0002), CUL4A (D-012610-01-005), CUL7 (D-017673-01-0002), SKP1 (D-003323-05-0002).

### Establishment of PDXs and primary cell cultures from clinical samples

To generate PDXs, fresh specimens from NUMB-deficient and NUMB-proficient human BCs were cut into 4 × 2 mm pieces using a razor blade, removing any necrotic tissue. The resulting fragments were embedded in ice-cold Matrigel (BD Matrigel™, BD Biosciences) and immediately injected bilaterally into the 4^th^ inguinal mammary glands of 8-week-old female NSG mice anesthetized by intraperitoneal injection of 150 mg/kg tribromoethanol (Avertin).^[31]^ Animals were euthanized when the tumor exceeded 10% body mass in compliance with regulations for use of vertebrate animals in research. Tumor volume was determined by measuring the tumor in two dimensions with calipers and calculated using the mathematical formula for a prolate ellipsoid: tumor volume = (L×W^2^)/2, where L = longest diameter; W = shortest diameter. Mice were monitored twice weekly by an investigator blinded to treatment assignment. Only animals that died during the experiments were excluded from the analysis (see Table S2, Supporting Information) (see also Supporting Methods, Supporting Information).

For the analysis of primary human BC cells derived from Numb-deficient (T1, T2) and NUMB-proficient (TA, TB) PDXs (Figure 7c,d, 8a,b, Figure S5a,b, Supporting Information), cells were isolated as follows. Tumor masses were mechanically minced and enzymatically digested in DMEM/F12 medium (Gibco) supplemented with 2 mM L-glutamine (Lonza), 2% BSA Fraction V (Sigma), 10 mM HEPES (Sigma), 100 U/mL hyaluronidase type IV (Sigma) and 200 U/mL collagenase type I (Sigma) at 37 °C for 4 h in a rotating wheel. Cell suspensions were centrifuged at 80 *g* for 5 min and red blood cells were lysed (ACK buffer, Lonza) to enrich the epithelial cell subpopulation. The cell suspension was sequentially passed through filters with decreasing pore size (100, 70, 40 μm) and the resulting flow-through was resuspended in Stem Cell Medium containing MEBM (Lonza), 2 mM L-glutamine (Lonza), 5 μg/mL human insulin (Sigma), 0.5 μg/mL hydrocortisone (Sigma), 20 ng/mL hEGF (Peprotech), 20 ng/mL hFGF (Peprotech), 1 U.I./mL heparin (Sigma), 2 % B-27 supplement (Gibco). Cell viability was assessed by trypan blue exclusion. Finally, cells were cultivated and expanded in nonadherent conditions in 6-well plates coated with poly-HEMA (Sigma) solution (1.2% in 95% Ethanol) in Stem Cell medium and a 5% CO_2_ humidified incubator at 37°C.

### *In vivo* tumorigenicity assays

Orthotopic xenografts of MDA-MB-361 and MDA-MB-231 cell lines were generated by injecting 25 μL of ice-cold 1:1 DPBS:Matrigel phenol red-free (Corning) single-cell suspension bilaterally into the 4^th^ inguinal mammary glands of 8-week-old female mice. To ensure synchronous onset of MDA-MB-361 and MDA-MB-231 tumors, different numbers of cells were transplanted according to their different proliferative rates: 2×10^6^ and 2.5×10^5^ cells/fat-pad, respectively. Xenografts of NUMB-deficient (T1, T2) and NUMB-proficient (TA, TB) primary BC cells were generated as described above.

For *in vivo* drug treatments, when xenografts reached palpable size (∼20 mm^3^), age-matched mice were assigned randomly to the various treatment or control (vehicle) branches. BTZ was administered intraperitoneally at 350 μg/Kg following the schedules shown in Figure 5a,e, and Figure 6a. MLN was dosed intraperitoneally 30 mg/Kg *bis in die* following the schedule shown in Figure 5a. Saline and 10% hydroxypropyl-β-cyclodextrin (HPBC, Sigma) were used as vehicle controls for BTZ and MLN, respectively. DOX was administered continuously by food pellets (Mucedola, 625 mg/Kg). *Ad libitum* ordinary food served as a negative control. Tumor growth was monitored over a 2-week period using Vernier calipers, after which animals were sacrificed and tumors excised.

For *in vivo* limiting dilution assays (Figure 8e,h), cells dissociated from NUMB-deficient (T2) and NUMB-proficient (TA) PDXs, were stably transduced with the appropriate pTRIPZ DOX-inducible shRNA lentiviral vectors as described above and treated *in vitro* with DOX (1 μg/mL, 72 h) to induce shRNA expression. Decreasing numbers of cells (1×10^6^-5×10^5^ -1×10^5^ cells/fat-pad) were transplanted into the abdominal mammary gland of >8-week-old female NSG mice. Transplantation frequency was calculated using ELDA software (https://bioinf.wehi.edu.au/software/elda/),^[32]^ as previously described.^[33]^

### *In vitro* tumorigenicity assays in 3D culture conditions

The colony (spheroid)-formation assay in methylcellulose was performed as previously described.^[1e]^ Briefly, single cell suspensions (1000-5000 cells/ml) were plated in 24-well plates in non-adherent conditions in Stem Cell medium containing 1% methylcellulose (Sigma) and incubated (humidified 5% CO_2_, 37°C). Treatments (20 nM BTZ, 0.5 μM MLN4924) were administered at plating. Colonies were manually counted after 7-15 days. The %SFE (number of spheres/number of plated cells x 100) was then calculated.

The 3D-Matrigel organoid assay was performed as previously described.^[34]^ Briefly, single cell suspensions were resuspended in 800 μL/well of complete medium containing 2% Matrigel, seeded in coated four-chamber slides (Nunc Lab-Tek II Chamber slide^TM^, Thermo Fisher Scientific, for Figures 2c and S4b, Supporting Information) or in a drop of pure/undiluted Matrigel (50 μl), and incubated (humidified 5% CO_2_, 37°C). Complete medium was changed every 4 days. The %OFE (number of organoids/number of plated cells x 100) was then calculated. When indicated, organoids were treated with 20 nM BTZ, 0.5 μM MLN4924, 1 μg/mL DOX or vehicle, once they reached an average size of ∼40 μm. For experiments in Figures 2c and 4b, organoids were recovered from Matrigel with a non-enzymatic procedure using Cell Recovery Solution (Corning) according to manufacturer’s instructions and then processed for further analysis.

### Statistical analyses

Statistical analyses and number of replicates for each experiment are indicated in the figures. In all figures: *, p <0.05, **, p < 0.01, *** p < 0.001. When p-values are reported in extenso, significant values are highlighted in red.

Additional methods (siRNA-based screening and NUMB capture ELISA assay, 2D culture of established BC cell lines, immunoblotting and immunoprecipitation, *in vivo* ubiquitination assays; immunofluorescence staining of 3D Matrigel organoids, histological and IHC analyses; total mRNA extraction, ALDH assays, and RT-qPCR; RNAseq) are in Supporting Methods, Supporting Information. A list of the antibodies used and their working dilutions is in Table S3, Supporting Information.

## Supporting information

Merged Supplementary Information

## Supporting Information

Supporting Information is available from the Wiley Online Library.

## Acknowledgements

We thank the IEO Technological Units, in particular the Biobank, the Molecular Pathology Unit, the Imaging Unit, the Flow Cytometry Unit, and the Genomics Unit. We also thank the Cogentech Real Time Quantitative PCR service.

This work was partially supported by core institutional funding through the Italian Ministry of Health Ricerca Corrente and 5×1000 funds, and by the following grants: Fondazione-AIRC (grant # IG 15538 and IG 23049 to SP, 23060 to PPDF, and 5x1000 MCO 10.000 to PPDF and SP); by the Italian Ministry of University and Scientific Research (PRIN2017 and PRIN2020 to SP; PRIN 2020 to PPDF; PNRR, project code CN00000041); by the Italian Ministry of Health (Ricerca Finalizzata RF-2016-02361540 and RF-2021-12373957 to PPDF; PNRR, project code PNRR-MCNT2-2023-12378490).

## Contributions

Conceptualization: PPDF, DT, SP

Investigation and data analysis: SS, MGF, LA, SCo, RB, SCa, INC, SF, GB, GF, EZ,

Funding acquisition: PPDF, SP

Supervision: PPDF, DT, SP, CM, MV

Writing - original draft: PPDF, DT, RHG

Writing - review & editing: All authors

SS and MGF contributed equally.

PPDF, DT and SP are equal last authors.

