## Supplementary material for "The CRL7^FBXW8^ Complex Controls the Mammary Stem Cell Compartment Through Regulation of NUMB Levels": Merged Supplementary Information

### SUPPORTING INFORMATION

### SUPPORTING INFORMATION

### SUPPORTING METHODS

#### siRNA-based screening and NUMB capture ELISA assay

A high-throughput siRNA-based screening coupled with a NUMB capture ELISA assay was performed to identify candidate proteins of the ubiquitination machinery potentially involved in the regulation of NUMB degradation. The screening was automated using the EVOware platform. The Dharmacon siGENOME® SMARTpool® siRNA Libraries, Human Ubiquitin Conjugation Subset 1-3 (Horizon) were used comprising ~ 600 siRNAs (4 siRNAs per target) targeting human Cullin, E1, E2 and HECT E3 ubiquitin ligases (Subset 1), F-box and SOCS box E3 ubiquitin ligases (Subset 2), RING finger and RING-finger-like E3 ligases (Subset 3) (see Table S1, Supporting Information). Transfection was performed with the reverse method. In brief, siRNA pools were spotted in 384-well plates (50 nM/well) in quadruplicate in a volume of 5 µL. Then, 5 µL of DharmaFECT 1 transfection reagent (Dharmacon), diluted 1:8 in DharmaFECT Cell Culture Reagent (Dharmacon), was dispensed into each well. The plates were incubated at RT for 20 min to allow formation of transfection complexes. Subsequently, MDA-MB-361 cells were plated at 6 x $10^3$ cells/well on the top of DharmaFECT 1 transfection reagent complexes. Plates were then incubated for 72 h at 37 °C.

As controls, we included wells with control siRNA, a siRNA targeting NUMB, MDA-MB-361 cells treated with MG-132 (5 µM for 24 h) and wells with no cells. We also included different amounts of recombinant NUMB to create a standard curve. Cell viability was controlled using the CellTiter-Fluor™ Cell Viability Assay (Promega).

The effects of RNAi on NUMB protein levels were assessed using a commercially available capture ELISA assay (DuoSet IC Human/Mouse/Rat Total NUMB, R&D Systems), which we modified to improve specificity and sensitivity. Cells were lysed, and lysates transferred to ELISA plates that had been coated with capture Ab, washed and blocked. We used the in-house mouse monoclonal anti-NUMB Ab (4 µg/ml) as the capture Ab (see Figure S2a, Supporting Information). For detection, we used the biotinylated anti-NUMB mouse polyclonal Ab and streptavidin-HRP system provided by the kit (100 ng/ml), and Supersignal ELISA Pico Luminol Chemiluminescence substrate (Thermo Scientific), instead of the TMB substrate provided by the kit.

Positive hits were identified from two independent biological replicates of the screening, as illustrated in the legend to Figure 2b, Supporting Information.

#### 2D culture of established BC cell lines

MDA-MB-361 and MDA-MB-231 were cultured in DMEM (Lonza), 2 mM L-glutamine and 10% FBS (North America origin, Hyclone). HEK293 and HEK293T cells were grown in DMEM, 2 mM L-glutamine and 10% FBS (South America origin, Microgem). Cells were incubated in a 5% $CO_2$ humidified incubator at 37°C. To prevent phenotypic and genetic drift, cell lines were always passaged in log phase with 0.05% trypsin and kept in culture for a maximum of 1 month, at which point a new batch of cells from the original stock was thawed. Mycoplasma tests were routinely performed on cultured cell lines.

#### Immunoblotting and immunoprecipitation

For the IB analysis of cell lines, cells were harvested, pelleted, washed twice with ice-cold DPBS and lysed in ice-cold RIPA buffer (50 mM TRIS pH 7.5, 150 mM NaCl, 0.5 mM EDTA pH

8, 1% NP-40, 0.5% Na-deoxycholate, 0.1% SDS) supplemented with Protease Inhibitor Cocktail III (Calbiochem) and PhosSTOP (Roche) for 30 min on ice. Samples were centrifuged (16000 *g,* 20 min, 4°C) and quantified with BCA protein assay (Pierce). Lysates were resuspended in 4X Laemmli loading buffer (250 mM TRIS pH 6.8, 40% glycerol, 8% SDS, 0.02% bromophenol blue, 50 mM DTT), boiled for 5 min at 95°C, and loaded onto SDS-PAGE gels. Proteins were transferred to nitrocellulose membranes. When necessary, identical samples were loaded onto different gels in parallel, to allow the detection of proteins with similar molecular weights. Membranes were also cut into horizontal strips to allow the detection of multiple proteins (see Table S3, Supporting Information for Ab details).

For the IB analysis of xenograft-derived cells (Figures 5c, 6e, and Figure S5b, Supporting Information), tumor masses were digested as described above, before following the IB protocol.

For the co-immunoprecipitation experiment (Figure 2f), HEK293 cells were transiently transfected, by the calcium phosphate method, with a pDNA 3.1 vector encoding FLAG-tagged human NUMB.[1] Briefly, 3 x $10^6$ cells were plated in 10 cm-dishes the day before transfection. A transfection mixture (15 µg DNA, 61 µL $CaCl_2$, 439 µL $H_2O$, 500 µL HBSS and 20 µM chloroquine) was left for 5 min at RT before adding to cells. Cells were then incubated for 6 h in a $CO_2$ incubator at 37°C before growth medium was replaced with fresh complete medium. Cells were harvested after 48 h and lysed in ice-cold JS buffer (50 mM HEPES pH 7.5, 150 mM NaCl, 5 mM EGTA, 1.5 mM $MgCl_2$, 10% glycerol, 1% Triton X-100) supplemented with protease and phosphatase inhibitors. Lysates were incubated with anti-FLAG M2 agarose beads (Sigma) at 4 °C for 2 h with rotation. To control for non-specific binding, 100 µg/mL FLAG peptide (Sigma) was used to compete with cellular proteins. As an additional negative control, an anti-HA (in-house produced mouse monoclonal Ab) immunoprecipitation was performed in parallel. Immunoprecipitates were washed 4 times with lysis buffer, eluted in 2X Laemmli loading buffer, boiled for 7 min at 95°C, centrifuged for 1 min and detected by IB.

#### *In vivo* ubiquitination assay

Ubiquitination assays were performed to evaluate the ubiquitination of NUMB in intact cells. Briefly, cells were cultured to 80% confluence and treated with MLN (0.5 µM) for 24 h (Figure 3a) or transiently silenced for RBX1 or FBXW8 (Figure 3b). Cells were then harvested, washed in ice-cold PBS, and lysed in RIPA buffer supplemented with protease, phosphatase and deubiquitinase inhibitors (PR619, 25 µM) on ice for 30 min. Cell lysates were then clarified by centrifugation at 14,000 × g for 10 min at 4°C. NUMB was immunoprecipitated from 6 mg of cell lysate from MDA-MD-361 and 3 mg from MDA-MD-231, respectively, for 2 h at 4°C using 0.8 µg/mg of anti-NUMB (Cell Signaling Technology) or 0.8 µg/mg Ctr-IgG (Santa Cruz). The immune complexes were washed five times, followed by SDS-PAGE and detection in IB with anti-ubiquitin (Santa Cruz) or anti-NUMB antibodies (Cell Signaling).

#### Immunofluorescence staining of 3D Matrigel organoids

3D organoids retrieved from Matrigel were spotted on polylysine coated coverslips (previously incubated with 15 µg/mL poly-D-lysine, Sigma, for 30 min at 37 °C) and then fixed with 4% paraformaldehyde (Themo Scientific) for 1 h. Coverslips were washed twice with 100 mM glycine in PBS to quench aldehyde-dependent auto-fluorescence. Organoids were permeabilized with 0.2% Triton X-100 in PBS for 1 h and non-specific hybridization was prevented by incubating samples with blocking buffer (10% normal donkey serum, 10% BSA, 0.1% Triton X-100 and 0.05% Tween-20 in PBS) for 3 h at RT. Appropriate primary and secondary Abs diluted in blocking buffer were incubated overnight at 4 °C. Nuclei were stained with DAPI for 20 min at RT. After each incubation, two extensive washes with 0.05% Tween-20 in PBS were performed. Finally, coverslips were dipped in dd$H_2O$ and mounted in glycerol mounting medium supplemented with DABCO antifade. Nail polish was used to anchor the coverslip to glass slide (see Table S3, Supporting Information for Ab details). Fluorescent dye signals were acquired sequentially using a Leica SP8 confocal microscope. To limit crosstalk between different channels a Leica HC PL APO 40X/1.13 oil immersion objective was used. Leica LasX and ImageJ software were used for image processing and analysis.

**Histological and IHC analyses**

FFPE samples were prepared from fresh tumor xenografts (Figure 5g, 7b, 8d, 8g, Figure S5c, Supporting Information). Samples were cut into pieces (< 5 mm diameter), placed in embedding cassettes, and fixed in 10% neutral buffered formalin at 4 °C. After ~18 h, samples were moved to cold 70% ethanol and were either stored for up to 7 days at 4 °C or immediately embedded in paraffin. For the latter, tissues were dehydrated through a graded alcohol series (70%, 95%, 100%) before embedding in paraffin. The FFPE specimens were sectioned (3 µm thick) and mounted onto positively charged glass slides (Menzel Glazer SuperFrost PLUS, Themo Scientific). Glass slides were deparaffinized in Bioclear, re-hydrated through a graded alcohol series (100%, 95%, 70%, and distilled $H_2O$) and routinely processed for H&E or IHC staining.

For NUMB IHC, samples were placed in epitope retrieval solution (1 mM EDTA pH 8, 0.05% Tween-20) for 50 min at 95°C. Endogenous peroxidase activity was then quenched with 3% $H_2O_2$. Non-specific Ab binding was reduced by incubation in blocking buffer (2% goat serum, 2% BSA, 0.05% Tween-20 in TBS), before adding primary anti-NUMB Ab (see Table S3, Supporting Information), diluted in blocking buffer, for 1 h at RT. The appropriate EnVision Plus Detection System (Dako), based on HRP-labeled polymer conjugated to secondary Ab, was used to detect the primary Ab, according to the manufacturer's instructions. DAB Plus Substrate Chromogen System (Dako) allowed visualization of bound Ab. Gill's hematoxylin (2 mg/100 mL) was added for 30 sec to counterstain nuclei. Before mounting slides with Eukitt medium (Sigma), samples were passed through a graded alcohol series (95%, 100%), 1 min each concentration, and three times in Bioclear (Bio-OPTICA) for 1 min. Representative images, from at least two sections, were acquired with Aperio AT2 slide scanner and processed with Aperio ImageScope software (Leica).

**ALDEFLUOR assay and flow cytometry**

To assess aldehyde dehydrogenase (ALDH) activity, the ALDEFLUOR™ assay kit (StemCell Technologies, Vancouver, Canada) was utilized according to the manufacturer's protocol. Briefly, $1 \times 10^6$ cells were collected and resuspended in 1 ml of ALDEFLUOR™ assay buffer. After addition of 5 µL ALDEFLUOR™ substrate (BODIPY-aminoacetaldehyde), 500 µL of the cell suspension were transferred to another tube supplemented with 5 µL DEAB (N,N-diethylaminobenzaldehyde) and pipetted to mix evenly. Both tubes (± DEAB) were then incubated at 37 °C for 45 min. Cells were washed twice with 2 ml ALDEFLUOR buffer and resuspended in 500 µL ALDEFLUOR buffer supplemented with DAPI to stain for dead cells, followed by analysis by flow cytometry with BD FACSCelesta (BD Biosciences) equipped with a 405/448 channel to delineate dead cells (filter 450/40 nm) and a 488/513 channel to collect the signal of the fluorescent dye used in ALDEFLUOR assay (filter 530/30 nm). The percentage of ALDH-positive live cells was determined based on fluorescence intensity, with gating established using DEAB-treated samples to set the baseline for ALDH activity. Data analysis was performed using FlowJo software (FlowJo LLC, Ashland, OR) (Figure S7a, Supporting Information).

**Total mRNA extraction and RT-qPCR**

RNeasy mini kit (Qiagen) was used for RNA extraction. Purified RNA, quantified with the Nanodrop spectrophotometer (Thermo Fisher Scientific), was retrotranscribed to cDNA using random primers and SuperScript VILO cDNA Synthesis Kit (Invitrogen). All procedures were performed following the manufacturer's instructions. RT-qPCR was performed with TaqMan probes (Thermo Fisher Scientific) using TaqMan Universal PCR Master Mix. Each gene was tested in technical replicates ($n \geq 2$). The ΔCt method was used to calculate the mRNA levels of target genes normalized against housekeeping genes. The $2^{-\Delta\Delta Ct}$ method was used to compare the mRNA levels of target genes, normalized to the housekeeping genes, relative to an external standard. Taqman Gene Expression Assay IDs are: NUMB (Hs01105433_m1), RBX1 (Hs00360274_m1), FBXW8 (Hs00395481_m1), ANKRD1 (Hs00173317_m1), CYR61 (hs00998500_g1), FN1 (hs00365052_m1), VIM (hs00185584_m1), Hey1 (hs00232618_m1), NOTCH1 (hs00413187_m1), NOTCH2 (hs00225747_m1), NOTCH3 (hs00166432_m1), JAG1 (hs00164982_m1), JAG2 (hs00171432_m1), B2M (Hs99999907_m1), TBP (Hs00427621_m1), HPRT1 (Hs02800695_m1), GAPDH (Hs03929097_g1), GUSB (Hs99999908_m1), ACTB (Hs01060665_g1) (Figure 2b, 3c, 7c and Figure S3, S4c, S5d,e, S7b, Supporting Information).

**RNAseq**

Total cellular RNA was extracted and quality assessed using the Bioanalyzer 2100 (Agilent). For each sample, total RNA was depleted of ribosomal RNA and the RNAseq libraries were prepared with the Illumina TruSeq Stranded Total RNA kit following the manufacturer's protocol. Following adapter ligation, libraries were amplified by PCR. Amplified libraries were checked on a Bioanalyzer 2100 and quantified with picogreen reagent (Invitrogen) and sequenced for 100 bases in the paired-end mode with 50 million reads coverage on a Novaseq 6000 sequencer. Raw data were acquired for all datasets, and the human reference genome (hg38) was employed as the alignment template for mapping the reads through Bowtie2 (version 2.4.5).[2] The estimation of gene expression abundance was performed using RSEM (version 1.3.3) with default parameters.[3]

### SUPPORTING TABLES

**Table S1. List of genes tested in the siRNA screening to identify Ub conjugation associated proteins involved in NUMB hyper-degradation in BC cells.** siGENOME® SMARTpool® siRNA Libraries (Human Ubiquitin Conjugation subset 1-3) contained siRNAs targeting 597 genes. Among these, 493 genes are unequivocally involved in the ubiquitination process (assessed by interrogating the Gene database at NCBI, https://www.ncbi.nlm.nih.gov/gene/), 87 genes are not known to be involved in the ubiquitination process, and 17 are pseudogenes or entries that were subsequently withdrawn from the NCBI database. Reported in the table: i) Pool number (original library), siRNA pool number identifier; ii) Gene symbol (original library), gene symbol as present in the original library; iii) HUGO, official HUGO nomenclature (HGNC, HUGO gene nomenclature committee, https://www.genenames.org/); iv) Official full name, descriptive official name (HGNC); v) Ubiquitination, function directly correlated to ubiquitination process.

| Pool Number (original library) | Gene Symbol (original library) | HUGO | Official full name | Ubiquitination |
|---|---|---|---|---|
| M-006522-01 | AMFR | AMFR | Autocrine motility factor receptor | YES |
| M-006992-02 | ANAPC11 | ANAPC11 | Anaphase promoting complex subunit 11 | YES |
| M-028950-01 | ANKIB1 | ANKIB1 | Ankyrin repeat and IBR domain containing 1 | YES |
| M-007184-02 | KIAA0317 | AREL1 | Apoptosis resistant E3 ubiquitin protein ligase 1 | YES |
| M-019984-00 | ARIH1 | ARIH1 | Ariadne RBR E3 ubiquitin protein ligase 1 | YES |
| M-020104-01 | ARIH2 | ARIH2 | Ariadne RBR E3 ubiquitin protein ligase 2 | YES |
| M-015579-02 | RNF165 | ARK2C | Arkadia (RNF111) C-terminal like ring finger ubiquitin ligase 2C | YES |
| M-017197-01 | ASB1 | ASB1 | Ankyrin repeat and SOCS box containing 1 | YES |
| M-007725-01 | ASB10 | ASB10 | Ankyrin repeat and SOCS box containing 10 | YES |
| M-013267-02 | ASB11 | ASB11 | Ankyrin repeat and SOCS box containing 11 | YES |
| M-013180-00 | ASB12 | ASB12 | Ankyrin repeat and SOCS box containing | YES |
| M-012929-00 | ASB13 | ASB13 | Ankyrin repeat and SOCS box containing | YES |
| M-013205-01 | ASB14 | ASB14 | Ankyrin repeat and SOCS box containing | YES |
| M-031158-01 | ASB15 | ASB15 | Ankyrin repeat and SOCS box containing | YES |
| M-013061-00 | ASB16 | ASB16 | Ankyrin repeat and SOCS box containing | YES |
| M-015284-00 | ASB17 | ASB17 | Ankyrin repeat and SOCS box containing | YES |

| | | | | |
|---|---|---|---|---|
| M-032545-01 | ASB18 | ASB18 | Ankyrin repeat and SOCS box containing | YES |
| M-009575-00 | ASB2 | ASB2 | Ankyrin repeat and SOCS box containing | YES |
| M-017457-00 | ASB3 | ASB3 | Ankyrin repeat and SOCS box containing | YES |
| M-013339-00 | ASB4 | ASB4 | Ankyrin repeat and SOCS box containing | YES |
| M-017458-01 | ASB5 | ASB5 | Ankyrin repeat and SOCS box containing | YES |
| M-013355-00 | ASB6 | ASB6 | Ankyrin repeat and SOCS box containing | YES |
| M-012930-00 | ASB7 | ASB7 | Ankyrin repeat and SOCS box containing | YES |
| M-014301-00 | ASB8 | ASB8 | Ankyrin repeat and SOCS box containing | YES |
| M-014295-00 | ASB9 | ASB9 | Ankyrin repeat and SOCS box containing | YES |
| M-003873-02 | BARD1 | BARD1 | BRCA1 associated RING domain 1 | YES |
| M-004386-01 | BFAR | BFAR | Bifunctional apoptosis regulator | YES |
| M-004390-02 | BIRC2 | BIRC2 | Baculoviral IAP repeat containing 2 | YES |
| M-004099-02 | BIRC3 | BIRC3 | Baculoviral IAP repeat containing 3 | YES |
| M-013857-01 | BIRC6 | BIRC6 | Baculoviral IAP repeat containing 6 | YES |
| M-004391-03 | BIRC7 | BIRC7 | Baculoviral IAP repeat containing 7 | YES |
| M-004392-00 | BIRC8 | BIRC8 | Baculoviral IAP repeat containing 8 | YES |
| M-005230-00 | BMI1 | BMI1 | BMI1 proto-oncogene, polycomb ring finger | YES |
| M-006597-01 | BRAP | BRAP | BRCA1 associated protein | YES |
| M-003461-02 | BRCA1 | BRCA1 | BRCA1 DNA repair associated | YES |
| M-003463-01 | BTRC | BTRC | Beta-transducin repeat containing E3 ubiquitin protein ligase | YES |
| M-016305-01 | C10ORF46 | CACUL1 | CDK2 associated cullin domain 1 | YES |
| M-015562-00 | TIP120A | CAND1 | Cullin associated and neddylation dissociated 1 | YES |
| M-023448-01 | CAND2 | CAND2 | Cullin associated and neddylation dissociated 2 (putative) | YES |
| M-003003-02 | CBL | CBL | Cbl proto-oncogene | YES |
| M-003004-02 | CBLB | CBLB | Cbl proto-oncogene B | YES |
| M-006962-00 | CBLC | CBLC | Cbl proto-oncogene C | YES |
| M-007069-00 | CBLL1 | CBLL1 | Cbl proto-oncogene like 1 | YES |
| M-007148-02 | ZNF645 | CBLL2 | Cbl proto-oncogene like 2 | YES |
| M-003215-02 | CCNF | CCNF | Cyclin F | YES |

| | | | | |
|---|---|---|---|---|
| M-003230-01 | CDC34 | CDC34 | Cell division cycle 34, ubiqiutin conjugating enzyme | YES |
| M-006933-01 | CGRRF1 | CGRRF1 | Cell growth regulator with ring finger domain 1 | YES |
| M-007018-01 | CHFR | CHFR | Checkpoint with forkhead and ring finger domains | YES |
| M-017381-01 | CISH | CISH | Cytokine inducible SH2 containing protein | YES |
| M-020323-01 | CNOT4 | CNOT4 | CCR4-NOT transcription complex subunit 4 | YES |
| M-007049-01 | RFWD2 | COP1 | COP1 E3 ubiquitin ligase | YES |
| M-004086-01 | CUL1 | CUL1 | Cullin 1 | YES |
| M-007277-00 | CUL2 | CUL2 | Cullin 2 | YES |
| M-010224-02 | CUL3 | CUL3 | Cullin 3 | YES |
| M-012610-01 | CUL4A | CUL4A | Cullin 4A | YES |
| M-017965-01 | CUL4B | CUL4B | Cullin 4B | YES |
| M-019553-01 | CUL5 | CUL5 | Cullin 5 | YES |
| M-017673-00 | CUL7 | CUL7 | Cullin 7 | YES |
| M-014128-01 | PARC | CUL9 | Cullin 9 | YES |
| M-019139-01 | DCUN1D1 | DCUN1D1 | Defective in cullin neddylation 1 domain containing 1 | YES |
| M-020261-01 | DCUN1D2 | DCUN1D2 | Defective in cullin neddylation 1 domain containing 2 | YES |
| M-018390-01 | DCUN1D3 | DCUN1D3 | Defective in cullin neddylation 1 domain containing 3 | YES |
| M-014118-02 | DCUN1D4 | DCUN1D4 | Defective in cullin neddylation 1 domain containing 4 | YES |
| M-014842-01 | DCUN1D5 | DCUN1D5 | Defective in cullin neddylation 1 domain containing 5 | YES |
| M-006525-00 | DTX1 | DTX1 | Deltex E3 ubiquitin ligase 1 | YES |
| M-007114-02 | DTX2 | DTX2 | Deltex E3 ubiquitin ligase 2 | YES |
| M-007156-01 | DTX3 | DTX3 | Deltex E3 ubiquitin ligase 3 | YES |
| M-007143-01 | DTX3L | DTX3L | Deltex E3 ubiquitin ligase 3L | YES |
| M-026663-01 | DTX4 | DTX4 | Deltex E3 ubiquitin ligase 4 | YES |
| M-006910-00 | DZIP3 | DZIP3 | DAZ interacting zinc finger protein 3 | YES |
| M-017404-00 | FBXO18 | FBH1 | F-box DNA helicase 1 | YES |
| M-005204-01 | FBXL12 | FBXL12 | F-box and leucine rich repeat protein 12 | YES |
| M-016001-01 | FBXL13 | FBXL13 | F-box and leucine rich repeat protein 13 | YES |
| M-015718-00 | FBXL14 | FBXL14 | F-box and leucine rich repeat protein 14 | YES |

| | | | | |
|---|---|---|---|---|
| M-026308-01 | FBXL15 | FBXL15 | F-box and leucine rich repeat protein 15 | YES |
| M-016797-00 | FBXL16 | FBXL16 | F-box and leucine rich repeat protein 16 | YES |
| M-024385-00 | FBXL17 | FBXL17 | F-box and leucine rich repeat protein 17 | YES |
| M-016587-01 | FBXL18 | FBXL18 | F-box and leucine rich repeat protein 18 | YES |
| M-031874-01 | FBXL19 | FBXL19 | F-box and leucine rich repeat protein 19 | YES |
| M-013562-01 | FBXL2 | FBXL2 | F-box and leucine rich repeat protein 2 | YES |
| M-015029-00 | FBXL20 | FBXL20 | F-box and leucine rich repeat protein 20 | YES |
| M-012423-02 | FBXL3P | FBXL21P | F-box and leucine rich repeat protein 21, pseudogene | YES |
| M-031751-00 | FBXL22 | FBXL22 | F-box and leucine rich repeat protein 22 | YES |
| M-012422-00 | FBXL3A | FBXL3 | F-box and leucine rich repeat protein 3 | YES |
| M-013564-00 | FBXL4 | FBXL4 | F-box and leucine rich repeat protein 4 | YES |
| M-012424-01 | FBXL5 | FBXL5 | F-box and leucine rich repeat protein 5 | YES |
| M-012425-02 | FBXL6 | FBXL6 | F-box and leucine rich repeat protein 6 | YES |
| M-012457-00 | FBXL7 | FBXL7 | F-box and leucine rich repeat protein 7 | YES |
| M-017504-00 | FBXL8 | FBXL8 | F-box and leucine rich repeat protein 8 | YES |
| M-012426-01 | LRRC29 | FBXL9P | F-box and leucine rich repeat protein 9, pseudogene | YES |
| M-026138-02 | FBXO10 | FBXO10 | F-box protein 10 | YES |
| M-012428-00 | FBXO11 | FBXO11 | F-box protein 11 | YES |
| M-016503-00 | FBXO15 | FBXO15 | F-box protein 15 | YES |
| M-017447-00 | FBXO16 | FBXO16 | F-box protein 16 | YES |
| M-013128-00 | FBXO17 | FBXO17 | F-box protein 17 | YES |
| M-012429-00 | FBXO2 | FBXO2 | F-box protein 2 | YES |
| M-012917-01 | FBXO21 | FBXO21 | F-box protein 21 | YES |
| M-010812-01 | FBXO22 | FBXO22 | F-box protein 22 | YES |
| M-012430-01 | FBXO24 | FBXO24 | F-box protein 24 | YES |
| M-019192-00 | FBXO25 | FBXO25 | F-box protein 25 | YES |
| M-018558-00 | FBXO27 | FBXO27 | F-box protein 27 | YES |
| M-014080-00 | FBXO28 | FBXO28 | F-box protein 28 | YES |
| M-012432-00 | FBXO3 | FBXO3 | F-box protein 3 | YES |

| | | | | |
|---|---|---|---|---|
| M-021407-01 | FBXO30 | FBXO30 | F-box protein 30 | YES |
| M-016541-01 | FBXO31 | FBXO31 | F-box protein 31 | YES |
| M-013005-02 | FBXO32 | FBXO32 | F-box protein 32 | YES |
| M-021948-00 | FBXO33 | FBXO33 | F-box protein 33 | YES |
| M-020989-00 | FBXO34 | FBXO34 | F-box protein 34 | YES |
| M-018460-01 | FBXO36 | FBXO36 | F-box protein 36 | YES |
| M-018163-01 | FBXO38 | FBXO38 | F-box protein 38 | YES |
| M-016871-00 | FBXO39 | FBXO39 | F-box protein 39 | YES |
| M-012433-01 | FBXO4 | FBXO4 | F-box protein 4 | YES |
| M-020905-01 | FBXO40 | FBXO40 | F-box protein 40 | YES |
| M-022720-01 | FBXO41 | FBXO41 | F-box protein 41 | YES |
| M-022191-01 | FBXO42 | FBXO42 | F-box protein 42 | YES |
| M-025904-02 | FBXO43 | FBXO43 | F-box protein 43 | YES |
| M-019201-01 | FBXO44 | FBXO44 | F-box protein 44 | YES |
| M-023542-02 | LOC200933 | FBXO45 | F-box protein 45 | YES |
| M-023753-01 | FBXO46 | FBXO46 | F-box protein 46 | YES |
| M-034909-00 | FBXO47 | FBXO47 | F-box protein 47 | YES |
| M-034947-00 | LOC554251 | FBXO48 | F-box protein 48 | YES |
| M-012434-01 | FBXO5 | FBXO5 | F-box protein 5 | YES |
| M-013314-00 | FBXO6 | FBXO6 | F-box protein 6 | YES |
| M-013606-01 | FBXO7 | FBXO7 | F-box protein 7 | YES |
| M-012435-00 | FBXO8 | FBXO8 | F-box protein 8 | YES |
| M-012469-01 | FBXO9 | FBXO9 | F-box protein 9 | YES |
| M-014733-01 | FBXW10 | FBXW10 | F-box and WD repeat domain containing 10 | YES |
| M-003490-01 | FBXW11 | FBXW11 | F-box and WD repeat domain containing 11 | YES |
| M-032001-00 | FBXW12 | FBXW12 | F-box and WD repeat domain containing 12 | YES |
| M-012427-01 | FBXW2 | FBXW2 | F-box and WD repeat domain containing 2 | YES |
| M-013956-01 | SHFM3 | FBXW4 | F-box and WD repeat domain containing 4 | YES |
| M-013389-01 | FBXW5 | FBXW5 | F-box and WD repeat domain containing 5 | YES |

| | | | | |
|---|---|---|---|---|
| M-004264-02 | FBXW7 | FBXW7 | F-box and WD repeat domain containing 7 | YES |
| M-012431-02 | FBXW8 | FBXW8 | F-box and WD repeat domain containing 8 | YES |
| M-014844-01 | FBXW9 | FBXW9 | F-box and WD repeat domain containing 9 | YES |
| M-007014-00 | KIAA1333 | G2E3 | G2/M-phase specific E3 ubiquitin protein ligase | YES |
| M-007193-01 | HACE1 | HACE1 | HECT domain and ankyrin repeat containing E3 ubiquitin protein ligase 1 | YES |
| M-007188-02 | HECTD1 | HECTD1 | HECT domain E3 ubiquitin protein ligase 1 | YES |
| M-007198-00 | HECTD2 | HECTD2 | HECT domain E3 ubiquitin protein ligase 2 | YES |
| M-027468-03 | HECTD3 | HECTD3 | HECT domain E3 ubiquitin protein ligase 3 | YES |
| M-018270-01 | FLJ34154 | HECTD4 | HECT domain E3 ubiquitin protein ligase 4 | YES |
| M-007186-01 | HECW1 | HECW1 | HECT, C2 and WW domain containing E3 ubiquitin protein ligase 1 | YES |
| M-007192-00 | HECW2 | HECW2 | HECT, C2 and WW domain containing E3 ubiquitin protein ligase 2 | YES |
| M-007181-02 | HERC1 | HERC1 | HECT and RLD domain containing E3 ubiquitin protein ligase family member 1 | YES |
| M-007180-02 | HERC2 | HERC2 | HECT and RLD domain containing E3 ubiquitin protein ligase family member 2 | YES |
| M-007179-01 | HERC3 | HERC3 | HECT and RLD domain containing E3 ubiquitin protein ligase family member 3 | YES |
| M-021426-01 | HERC4 | HERC4 | HECT and RLD domain containing E3 ubiquitin protein ligase family member 4 | YES |
| M-005174-02 | HERC5 | HERC5 | HECT and RLD domain containing E3 ubiquitin protein ligase family member 5 | YES |
| M-005175-03 | HERC6 | HERC6 | HECT and RLD domain containing E3 ubiquitin protein ligase family member 6 | YES |
| M-006448-02 | SMARCA3 | HLTF | Helicase like transcription factor | YES |
| M-007185-01 | HUWE1 | HUWE1 | HECT, UBA and WWE domain containing E3 ubiquitin protein ligase 1 | YES |
| M-006969-00 | IRF2BP1 | IRF2BP1 | Interferon regulatory factor 2 binding protein 1 | YES |
| M-007196-01 | ITCH | ITCH | Itchy E3 ubiquitin protein ligase | YES |
| M-019252-02 | LMO7 | LMO7 | LIM domain 7 | YES |
| M-014940-01 | LNX1 | LNX1 | Ligand of numb-protein X 1 | YES |
| M-007164-01 | LNX2 | LNX2 | Ligand of numb-protein X 2 | YES |
| M-007108-01 | LONRF1 | LONRF1 | LON peptidase N-terminal domain and ring finger 1 | YES |
| M-027218-02 | LONRF2 | LONRF2 | LON peptidase N-terminal domain and ring finger 2 | YES |
| M-007067-01 | LONRF3 | LONRF3 | LON peptidase N-terminal domain and ring finger 3 | YES |
| M-007103-01 | LRSAM1 | LRSAM1 | Leucine rich repeat and sterile alpha motif containing 1 | YES |
| M-006968-00 | ZNF294 | LTN1 | Listerin E3 ubiquitin protein ligase 1 | YES |

| | | | | |
|---|---|---|---|---|
| M-003575-02 | MAP3K1 | MAP3K1 | Mitogen-activated protein kinase kinase kinase 1 | YES |
| M-007006-02 | MARCH1 | MARCHF1 | Membrane associated ring-CH-type finger 1 | YES |
| M-007150-02 | RNF190 | MARCHF10 | Membrane associated ring-CH-type finger 10 | YES |
| M-033939-01 | LOC441061 | MARCHF11 | Membrane associated ring-CH-type finger 11 | YES |
| M-006986-02 | MARCH2 | MARCHF2 | Membrane associated ring-CH-type finger 2 | YES |
| M-007116-02 | MARCH3 | MARCHF3 | Membrane associated ring-CH-type finger 3 | YES |
| M-023172-01 | MARCH4 | MARCHF4 | Membrane associated ring-CH-type finger 4 | YES |
| M-007001-01 | MARCH5 | MARCHF5 | Membrane associated ring-CH-type finger 5 | YES |
| M-006925-00 | MARCH6 | MARCHF6 | Membrane associated ring-CH-type finger 6 | YES |
| M-007055-01 | MARCH7 | MARCHF7 | Membrane associated ring-CH-type finger 7 | YES |
| M-007161-03 | MARCH8 | MARCHF8 | Membrane associated ring-CH-type finger 8 | YES |
| M-015561-01 | MARCH9 | MARCHF9 | Membrane associated ring-CH-type finger 9 | YES |
| M-003279-04 | MDM2 | MDM2 | MDM2 proto-oncogene | YES |
| M-006536-03 | MDM4 | MDM4 | MDM4 regulator of p53 | YES |
| M-011081-00 | MEFV | MEFV | MEFV innate immunity regulator, pyrin | YES |
| M-022355-01 | LOC92312 | MEX3A | Mex-3 RNA binding family member A | YES |
| M-022292-00 | RKHD3 | MEX3B | Mex-3 RNA binding family member B | YES |
| M-006989-01 | RKHD2 | MEX3C | Mex-3 RNA binding family member C | YES |
| M-031965-01 | RKHD1 | MEX3D | Mex-3 RNA binding family member D | YES |
| M-022620-00 | MGRN1 | MGRN1 | Mahogunin ring finger 1 | YES |
| M-014033-01 | MIB1 | MIB1 | MIB E3 ubiquitin protein ligase 1 | YES |
| M-015287-02 | MIB2 | MIB2 | MIB E3 ubiquitin protein ligase 2 | YES |
| M-006537-01 | MID1 | MID1 | Midline 1 | YES |
| M-006938-00 | MID2 | MID2 | Midline 2 | YES |
| M-006959-00 | MKRN1 | MKRN1 | Makorin ring finger protein 1 | YES |
| M-006960-01 | MKRN2 | MKRN2 | makorin ring finger protein 2 | YES |
| M-006581-01 | MKRN3 | MKRN3 | makorin ring finger protein 3 | YES |
| M-020828-01 | MSL2L1 | MSL2 | MSL complex subunit 2 | YES |
| M-007062-02 | C1orf166 | MUL1 | Mitochondrial E3 ubiquitin protein ligase 1 | YES |

| | | | | |
|---|---|---|---|---|
| M-006951-01 | MYCBP2 | MYCBP2 | MYC binding protein 2 | YES |
| M-032307-01 | LOC342897 | NCCRP1 | NCCRP1, F-box associated domain containing | YES |
| M-007178-02 | NEDD4 | NEDD4 | NEDD4 E3 ubiquitin protein ligase | YES |
| M-007187-02 | NEDD4L | NEDD4L | NEDD4 like E3 ubiquitin protein ligase | YES |
| M-016715-00 | NEURL | NEURL1 | Neuralized E3 ubiquitin protein ligase 1 | YES |
| M-015269-00 | NEURL2 | NEURL2 | Neuralized E3 ubiquitin protein ligase 2 | YES |
| M-015556-01 | LINCR | NEURL3 | Neuralized E3 ubiquitin protein ligase 3 | YES |
| M-027323-00 | NHLRC1 | NHLRC1 | NHL repeat containing E3 ubiquitin protein ligase 1 | YES |
| M-016380-01 | C13ORF7 | OBI1 | ORC ubiquitin ligase 1p | YES |
| M-008670-01 | ZA20D1 | OTUD7B | OTU deubiquitinase 7B | YES |
| M-013539-00 | ZNF278 | PATZ1 | POZ/BTB and AT hook containing zinc finger 1 | YES |
| M-007094-02 | PCGF1 | PCGF1 | Polycomb group ring finger 1 | YES |
| M-006584-02 | PCGF2 | PCGF2 | Polycomb group ring finger 2 | YES |
| M-006926-01 | PCGF3 | PCGF3 | Polycomb group ring finger 3 | YES |
| M-007089-01 | PCGF5 | PCGF5 | Polycomb group ring finger 5 | YES |
| M-007084-01 | PCGF6 | PCGF6 | Polycomb group ring finger 6 | YES |
| M-023265-01 | PDZRN3 | PDZRN3 | PDZ domain containing ring finger 3 | YES |
| M-020442-01 | PDZRN4 | PDZRN4 | PDZ domain containing ring finger 4 | YES |
| M-032200-03 | KUA-UEV | PEDS1-UBE2V1 | PEDS1-UBE2V1 readthrough | YES |
| M-006545-00 | PEX10 | PEX10 | Peroxisomal biogenesis factor 10 | YES |
| M-006548-02 | PXMP3 | PEX2 | peroxisomal biogenesis factor 2 | YES |
| M-026727-00 | KIAA1542 | PHRF1 | PHD and ring finger domains 1 | YES |
| M-007045-01 | PJA1 | PJA1 | Praja ring finger ubiquitin ligase 1 | YES |
| M-006916-00 | PJA2 | PJA2 | Praja ring finger ubiquitin ligase 2 | YES |
| M-006547-01 | PML | PML | PML nuclear body scaffold | YES |
| M-007205-01 | PPIL2 | PPIL2 | Peptidylprolyl isomerase like 2 | YES |
| M-003603-00 | PARK2 | PRKN | Parkin RBR E3 ubiquitin protein ligase | YES |
| M-004668-02 | PRPF19 | PRPF19 | Pre-mRNA processing factor 19 | YES |
| M-008924-02 | RAB40A | RAB40A | RAB40A, member RAS oncogene family | YES |

| | | | | |
|---|---|---|---|---|
| M-008353-00 | RAB40B | RAB40B | RAB40B, member RAS oncogene family | YES |
| M-010368-00 | RAB40C | RAB40C | RAB40C, member RAS oncogene family | YES |
| M-004591-00 | RAD18 | RAD18 | RAD18 E3 ubiquitin protein ligase | YES |
| M-006549-03 | RAG1 | RAG1 | Recombination activating 1 | YES |
| M-006551-00 | RBBP6 | RBBP6 | RB binding protein 6, ubiquitin ligase | YES |
| M-006932-02 | C20ORF18 | RBCK1 | RANBP2-type and C3HC4-type zinc finger containing 1 | YES |
| M-004087-01 | RBX1 | RBX1 | Ring-box 1 | YES |
| M-021878-02 | RC3H1 | RC3H1 | Ring finger and CCCH-type domains 1 | YES |
| M-020453-02 | RC3H2 | RC3H2 | Ring finger and CCCH-type domains 2 | YES |
| M-006966-01 | RCHY1 | RCHY1 | Ring finger and CHY zinc finger domain containing 1 | YES |
| M-007120-03 | RFFL | RFFL | Ring finger and FYVE like domain containing E3 ubiquitin protein ligase | YES |
| M-006553-01 | RFPL1 | RFPL1 | Ret finger protein like 1 | YES |
| M-006935-01 | RFPL2 | RFPL2 | Ret finger protein like 2 | YES |
| M-006934-01 | RFPL3 | RFPL3 | Ret finger protein like 3 | YES |
| M-180651-01 | LOC729974 | RFPL4AL1 | Ret finger protein like 4A like 1 | YES |
| M-032292-03 | RFPL4B | RFPL4B | Ret finger protein like 4B | YES |
| M-017095-01 | RFWD3 | RFWD3 | Ring finger and WD repeat domain 3 | YES |
| M-006554-03 | RING1 | RING1 | Ring finger protein 1 | YES |
| M-006982-01 | RNF12 | RLIM | Ring finger protein, LIM domain interacting | YES |
| M-006918-01 | RNF10 | RNF10 | Ring finger protein 10 | YES |
| M-006594-00 | RNF103 | RNF103 | Ring finger protein 103 | YES |
| M-006971-01 | RNF11 | RNF11 | Ring finger protein 11 | YES |
| M-007002-01 | RNF111 | RNF111 | Ring finger protein 111 | YES |
| M-006588-01 | ZNF179 | RNF112 | Ring finger protein 112 | YES |
| M-006590-01 | RNF113A | RNF113A | Ring finger protein 113A | YES |
| M-007135-02 | RNF113B | RNF113B | Ring finger protein 113B | YES |
| M-007024-00 | ZNF313 | RNF114 | Ring finger protein 114 | YES |
| M-006974-00 | ZNF364 | RNF115 | Ring finger protein 115 | YES |
| M-007011-02 | RNF121 | RNF121 | Ring finger protein 121 | YES |

| | | | | |
|---|---|---|---|---|
| M-007068-00 | RNF122 | RNF122 | Ring finger protein 122 | YES |
| M-007041-00 | RNF123 | RNF123 | Ring finger protein 123 | YES |
| M-007005-01 | RNF125 | RNF125 | Ring finger protein 125 | YES |
| M-007015-01 | RNF126 | RNF126 | Ring finger protein 126 | YES |
| M-007061-01 | RNF128 | RNF128 | Ring finger protein 128 | YES |
| M-006944-01 | RNF13 | RNF13 | Ring finger protein 13 | YES |
| M-007021-00 | RNF130 | RNF130 | Ring finger protein 130 | YES |
| M-007155-00 | RNF133 | RNF133 | Ring finger protein 133 | YES |
| M-007087-01 | RNF135 | RNF135 | Ring finger protein 135 | YES |
| M-006991-01 | RNF138 | RNF138 | Ring finger protein 138 | YES |
| M-006942-01 | RNF139 | RNF139 | Ring finger protein 139 | YES |
| M-006906-02 | RNF14 | RNF14 | Ring finger protein 14 | YES |
| M-006980-00 | RNF141 | RNF141 | Ring finger protein 141 | YES |
| M-006912-01 | RNF144 | RNF144A | Ring finger protein 144A | YES |
| M-025119-01 | IBRDC2 | RNF144B | Ring finger protein 144B | YES |
| M-007146-01 | FLJ31951 | RNF145 | Ring finger protein 145 | YES |
| M-007080-00 | RNF146 | RNF146 | Ring finger protein 146 | YES |
| M-010758-00 | RNF148 | RNF148 | Ring finger protein 148 | YES |
| M-007169-00 | RNF149 | RNF149 | Ring finger protein 149 | YES |
| M-010713-01 | RNF150 | RNF150 | Ring finger protein 150 | YES |
| M-030777-01 | RNF151 | RNF151 | Ring finger protein 151 | YES |
| M-007160-01 | RNF152 | RNF152 | Ring finger protein 152 | YES |
| M-022965-01 | RNF157 | RNF157 | Ring finger protein 157 | YES |
| M-007119-00 | RNF166 | RNF166 | Ring finger protein 166 | YES |
| M-006967-01 | RNF167 | RNF167 | Ring finger protein 167 | YES |
| M-007152-03 | RNF168 | RNF168 | Ring finger protein 168 | YES |
| M-032290-02 | RNF169 | RNF169 | Ring finger protein 169 | YES |
| M-007026-01 | RNF17 | RNF17 | Ring finger protein 17 | YES |
| M-007078-01 | RNF170 | RNF170 | Ring finger protein 170 | YES |

| | | | | |
|---|---|---|---|---|
| M-007170-02 | RNF175 | RNF175 | Ring finger protein 175 | YES |
| M-017714-01 | RNF180 | RNF180 | Ring finger protein 180 | YES |
| M-007162-02 | RNF182 | RNF182 | Ring finger protein 182 | YES |
| M-007134-02 | RNF183 | RNF183 | Ring finger protein 183 | YES |
| M-007107-01 | RNF185 | RNF185 | Ring finger protein 185 | YES |
| M-006999-01 | RNF186 | RNF186 | Ring finger protein 186 | YES |
| M-021939-01 | RNF187 | RNF187 | Ring finger protein 187 | YES |
| M-006965-00 | RNF19 | RNF19 | Ring finger protein 19 | YES |
| M-010203-00 | IBRDC3 | RNF19B | Ring finger protein 19B | YES |
| M-006556-01 | RNF2 | RNF2 | Ring finger protein 2 | YES |
| M-007027-00 | RNF20 | RNF20 | Ring finger protein 20 | YES |
| M-032070-01 | RNF207 | RNF207 | Ring finger protein 207 | YES |
| M-180652-00 | RNF208 | RNF208 | Ring finger protein 208 | YES |
| M-023324-02 | C17ORF27 | RNF213 | Ring finger protein 213 | YES |
| M-026520-01 | DKFZP547C195 | RNF214 | Ring finger protein 214 | YES |
| M-024565-01 | RNF215 | RNF215 | Ring finger protein 215 | YES |
| M-017305-00 | TRIAD3 | RNF216 | Ring finger protein 216 | YES |
| M-007147-00 | IBRDC1 | RNF217 | Ring finger protein 217 | YES |
| M-021044-01 | C1orf164 | RNF220 | Ring finger protein 220 | YES |
| M-036200-00 | LOC643904 | RNF222 | Ring finger protein 222 | YES |
| M-006943-00 | RNF24 | RNF24 | Ring finger protein 24 | YES |
| M-007047-00 | RNF25 | RNF25 | Ring finger protein 25 | YES |
| M-007060-01 | RNF26 | RNF26 | Ring finger protein 26 | YES |
| M-021419-01 | RNF31 | RNF31 | Ring finger protein 31 | YES |
| M-007136-01 | RNF32 | RNF32 | Ring finger protein 32 | YES |
| M-007072-00 | RNF34 | RNF34 | Ring finger protein 33 | YES |
| M-007144-01 | RNF38 | RNF38 | Ring finger protein 38 | YES |
| M-007074-00 | RNF39 | RNF39 | Ring finger protein 39 | YES |
| M-006557-03 | RNF4 | RNF4 | Ring finger protein 4 | YES |

| | | | | |
|---|---|---|---|---|
| M-006913-00 | RNF40 | RNF40 | Ring finger protein 40 | YES |
| M-006922-01 | RNF41 | RNF41 | Ring finger protein 41 | YES |
| M-007004-02 | RNF43 | RNF43 | Ring finger protein 43 | YES |
| M-006947-00 | RNF44 | RNF44 | Ring finger protein 44 | YES |
| M-006558-02 | RNF5 | RNF5 | Ring finger protein 5 | YES |
| M-006559-01 | RNF6 | RNF6 | Ring finger protein 6 | YES |
| M-006907-02 | RNF7 | RNF7 | Ring finger protein 7 | YES |
| M-006900-01 | RNF8 | RNF8 | Ring finger protein 8 | YES |
| M-007097-01 | FLJ14627 | RNFT2 | Ring finger protein, transmembrane 2 | YES |
| M-010205-01 | LOC51136 | RNTF1 | Ring finger protein, transmembrane 1 | YES |
| M-006985-01 | LOC51255 | RNTF181 | Ring finger protein 181 | YES |
| M-022564-02 | RSPRY1 | RSPRY1 | Ring finger and SPRY domain containing 1 | YES |
| M-007037-01 | SH3MD2 | SH3RF1 | SH3 domain containing ring finger 1 | YES |
| M-007145-00 | SH3RF2 | SH3RF2 | SH3 domain containing ring finger 2 | YES |
| M-027806-01 | SH3MD4 | SH3RF3 | SH3 domain containing ring finger 3 | YES |
| M-007167-01 | SHPRH | SHPRH | SNF2 histone linker PHD RING helicase | YES |
| M-006561-02 | SIAH2 | SIAH2 | Siah E3 ubiquitin protein ligase 2 | YES |
| M-003324-04 | SKP2 | SKP2 | S-phase kinase associated protein 2 | YES |
| M-007191-01 | SMURF1 | SMURF1 | SMAD specific E3 ubiquitin protein ligase 1 | YES |
| M-007194-01 | SMURF2 | SMURF2 | SMAD specific E3 ubiquitin protein ligase 2 | YES |
| M-011511-04 | SOCS1 | SOCS1 | Suppressor of cytokine signaling 1 | YES |
| M-017604-00 | SOCS2 | SOCS2 | Suppressor of cytokine signaling 2 | YES |
| M-004299-02 | SOCS3 | SOCS3 | Suppressor of cytokine signaling 3 | YES |
| M-009037-01 | SOCS4 | SOCS4 | Suppressor of cytokine signaling 4 | YES |
| M-017374-00 | SOCS5 | SOCS5 | Suppressor of cytokine signaling 5 | YES |
| M-017375-01 | SOCS6 | SOCS6 | Suppressor of cytokine signaling 6 | YES |
| M-027197-00 | SOCS7 | SOCS7 | Suppressor of cytokine signaling 7 | YES |
| M-015262-01 | SPSB1 | SPSB1 | SplA/ryanodine receptor domain and SOCS box containing 1 | YES |
| M-014947-02 | SPSB2 | SPSB2 | splA/ryanodine receptor domain and SOCS box containing 2 | YES |

| | | | | |
|---|---|---|---|---|
| M-017713-01 | SPSB3 | SPSB3 | splA/ryanodine receptor domain and SOCS box containing 3 | YES |
| M-015283-01 | SPSB4 | SPSB4 | splA/ryanodine receptor domain and SOCS box containing 4 | YES |
| M-007201-02 | STUB1 | STUB1 | STIP1 homology and U-box containing protein 1 | YES |
| M-007090-01 | SYVN1 | SYVN1 | Synoviolin 1 | YES |
| M-009919-00 | TNFAIP3 | TNFAIP3 | TNF alpha induced protein 3 | YES |
| M-020048-00 | TOPORS | TOPORS | TOP1 binding arginine/serine rich protein, E3 ubiquitin ligase | YES |
| M-005252-02 | TRAF3 | TRAF3 | TNF receptor associated factor 3 | YES |
| M-006908-01 | TRAF4 | TRAF4 | TNF receptor associated factor 4 | YES |
| M-006568-01 | TRAF5 | TRAF5 | TNF receptor associated factor 5 | YES |
| M-004712-00 | TRAF6 | TRAF6 | TNF receptor associated factor 6 | YES |
| M-007086-00 | TRAF7 | TRAF7 | TNF receptor associated factor 7 | YES |
| M-006924-00 | TRIP | TRAIP | TRAF interacting protein | YES |
| M-006920-01 | TRIM10 | TRIM10 | Tripartite motif containing 10 | YES |
| M-007075-00 | TRIM11 | TRIM11 | Tripartite motif containing 11 | YES |
| M-006923-00 | RFP2 | TRIM13 | Tripartite motif containing 13 | YES |
| M-010976-00 | TRIM14 | TRIM14 | Tripartite motif containing 14 | YES |
| M-007102-01 | TRIM15 | TRIM15 | Tripartite motif containing 15 | YES |
| M-006981-01 | TRIM17 | TRIM17 | Tripartite motif containing 17 | YES |
| M-006955-00 | TRIM2 | TRIM2 | Tripartite motif containing 2 | YES |
| M-006563-02 | SSA1 | TRIM21 | Tripartite motif containing 21 | YES |
| M-006927-03 | TRIM22 | TRIM22 | Tripartite motif containing 22 | YES |
| M-006523-00 | TRIM23 | TRIM23 | Tripartite motif containing 23 | YES |
| M-005387-03 | TIF1 | TRIM24 | Tripartite motif containing 24 | YES |
| M-006585-00 | TRIM25 | TRIM25 | Tripartite motif containing 25 | YES |
| M-006587-01 | TRIM26 | TRIM26 | Tripartite motif containing 26 | YES |
| M-006552-01 | RFP | TRIM27 | Tripartite motif containing 27 | YES |
| M-005046-01 | TRIM28 | TRIM28 | Tripartite motif containing 28 | YES |
| M-012409-01 | TRIM29 | TRIM29 | Tripartite motif containing 29 | YES |
| M-006931-00 | TRIM3 | TRIM3 | Tripartite motif containing 3 | YES |

| | | | | |
|---|---|---|---|---|
| M-006939-01 | TRIM31 | TRIM31 | Tripartite motif containing 31 | YES |
| M-006950-01 | TRIM32 | TRIM32 | Tripartite motif containing 32 | YES |
| M-005392-03 | TRIM33 | TRIM33 | Tripartite motif containing 33 | YES |
| M-006952-02 | TRIM35 | TRIM35 | Tripartite motif containing 35 | YES |
| M-007012-01 | TRIM36 | TRIM36 | Tripartite motif containing 36 | YES |
| M-006538-02 | TRIM37 | TRIM37 | Tripartite motif containing 37 | YES |
| M-006929-01 | TRIM38 | TRIM38 | Tripartite motif containing 38 | YES |
| M-007028-01 | TRIM39 | TRIM39 | Tripartite motif containing 39 | YES |
| M-007101-00 | TRIM4 | TRIM4 | Tripartite motif containing 4 | YES |
| M-007129-01 | TRIM40 | TRIM40 | Tripartite motif containing 40 | YES |
| M-007105-02 | TRIM41 | TRIM41 | Tripartite motif containing 41 | YES |
| M-007173-00 | TRIM42 | TRIM42 | Tripartite motif containing 42 | YES |
| M-007127-00 | TRIM43 | TRIM43 | Tripartite motif containing43 | YES |
| M-039853-00 | LOC653192 | TRIM43B | Tripartite motif containing 43B | YES |
| M-007073-01 | TRIM45 | TRIM45 | Tripartite motif containing 45 | YES |
| M-007071-01 | TRIM46 | TRIM46 | Tripartite motif containing 46 | YES |
| M-007106-02 | TRIM47 | TRIM47 | Tripartite motif containing 47 | YES |
| M-007059-00 | TRIM48 | TRIM48 | Tripartite motif containing 48 | YES |
| M-007030-01 | TRIM49 | TRIM49 | Tripartite motif containing 49B | YES |
| M-026798-01 | LOC283116 | TRIM49B | Tripartite motif containing 49B | YES |
| M-007100-00 | TRIM5 | TRIM5 | Tripartite motif containing 5 | YES |
| M-007130-00 | TRIM50A | TRIM50A | Tripartite motif containing 50A | YES |
| M-010079-02 | SPRYD5 | TRIM51 | Tripartite motif containing 51 | YES |
| M-026885-01 | LOC120824 | TRIM51G | Tripartite motif-containing 51G | YES |
| M-007095-00 | TRIM52 | TRIM52 | Tripartite motif containing 52 | YES |
| M-007032-01 | TRIM54 | TRIM54 | Tripartite motif containing 54 | YES |
| M-007092-01 | TRIM55 | TRIM55 | Tripartite motif containing 55 | YES |
| M-007079-00 | TRIM56 | TRIM56 | Tripartite motif containing 56 | YES |
| M-013985-02 | TRIM58 | TRIM58 | Tripartite motif containing 58 | YES |

| | | | | |
|---|---|---|---|---|
| M-007172-01 | TRIM59 | TRIM59 | Tripartite motif containing 59 | YES |
| M-007121-01 | TRIM6 | TRIM6 | Tripartite motif containing 6 | YES |
| M-032369-00 | TRIM6-TRIM34 | TRIM6-TRIM34 | TRIM6-TRIM34 readthrough | YES |
| M-007153-00 | TRIM60 | TRIM60 | Tripartite motif containing 60 | YES |
| M-028281-01 | TRIM61 | TRIM61 | Tripartite motif containing 61 | YES |
| M-007010-01 | TRIM62 | TRIM62 | Tripartite motif containing 62 | YES |
| M-007093-01 | TRIM63 | TRIM63 | Tripartite motif containing 63 | YES |
| M-026740-03 | TRIM64 | TRIM64 | Tripartite motif containing 64 | YES |
| M-035527-02 | LOC642446 | TRIM64B | Tripartite motif containing 64B | YES |
| M-037583-01 | LOC646754 | TRIM64C | Tripartite motif containing 64C | YES |
| M-018490-01 | TRIM65 | TRIM65 | Tripartite motif containing 65 | YES |
| M-032288-01 | TRIM67 | TRIM67 | Tripartite motif containing 67 | YES |
| M-007007-01 | TRIM68 | TRIM68 | Tripartite motif containing 68 | YES |
| M-007137-01 | TRIM69 | TRIM69 | Tripartite motif containing 69 | YES |
| M-007077-00 | TRIM7 | TRIM7 | Tripartite motif containing 7 | YES |
| M-023459-01 | TRIM71 | TRIM71 | Tripartite motif containing 71 | YES |
| M-032293-02 | TRIM72 | TRIM72 | Tripartite motif containing 72 | YES |
| M-028896-01 | TRIM73 | TRIM73 | Tripartite motif containing 73 | YES |
| M-031736-01 | TRIM74 | TRIM74 | Tripartite motif containing 74 | YES |
| M-028282-02 | TRIM75 | TRIM75 | Tripartite motif containing 75 | YES |
| M-029945-01 | LOC390231 | TRIM77 | Tripartite motif containing 77 | YES |
| M-007076-01 | TRIM8 | TRIM8 | Tripartite motif containing 8 | YES |
| M-007174-01 | TRIML1 | TRIML1 | Tripartite motif family like 1 | YES |
| M-007182-01 | TRIP12 | TRIP12 | Thyroid hormone receptor interactor 12 | YES |
| M-006570-00 | TTC3 | TTC3 | Tetratricopeptide repeat domain 3 | YES |
| M-013785-01 | TULP4 | TULP4 | TUB like protein 4 | YES |
| M-004509-01 | UBE1 | UBA1 | Ubiquitin like modifier activating enzyme 1 | YES |
| M-005249-00 | UBE1C | UBA3 | ubiquitin like modifier activating enzyme 3 | YES |
| M-006405-01 | UBE1DC1 | UBA5 | ubiquitin like modifier activating enzyme 5 | YES |

| | | | | |
|---|---|---|---|---|
| M-006403-02 | UBE1L2 | UBA6 | ubiquitin like modifier activating enzyme 6 | YES |
| M-019759-00 | UBE1L | UBA7 | ubiquitin like modifier activating enzyme 7 | YES |
| M-009424-00 | UBE2A | UBE2A | Ubiquitin conjugating enzyme E2 A | YES |
| M-009930-00 | UBE2B | UBE2B | Ubiquitin conjugating enzyme E2 B | YES |
| M-004693-03 | UBE2C | UBE2C | Ubiquitin conjugating enzyme E2 C | YES |
| M-009387-01 | UBE2D1 | UBE2D1 | Ubiquitin conjugating enzyme E2 D1 | YES |
| M-010383-02 | UBE2D2 | UBE2D2 | Ubiquitin conjugating enzyme E2 D2 | YES |
| M-008478-02 | UBE2D3 | UBE2D3 | Ubiquitin conjugating enzyme E2 D3 | YES |
| M-009435-01 | UBE2D4 | UBE2D4 | Ubiquitin conjugating enzyme E2 D4 | YES |
| M-008850-01 | UBE2E1 | UBE2E1 | Ubiquitin conjugating enzyme E2 E1 | YES |
| M-031782-01 | UBE2E2 | UBE2E2 | Ubiquitin conjugating enzyme E2 E2 | YES |
| M-008845-00 | UBE2E3 | UBE2E3 | Ubiquitin conjugating enzyme E2 E3 | YES |
| M-009081-01 | UBE2F | UBE2F | Ubiquitin conjugating enzyme E2 2F | YES |
| M-010154-02 | UBE2G1 | UBE2G1 | Ubiquitin conjugating enzyme E2 G1 | YES |
| M-009095-01 | UBE2G2 | UBE2G2 | Ubiquitin conjugating enzyme E2 G2 | YES |
| M-009134-02 | UBE2H | UBE2H | Ubiquitin conjugating enzyme E2 H | YES |
| M-004910-00 | UBE2I | UBE2I | Ubiquitin conjugating enzyme E2 I | YES |
| M-007266-02 | UBE2J1 | UBE2J1 | Ubiquitin conjugating enzyme E2 J1 | YES |
| M-008614-02 | UBE2J2 | UBE2J2 | Ubiquitin conjugating enzyme E2 J2 | YES |
| M-009431-01 | HIP2 | UBE2K | Ubiquitin conjugating enzyme E2 K | YES |
| M-010384-01 | UBE2L3 | UBE2L3 | Ubiquitin conjugating enzyme E2 L3 | YES |
| M-008569-02 | UBE2L6 | UBE2L6 | Ubiquitin conjugating enzyme E2 L6 | YES |
| M-004348-01 | UBE2M | UBE2M | Ubiquitin conjugating enzyme E2 M | YES |
| M-003920-01 | UBE2N | UBE2N | Ubiquitin conjugating enzyme E2 N | YES |
| M-031625-01 | UBE2NL | UBE2NL | Ubiquitin conjugating enzyme E2 NL | YES |
| M-008979-01 | UBE2O | UBE2O | Ubiquitin conjugating enzyme E2 O | YES |
| M-008631-03 | UBE2Q1 | UBE2Q1 | Ubiquitin conjugating enzyme E2 Q1 | YES |
| M-008326-01 | UBE2Q2 | UBE2Q2 | Ubiquitin conjugating enzyme E2 Q2 | YES |
| M-024273-02 | FLJ25076 | UBE2QL1 | Ubiquitin conjugating enzyme E2 Q family like 1 | YES |

| | | | | |
|---|---|---|---|---|
| M-009700-02 | UBE2R2 | UBE2R2 | Ubiquitin conjugating enzyme E2 R2 | YES |
| M-009707-01 | UBE2S | UBE2S | Ubiquitin conjugating enzyme E2 S | YES |
| M-004898-01 | UBE2T | UBE2T | Ubiquitin conjugating enzyme E2 T | YES |
| M-008998-00 | UBE2U | UBE2U | Ubiquitin conjugating enzyme E2 U | YES |
| M-010064-03 | UBE2V1 | UBE2V1 | Ubiquitin conjugating enzyme E2 V1 | YES |
| M-008823-00 | UBE2V2 | UBE2V2 | Ubiquitin conjugating enzyme E2 V2 | YES |
| M-009643-03 | UBE2W | UBE2W | Ubiquitin conjugating enzyme E2 W | YES |
| M-008596-02 | UBE2Z | UBE2Z | Ubiquitin conjugating enzyme E2 Z | YES |
| M-005137-00 | UBE3A | UBE3A | Ubiquitin protein ligase E3A | YES |
| M-007197-01 | UBE3B | UBE3B | Ubiquitin protein ligase E3B | YES |
| M-007183-01 | UBE3C | UBE3C | Ubiquitin protein ligase E3C | YES |
| M-007200-00 | UBE4A | UBE4A | Ubiquitination factor E4B | YES |
| M-007202-02 | UBE4B | UBE4B | Ubiquitination factor E4B | YES |
| M-006949-01 | UBOX5 | UBOX5 | U-box domain containing 5 | YES |
| M-010691-02 | UBR1 | UBR1 | E3 ubiquitin-protein ligase UBR1 | YES |
| M-006954-01 | UBR2 | UBR2 | Ubiquitin protein ligase E3 component n-recognin 2 | YES |
| M-016653-01 | ZNF650 | UBR3 | Ubiquitin protein ligase E3 component n-recognin 3 | YES |
| M-007189-02 | EDD1 | UBR5 | Ubiquitin protein ligase E3 component n-recognin 5 | YES |
| M-008494-02 | UEVLD | UEVLD | UEV and lactate/malate dehyrogenase domains | YES |
| M-006977-01 | UHRF1 | UHRF1 | Ubiquitin like with PHD and ring finger domains 1 | YES |
| M-007117-01 | UHRF2 | UHRF2 | Ubiquitin like with PHD and ring finger domains 2 | YES |
| M-022950-01 | UNK | UNK | unk zinc finger | YES |
| M-014266-01 | UNKL | UNKL | unk like zinc finger | YES |
| M-007022-01 | VPS11 | VPS11 | VPS11 core subunit of CORVET and HOPS complexes | YES |
| M-013178-00 | VPS18 | VPS18 | VPS18 core subunit of CORVET and HOPS complexes | YES |
| M-006972-01 | VPS41 | VPS41 | VPS41 subunit of HOPS complex | YES |
| M-023668-02 | KIAA0804 | VPS8 | VPS8 subunit of CORVET complex | YES |
| M-014822-02 | WDR24 | WDR24 | WD repeat domain 24 | YES |
| M-022683-00 | WDR59 | WDR59 | WD repeat domain 59 | YES |

| | | | | |
|---|---|---|---|---|
| M-007203-01 | WDSUB1 | WDSUB1 | WD repeat, sterile alpha motif and U-box domain containing 1 | YES |
| M-013015-01 | WSB1 | WSB1 | WD repeat and SOCS box containing 1 | YES |
| M-017223-00 | WSB2 | WSB2 | WD repeat and SOCS box containing 2 | YES |
| M-004251-00 | WWP1 | WWP1 | WW domain containing E3 ubiquitin protein | YES |
| M-004252-01 | WWP2 | WWP2 | WW domain containing E3 ubiquitin protein ligase 2 | YES |
| M-004098-01 | BIRC4 | XIAP | X-linked inhibitor of apoptosis | YES |
| M-009701-01 | ZNF216 | ZFAND5 | Zinc finger AN1-type containing 5 | YES |
| M-009477-01 | ZFAND6 | ZFAND6 | Zinc finger AN1-type containing 6 | YES |
| M-007098-00 | ZNRF1 | ZNRF1 | Zinc and ring finger 1 | YES |
| M-007165-00 | ZNRF2 | ZNRF2 | Zinc and ring finger 2 | YES |
| M-010747-02 | ZNRF3 | ZNRF3 | Zinc and ring finger 3 | YES |
| M-007141-01 | ZNRF4 | ZNRF4 | Zinc and ring finger 4 | YES |
| M-007142-01 | ZSWIM2 | ZSWIM2 | Zinc finger SWIM-type containing 2 | YES |
| M-010993-01 | AIRE | AIRE | Autoimmune regulator | NO |
| M-020357-01 | BAHD1 | BAHD1 | Bromo adjacent homology domain containing 1 | NO |
| M-006901-02 | BAZ1B | BAZ1B | Bromodomain adjacent to zinc finger domain 1B | NO |
| M-020487-01 | BAZ2B | BAZ2B | Bromodomain adjacent to zinc finger domain 2B | NO |
| M-011900-01 | BRPF1 | BRPF1 | Bromodomain and PHD finger containing 1 | NO |
| M-025088-01 | BRPF3 | BRPF3 | Bromodomain and PHD finger containing 3 | NO |
| M-006976-00 | MYLIP | CALCOCO2 | Calcium binding and coiled-coil domain 2 | NO |
| M-007832-01 | CCL20 | CCL20 | C-C motif chemokine ligand 20 | NO |
| M-009774-01 | CHD4 | CHD4 | Chromodomain helicase DNA binding protein 4 | NO |
| M-009878-00 | CHD5 | CHD5 | Chromodomain helicase DNA binding protein 5 | NO |
| M-008545-01 | CXXC1 | CXXC1 | CXXC finger protein 1 | NO |
| M-031779-01 | DPF1 | DPF1 | Double PHD fingers 1 | NO |
| M-004444-00 | DPF2 | DPF2 | Double PHD fingers 2 | NO |
| M-012798-02 | DPF3 | DPF3 | Double PHD fingers 3 | NO |
| M-004012-02 | EEA1 | EEA1 | Early endosome antigen 1 | NO |
| M-014752-03 | FSD1L | FSD1L | Fibronectin type III and SPRY domain containing 1 like | NO |

| | | | | |
|---|---|---|---|---|
| M-011872-01 | HR | HR | HR lysine demethylase and nuclear receptor corepressor | NO |
| M-011697-01 | HRC | HRC | Histidine rich calcium binding protein | NO |
| M-015624-01 | INTS12 | INTS12 | Integrator complex subunit 12 | NO |
| M-011707-01 | ISL1 | ISL1 | ISL LIM homeobox | NO |
| M-017133-01 | PHF17 | JADE1 | Jade family PHD finger 1 | NO |
| M-018308-00 | PHF15 | JADE2 | Jade family PHD finger 2 | NO |
| M-013431-01 | PHF16 | JADE3 | Jade family PHD finger 3 | NO |
| M-008121-01 | AOF1 | KDM1B | Lysine demethylase 1B | NO |
| M-012458-00 | FBXL11 | KDM2A | Lysine demethylase 2A | NO |
| M-014930-01 | FBXL10 | KDM2B | Lysine demethylase 2B | NO |
| M-009899-01 | JARID1B | KDM5B | Lysine demethylase 5B | NO |
| M-010097-01 | JARID1C | KDM5C | Lysine demethylase 5C | NO |
| M-010820-01 | JARID1D | KDM5D | Lysine demethylase 5D | NO |
| M-025357-01 | KIAA1718 | KDM7A | Lysine demethylase 7A | NO |
| M-004828-02 | MLL2 | KMT2B | Lysine methyltransferase 2B | NO |
| M-007039-02 | MLL3 | KMT2C | Lysine methyltransferase 2C | NO |
| M-009670-00 | MLL4 | KMT2D | Lysine methyltransferase 2D | NO |
| M-020008-03 | KRTAP5-9 | KRTAP5-9 | Keratin associated protein 5-9 | NO |
| M-005648-01 | LGR6 | LGR6 | Leucine rich repeat containing G protein-coupled receptor 6 | NO |
| M-005338-03 | LMTK3 | LMTK3 | Lemur tyrosine kinase 3 | NO |
| M-009746-01 | LPXN | LPXN | Leupaxin | NO |
| M-019827-01 | MLLT10 | MLLT10 | MLLT10 histone lysine methyltransferase DOT1L cofactor | NO |
| M-010524-01 | MLLT6 | MLLT6 | MLLT6, PHD finger containing | NO |
| M-003281-04 | MNAT1 | MNAT1 | MNAT1 component of CDK activating kinase | NO |
| M-012796-02 | M96 | MTF2 | Metal response element binding transcription factor 2 | NO |
| M-010637-01 | NDP52 | CALCOCO2 | Calcium binding and coiled-coil domain 2 | NO |
| M-018267-01 | NLRC5 | NLRC5 | NLR family CARD domain containing 5 | NO |
| M-007048-01 | NSD1 | NSD1 | Nuclear receptor binding SET domain protein 1 | NO |
| M-006571-01 | WHSC1 | NSD2 | Nuclear receptor binding SET domain protein 2 | NO |

| | | | | |
|---|---|---|---|---|
| M-012875-00 | WHSC1L1 | NSD3 | Nuclear receptor binding SET domain protein 3 | NO |
| M-016942-00 | OIT3 | OIT3 | Oncoprotein induced transcript 3 | NO |
| M-014599-01 | WDR71 | PAAF1 | Proteasomal ATPase associated factor 1 | NO |
| M-013000-00 | PDC | PDC | Phosducin | NO |
| M-011353-00 | PHF1 | PHF1 | PHD finger protein 1 | NO |
| M-013349-00 | PHF10 | PHF10 | PHD finger protein 10 | NO |
| M-021310-01 | PHF11 | PHF11 | PHD finger protein 11 | NO |
| M-009736-01 | PHF12 | PHF12 | PHD finger protein 12 | NO |
| M-016306-00 | PHF13 | PHF13 | PHD finger protein 13 | NO |
| M-020678-01 | PHF14 | PHF14 | PHD finger protein 14 | NO |
| M-026200-02 | PHF19 | PHF19 | PHD finger protein 19 | NO |
| M-012912-01 | PHF2 | PHF2 | PHD finger protein 2 | NO |
| M-015234-02 | PHF20 | PHF20 | PHD finger protein 20 | NO |
| M-027322-02 | PHF20L1 | PHF20L1 | PHD finger protein 20 LIKE 1 | NO |
| M-006988-02 | PHF21A | PHF21A | PHD finger protein 21A | NO |
| M-007113-02 | PHF21B | PHF21B | PHD finger protein 21B | NO |
| M-014324-02 | PHF23 | PHF23 | PHD finger protein 23 | NO |
| M-014075-01 | PHF3 | PHF3 | PHD finger protein 3 | NO |
| M-014987-01 | PHF5A | PHF5A | PHD finger protein 5A | NO |
| M-014862-01 | PHF6 | PHF6 | PHD finger protein 6 | NO |
| M-019179-00 | PHF7 | PHF7 | PHD finger protein 7 | NO |
| M-004291-00 | PHF8 | PHF8 | PHD finger protein 8 | NO |
| M-016677-00 | PRICKLE1 | PRICKLE1 | Prickle planar cell polarity protein 1 | NO |
| M-023495-01 | PRICKLE2 | PRICKLE2 | Prickle planar cell polarity protein 2 | NO |
| M-013079-00 | LMO6 | PRICKLE3 | Prickle planar cell polarity protein 3 | NO |
| M-006979-01 | C6ORF49 | PRICKLE4 | Prickle planar cell polarity protein 4 | NO |
| M-006550-02 | RAPSN | RAPSN | Receptor associated protein of the synapse | NO |
| M-020374-00 | HBXAP | RSF1 | Remodeling and spacing factor 1 | NO |
| M-016355-01 | RUFY1 | RUFY1 | RUN and FYVE domain containing 1 | NO |

| | | | | |
|---|---|---|---|---|
| M-011533-00 | SCEL | SCEL | Sciellin | NO |
| M-004544-02 | SYTL3 | SYTL3 | Synaptotagmin like 3 | NO |
| M-007111-00 | SYTL4 | SYTL4 | Synaptotagmin like 4 | NO |
| M-020368-01 | FLJ10916 | THNSL2 | Threonine synthase like 2 | NO |
| M-004449-02 | TNFRSF25 | TNFRSF25 | TNF receptor superfamily member 25 | NO |
| M-021245-01 | KIAA0644 | TRIL | TLR4 interactor with leucine rich repeats | NO |
| M-006936-00 | ZMYND11 | ZMYND11 | Zinc finger MYND-type containing 11 | NO |
| M-019519-01 | ZNF185 | ZNF185 | Zinc finger protein 185 with LIM domain | NO |
| M-014202-01 | ZNF330 | ZNF330 | Zinc finger protein 330 | NO |
| M-018212-01 | ZNF547 | ZNF547 | Zinc finger protein 547 | NO |
| M-020790-01 | ZNF592 | ZNF592 | Zinc finger protein 592 | NO |
| M-008768-03 | AKTIP | AKTIP | AKT interacting protein | Catalytically dead |
| M-003549-01 | TSG101 | TSG101 | Tumor susceptibility 101 | Catalytically dead |
| M-029937-01 | LOC399937 | TRIM51BP | Tripartite motif-containing 51B, pseudogene | |
| M-029940-01 | LOC399940 | TRIM51EP | Tripartite motif-containing 51E, pseudogene | |
| M-036244-00 | LOC644006 | LOC644006 | Ring finger protein 4, pseudogene | |
| M-033531-00 | LOC440456 | PLEKHM1P1 | pleckstrin homology and RUN domain containing M1, pseudogene 1 | |
| M-026888-01 | LOC196346 | UBTFL7 | UBTF like 7, pseudogene | |
| M-184616-01 | LL0XNC01-237H1.1 | LL0XNC01-237H1.1 | Uncharacterized | |
| M-035419-01 | LOC642219 | Unmatched | Withdrawn | |
| M-035643-01 | LOC642678 | Unmatched | Withdrawn | |
| M-039499-02 | LOC652436 | Unmatched | Withdrawn | |
| M-039572-00 | LOC652591 | Unmatched | Withdrawn | |
| M-039678-01 | LOC652759 | Unmatched | Withdrawn | |
| M-039737-02 | LOC652859 | Unmatched | Withdrawn | |
| M-180259-01 | LOC653978 | Unmatched | Withdrawn | |
| M-182172-01 | LOC728919 | Unmatched | Withdrawn | |
| M-038402-01 | LOC649055 | Unmatched | Unmatched | |
| M-039811-00 | LOC653111 | Unmatched | Unmatched | |

| | | | |
|---|---|---|---|
| M-039497-01 | LOC652433 | Unmatched | Unmatched |

**Table S2. Details of *in vivo* tumorigenesis experiments.** For each figure and treatment, the number of injected animals and the number of tumors analyzed is shown. Typically, each mouse was injected twice, in both the contralateral 4$^{th}$ inguinal mammary glands, to minimize the number of animals per experiment; therefore, the number of tumors is twice the number of animals. Sometimes, however, one animal was injected only once for lack of cells. In some experiments, mice died during the observation period and their tumors were therefore excluded from the measurements. The number of tumors, considered and measured at each time point is shown, and the bold/red notation indicates the day in which a mouse was found dead.

| | | Day 0 | | Number of tumors | | | | | |
|---|---|---|---|---|---|---|---|---|---|
| Experiment | Cell type and treatment | Number of animals | Number of Tumors | Day 1 | Day 3 | Day 6 | Day 9 | Day 12 | Day 15 |
| Figure 4b | MDA-MB-361 Veh | 3 | 6 | No animals lost | | | | | |
| | MDA-MB-361 MLN | 4 | 8 | | | | | | |
| | MDA-MB-361 BTZ | 3 | 6 | | | | | | |
| | MDA-MB-231 Veh | 3 | 6 | | | | | | |
| | MDA-MB-231 MLN | 4 | 8 | | | | | | |
| | MDA-MB-231 BTZ | 3 | 6 | | | | | | |
| Figure 4f | MDA-MB-361 Veh/-DOX | 7 | 13 | 13 | 13 | **11** | **9** | 9 | 9 |
| | MDA-MB-361 Veh/+DOX | 7 | 14 | 14 | 14 | **12** | **10** | 10 | 10 |
| | MDA-MB-361 BTZ/-DOX | 6 | 11 | 11 | 11 | 11 | **9** | 9 | 9 |
| | MDA-MB-361 BTZ/+DOX | 6 | 11 | 11 | 11 | 11 | **9** | 9 | 9 |
| Figure 5b | MDA-MB-361 shCtrl Veh | 3 | 6 | No animals lost | | | | | |
| | MDA-MB-361 shCtrl DOX | 7 | 14 | 14 | 14 | 14 | **12** | **10** | 10 |
| | MDA-MB-361 shCtrl BTZ | 3 | 6 | No animals lost | | | | | |
| | MDA-MB-361 shRBX1 Veh | 6 | 12 | 12 | 12 | 12 | 12 | **10** | 10 |
| | MDA-MB-361 shRBX1 DOX | 6 | 12 | 12 | 12 | 12 | 12 | **10** | 10 |
| | MDA-MB-361 shRBX1 BTZ | 6 | 12 | 12 | **10** | 10 | 10 | **8** | 8 |
| | MDA-MB-361 shFBXW8 Veh | 6 | 12 | 12 | 12 | 12 | **10** | 10 | 10 |
| | MDA-MB-361 shFBXW8 DOX | 7 | 13 | 13 | 13 | 13 | **11** | 11 | 11 |
| | MDA-MB-361 shFBXW8BTZ | 5 | 10 | 10 | 10 | 10 | **8** | 8 | 8 |

| | | | | | | | | | |
|---|---|---|---|---|---|---|---|---|---|
| Figure 5c | MDA-MB-231 shCtrl Veh | 3 | 6 | No animals lost | | | | | |
| | MDA-MB-231 shCtrl DOX | 8 | 15 | 15 | 15 | 15 | **13** | **11** | 11 |
| | MDA-MB-231 shCtrl BTZ | 3 | 6 | No animals lost | | | | | |
| | MDA-MB-231 shRBX1 Veh | 6 | 12 | 12 | 12 | 12 | 12 | **10** | 10 |
| | MDA-MB-231 shRBX1 DOX | 6 | 12 | 12 | 12 | 12 | 12 | **10** | 10 |
| | MDA-MB-231 shRBX1 BTZ | 5 | 9 | 9 | 9 | 9 | 9 | **7** | 7 |
| | MDA-MB-231 shFBXW8 Veh | 6 | 12 | 12 | 12 | 12 | **10** | 10 | 10 |
| | MDA-MB-231 shFBXW8 DOX | 6 | 12 | 12 | 12 | 12 | **10** | 10 | 10 |
| | MDA-MB-231 shFBXW8BTZ | 5 | 10 | 10 | 10 | 10 | **8** | 8 | 8 |
| Figure 6e | T1 Veh | 5 | 9 | No animals lost | | | | | |
| | T1 BTZ | 5 | 9 | | | | | | |
| | T2 Veh | 4 | 7 | | | | | | |
| | T2 BTZ | 4 | 8 | | | | | | |
| Figure 6f | TA Veh | 4 | 8 | No animals lost | | | | | |
| | TA BTZ | 5 | 10 | | | | | | |
| | TB Veh | 5 | 9 | | | | | | |
| | TB BTZ | 5 | 9 | | | | | | |
| Figure 7c | T2 shCtrl | 4 | 8 | No animals lost | | | | | |
| | T2 shRBX1 | 5 | 9 | | | | | | |
| | T2 shFBXW8 | 5 | 10 | | | | | | |
| Figure 7f | TA shCtrl | 4 | 8 | No animals lost | | | | | |
| | TA shRBX1 | 4 | 8 | | | | | | |
| | TA shFBXW8 | 4 | 8 | | | | | | |

**Table S3. List of antibodies and working dilutions.** The antibodies used in the various experiments are shown, together with the source, the product code, the host species and the clonality. The application (WB, IP, IF or IHC) is also indicated together with the usage information and the working concentrations.

| Antigen | Source | Product code | Host | Clonality | Usage information and working concentrations | | |
|---|---|---|---|---|---|---|---|
| | | | | | WB | IF | IHC |
| Actin | Sigma | A3853 | Mouse | M | 1:1000 | | |
| GAPDH | Cell Signaling | 5174 | Rabbit | M | 1:2000 | | |
| NUMB | In house | Ref. [1] | Mouse | M | 1:1000 | 1:350 | |
| NUMB | Cell Signaling | 2756 | Rabbit | M | 1:1000 | | 1:6000 |
| Cullin 7 | Thermo Fisher | PA5-22313 | Rabbit | P | 1:1000 | | |
| Cullin 1 | Thermo Fisher | 71-8700 | Rabbit | P | 1:500 | | |
| Cullin 4A | Cell Signaling | 2699 | Rabbit | P | 1:1000 | | |
| FBXW8 | Sigma | HPA038851 | Rabbit | P | 1:1000 | | |
| RBX1 | Abcam | ab133565 | Rabbit | M | 1:1000 | | |
| SKP1 | Cell Signaling | 12248 | Rabbit | M | 1:500 | | |
| p53 (DO-1) | Santa Cruz | sc-126 | Mouse | M | 1:1000 | | |
| p21 | Cell Signaling | 2947 | Rabbit | M | 1:1000 | 1:400 | |
| p21 | Santa Cruz | sc-6246 | Mouse | M | 1:1000 | | |
| Ubiquitin | Santa Cruz | sc-8017 | Mouse | M | 1:1000 | | |
| FLAG (M2)-Tag | Cell Signaling | 14793 | Rabbit | M | 1:1000 | | |
| HA-Tag | Merk | 11583816001 | Mouse | M | 1:1000 | | |
| RFP | Abcam | ab185921 | Rabbit | M | 1:5000 | | |
| Anti-mouse IgG (H+L) HRP-Conjugate | Biorad | 1721011 | Goat | P | 1:5000 | | |
| Anti-rabbit IgG (H+L) HRP-Conjugate | Biorad | 1721019 | Goat | P | 1:5000 | | |

| | | | | | | |
|---|---|---|---|---|---|---|
| Anti-mouse IgG (H+L) Alexa647-Conjugate | Jackson Immuno Research Europe | 715-605-150 | Donkey | P | | 1:200 |
| Anti-rabbit IgG (H+L) Alexa488-Conjugate | Jackson Immuno Research Europe | 715-545-152 | Donkey | P | | 1:200 |
| VeriBlot | Abcam | ab131366 | | | 1:1000 | |

### SUPPORTING FIGURES AND LEGENDS

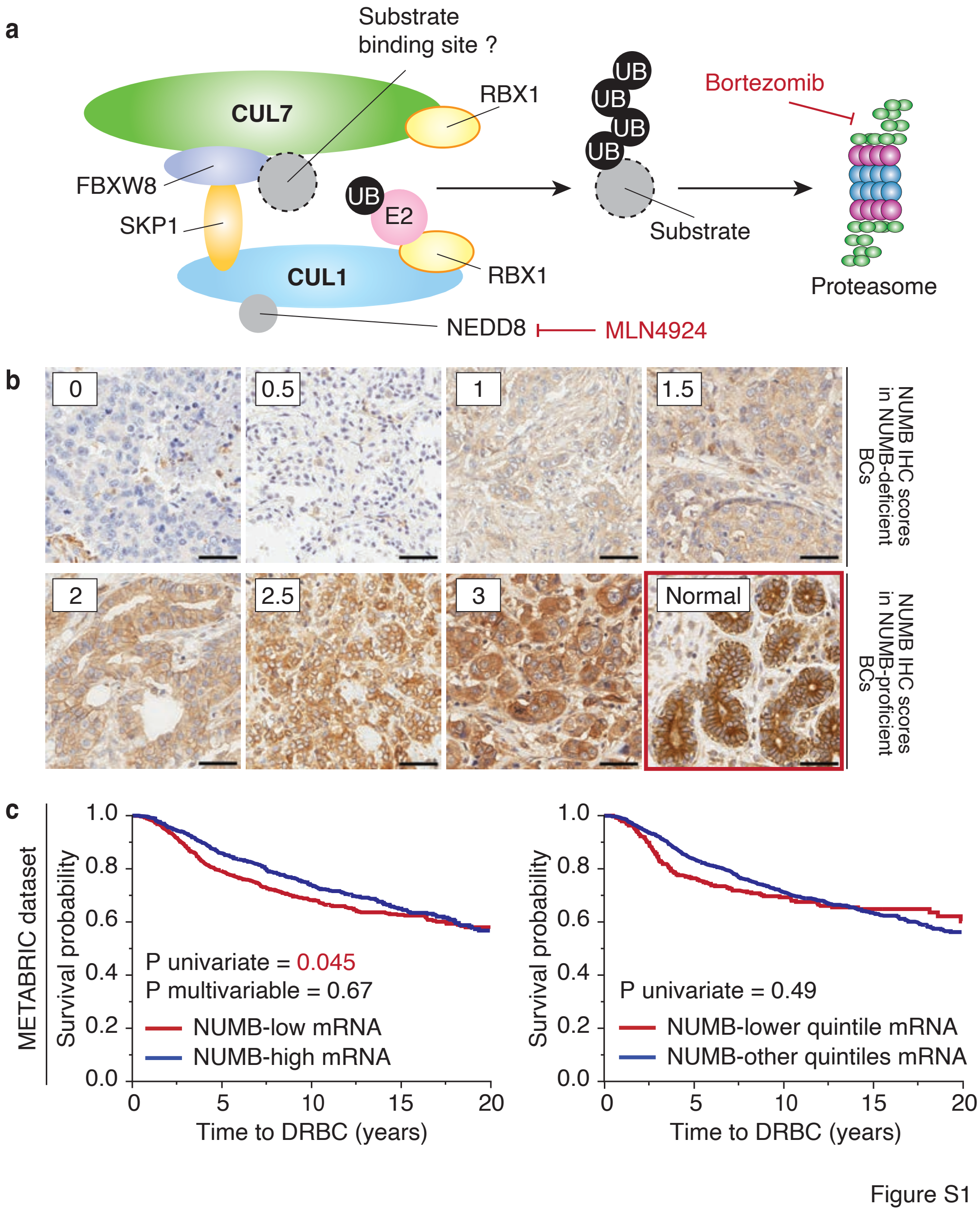

**Figure S1. Additional data to Figure 1 of the main text. a.** Cartoon depicting the putative composition and subunit arrangement of the CRL7$^{FBXW8}$ complex, as from.[4] The points of action of MLN4924 and Bortezomib are indicated. **b.** Representative images of BCs stained for NUMB in IHC. The scores used for semiquantitative evaluation and stratification into NUMB-deficient and NUMB-proficient tumors are indicated in the inset boxes. In the lower-right corner (boxed in red), an image of normal mammary gland at the margin of a tumor, displaying high NUMB intensity. Bar, 50 mm. **C.** The METABRIC BC dataset was used to analyze the possible correlation between NUMB mRNA levels and prognostic outcome (DRBC, death related to breast cancer).[5] Left panel, BCs were classified as *NUMB*-low or *NUMB*-high relative to the average *NUMB* mRNA expression across the entire cohort. A modest increase in risk was detected in the *NUMB*-low group (HR, 1.18; 95% CI, 1.00-1.38; P, 0.045). However, this correlation was not significant in multivariable analysis (P = 0.67), indicating that the modest predictive value of *NUMB* mRNA levels is not independent of other prognostic parameters. Therefore, it is likely a consequence of other alterations rather than a putative causal one. This interpretation is further supported by the analysis shown in the right panel, where tumors were stratified based on the quintiles of *NUMB* mRNA levels. In this analysis, no correlation between *NUMB* mRNA levels and prognostic outcome was observed (P = 0.49).

Kaplan-Meier, univariate and multivariable survival analyses were performed with the Survival platform and Cox proportional hazards model, as appropriate, within JMP software, version 14.3 (SAS Institute Inc., Cary, NC, 1989–2023).

a

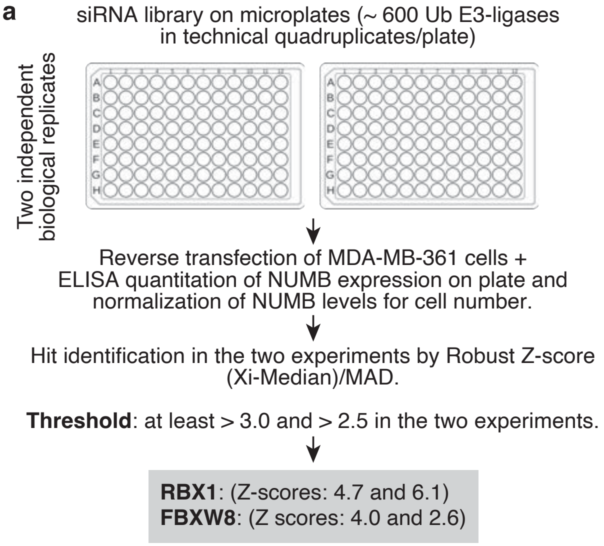

b

| Gene KD | Mod. Z-score 1st replicate | Mod. Z-score 2nd replicate |
|---|---|---|
| RBX1 | 4.67 | 6.09 |
| FBXW8 | 4.00 | 2.61 |
| TRIP12 | 4.31 | 2.09 |
| DTX1 | 3.47 | 2.29 |
| UBE2Q2 | 2.32 | 3.42 |
| UBE2G1 | 4.76 | 0.67 |
| TRIM64B | 3.31 | 1.61 |
| UBR3 | 3.77 | 1.08 |
| FBXL13 | 3.36 | 1.25 |
| FBXO38 | 3.21 | 0.12 |
| MDM2 | -0.42 | 3.39 |
| KDM7A | 3.35 | 3.11 |
| BRPF1 | 2.32 | 3.75 |
| NSD3 | 4.04 | 1.68 |
| KRTAP5-9 | 3.20 | 0.36 |

Figure S2

**Figure S2. Additional data to Figure 2 of the main text. a.** Schematic of the high-throughput screening of MDA-MB-361 cells with siGENOME® SMARTpool® siRNA Libraries (Human Ubiquitin Conjugation subset 1-3) containing ~600 siRNAs (see Experimental Section in the main text, and Table S1, Supporting Information). Two independent biological replicates of the screening were performed, in which target siRNAs were tested in quadruplicates. Seventy-two hours after reverse transfection, cell plates were processed to identify candidate hits.

For hit identification, we used a 3-step analysis:

**1) Normalization of NUMB levels to cell number**. For each plate, we determined cell viability and number (CellTiter-Fluor™), and NUMB levels (ELISA), in order to normalize NUMB levels to cell number.

**2) Determination of different scores for hit identification**. Following the normalization, we determined three different scores for each plate: the Fraction of Control, the Fraction of Sample Median and the Robust Z-score. For the Fraction of Control, each sample was divided by the mean of the control (siCTRL=1), while for the Fraction of Sample Median, each sample was divided by the median of the samples on the plate assuming that most samples have no effect. The Robust Z-score (the number of median absolute deviations (MAD) from the median) is a method that uses the screening samples as if they were negative controls, with the assumption that no biological effect is determined by most of the samples. For each plate, the Robust Z-score was calculated based on the following parameters: i) Median of samples; ii) MAD (Median Absolute deviation; MAD: = Median(|$X_i$ – median(X)|), with X representing the normalized sample values across one plate's wells and $X_i$ the sample's value at the *i* position of the plate

**3) Hit identification**: For hit identification, the Robust Z-scores were ranked. For subsequent validations, we selected hits with a Robust Z-score of ≥ 3.0 in at least one the two independent experiments and ≥ 2.5 in the other experiment (see panel b).

**B.** Selection of hits. The fifteen targets showing a Robust Z-score ≥3.0, in at least one of the two independent biological replicate screenings, are listed in the column "Gene KD". Of these only RBX1 and FBXW8 (highlighted in red) displayed a Robust Z-score ≥ 2.5 in the other experiment, and were therefore selected for further validation. Note that a number of candidates shown in this list are not connected to the ubiquitin pathway (highlighted in green, see explanation in the legend to Table S1, Supporting Information). One of these targets, KDM7A actually exceeded the threshold (≥3.0 and ≥2.5). KDM7A is a histone demethylase and was, therefore, not further considered for the purposes of the present study.

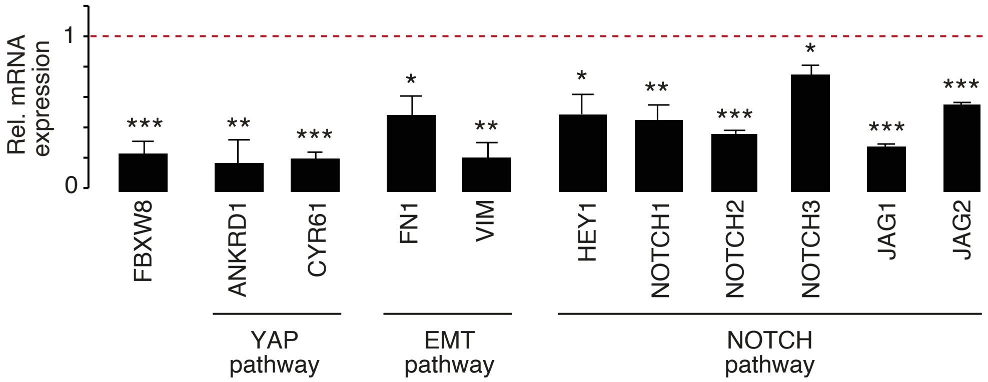

Figure S3

**Figure S3. Inhibition of the CRL7$^{FBXW8}$ complex dampens the activation of YAP, EMT, and Notch pathways.** NUMB negatively regulates the activity of the NOTCH, YAP, and EMT pathways.[6-8] Thus, the restoration of NUMB levels in a NUMB-deficient background, through the inhibition of the CRL7$^{FBXW8}$ complex, is predicted to dampen the expression of genes regulated in these pathways. We silenced FBXW8 in MDA-MB-361 cells and measured the expression of a number of genes activated in these pathways (shown at the bottom) by qPCR. Data are expressed as relative to siRNA control (dashed line). Bars represent means + SE (n=3). Statistical analysis was performed with the unpaired *t*-test, and significant differences are indicated: *, p <0.05, **, p < 0.01, *** p < 0.001.

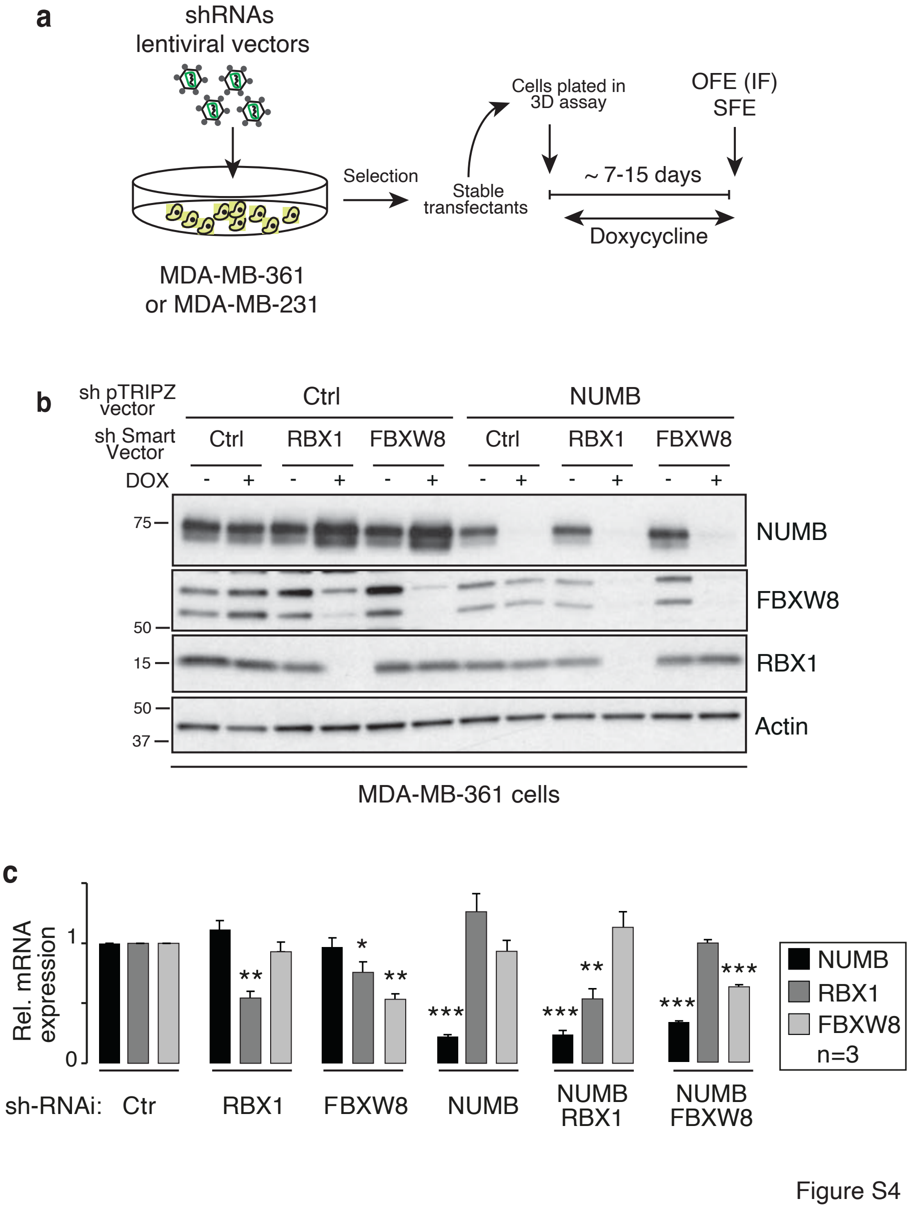

**Figure S4. Additional data to Figure 4 of the main text. a.** General scheme of the experiments performed with DOX-inducible shRNA constructs. **b.** To perform the simultaneous KD of NUMB and either RBX1 or FBXW8, and to maximize the transduction efficiency, we employed a pTRIPZ-based vector encoding shRNA-NUMB and an RFP reporter and the SMARTVector lentiviral particles encoding RBX1 or FBXW8 shRNAs and a GFP reporter (see Experimental Section). Cells were then sorted according to GFP fluorescence. Infected cells (± DOX) were IB as indicated. Actin, loading control. **c.** RT-qPCR analysis of the indicated genes (NUMB, RBX1, FBXW8) in MDA-MB-361 cells silenced or not as described (and shown in Figure 4f). Data are expressed as relative to control shRNA (Ctrl) for each condition. Bars represent the means + SE (n=3). Statistical analysis was performed with the unpaired *t*-test, and significant differences are indicated: *, p <0.05, **, p < 0.01, *** p < 0.001.

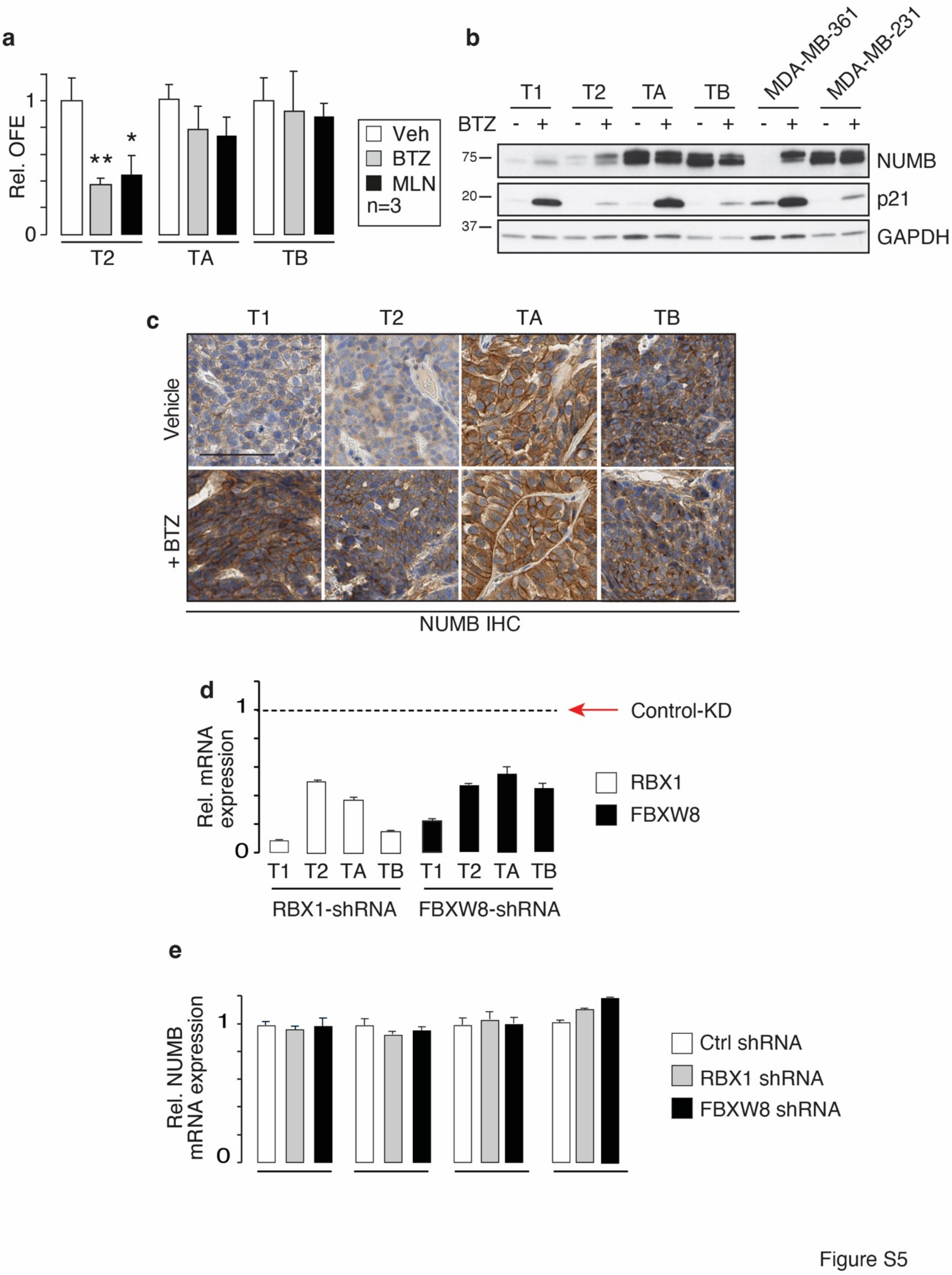

**Figure S5. Additional data to Figures 7 and 8 of the main text. a.** OFE (growth in Matrigel) of the indicated primary cultures, treated or not with BTZ (20 nM) and MLN (0.5 mM). Data are normalized to the corresponding Vehicle (Veh) control and expressed as mean + SD. Statistical analysis was performed with the unpaired t-test, and significant differences are indicated by asterisks (n=3). **b, c.** Tumors at the end of the experiments reported in Figure 7e,f of the main text were excised and analyzed by IB (panel b) and IHC for NUMB (panel c), confirming the effects of BTZ on NUMB levels throughout the experiment. Bar, 100 µm. **d, e.** RT-qPCR analysis of

RBX1/FBXW8 KD efficiency (panel d) and NUMB mRNA levels (panel e) in the variously transduced primary BC cultures used in Figure 8 of the main text. Data are the mean + SD relative to control-silenced cells (indicated by a horizontal dashed line in panel d or Ctrl shRNA in panel e) for each cell line (technical triplicates).

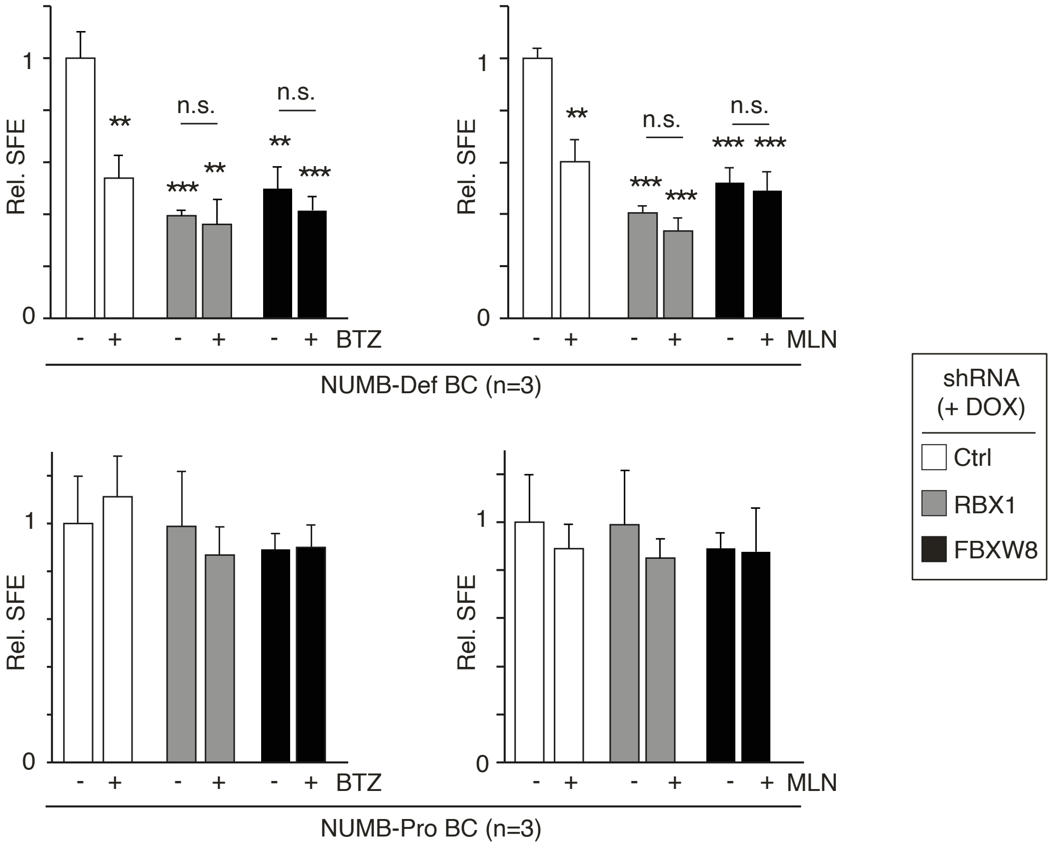

Figure S6

**Figure S6. BTZ and MLN likely act on the same pathway controlled by the CRL7$^{FBXW8}$ complex.** SFE in methylcellulose of PDX-derived primary cells transduced with the indicated pTRIPZ lentiviral vectors encoding DOX-inducible shRNA constructs, in the presence of DOX and treated or not at plating with BTZ (20 nM) or MLN (0.5 µM), as indicated. Data are normalized to the corresponding Ctrl-shRNA sample and expressed as the mean ± SD (n=3). Statistical analysis was performed with the unpaired t-test. Significant differences are indicated, *vs*. non-treated controls of control shRNA, or between the indicated pairs. NUMB-Def BC, NUMB-deficient breast cancer; NUMB-Pro BC, NUMB-proficient breast cancer.

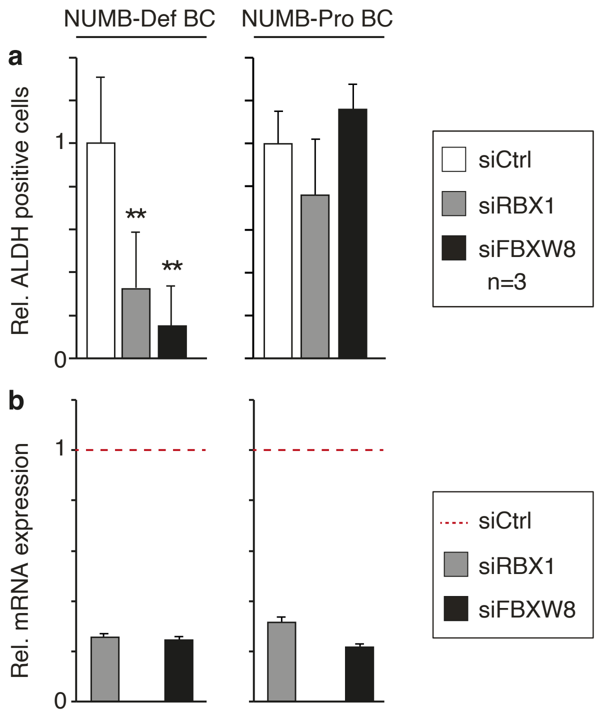

Figure S7

**Figure S7. The levels of the SC/CSC marker ALDH are decreased by the inhibition of the CRL7$^{FBXW8}$ complex in NUMB-deficient BCs. a.** ALDEFLUOR assay showing the ALDH+ cells upon RBX1 and FBXW8 silencing in NUMB-Def (left) and NUMB-Pro (right) primary tumor cells. Data are normalized to the corresponding siCtrl sample and expressed as mean ± SD (n=3). Statistical analysis was performed with the unpaired t-test. Significant differences are indicated. **b.** RT-qPCR analysis of the levels of RBX1 and FBXW8 in the same cells as in a. Data are the mean ± SD relative to siCtrl cells – indicated by the red dashed line – (technical triplicates).

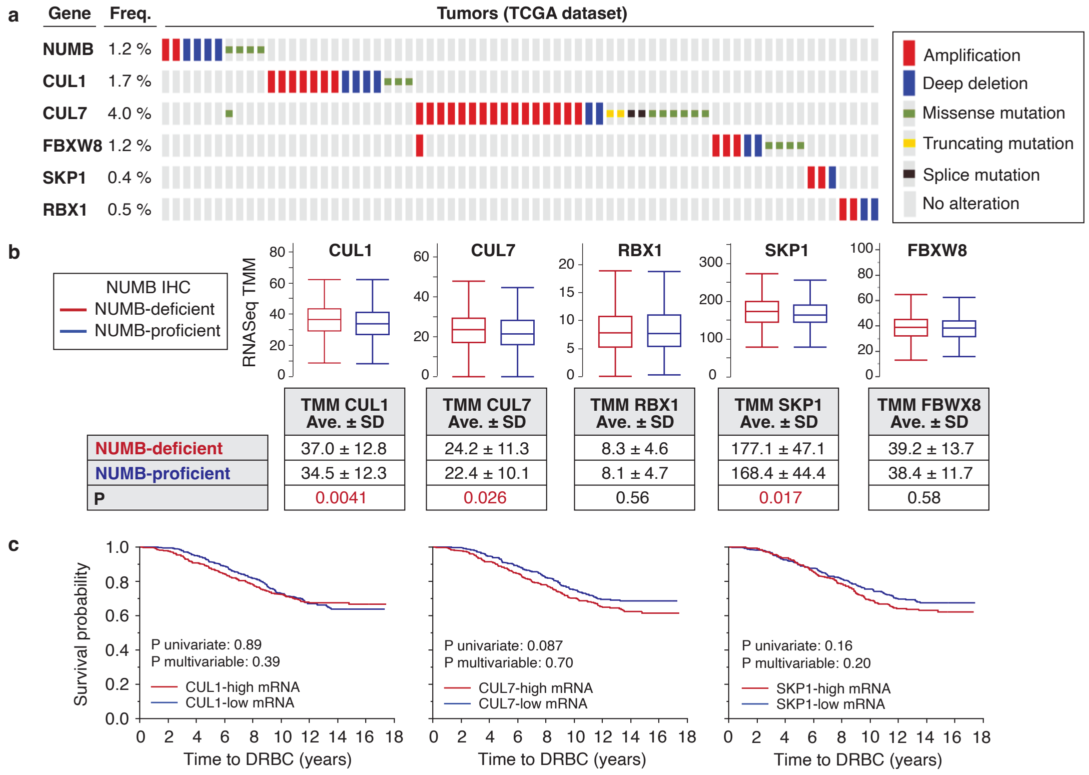

Figure S8

**Figure S8. Alterations of components of the CRL7$^{FBXW8}$ complex in BC. a.** The TCGA BC database was interrogated for mutational alterations of the *CUL1, CUL7, FBXW8, SKP1, RBX1,* and *NUMB* genes.[9] As visible, rare alterations were detected which cannot account for the frequency of NUMB loss-of-expression at the protein level herein reported (the analysis was performed on the cBioPortal, http://www.cbioportal.org/).[10] A similar analysis on the METABRIC dataset[5] (only amplifications and deep deletions available) yielded comparable results, in particular: NUMB, <1%; CUL1, 2%; CUL7, 2%; RBX1, <1%; SKP1, <1%; FBXW8, <1%. **b.** The IEO-cohort was analyzed by RNAseq and the data on *CUL1, CUL7, RBX1, SKP1,* and *FBXW8* mRNA expression were extracted. The gene TMM values in Numb-deficient and NUMB-proficient tumors, defined in Figure 1a, are shown. The analysis was performed on 716 patients for whom both reliable IHC and RNAseq data were available. P-values were calculated by the non-parametric Wilcoxon test using JMP. As shown, the mRNA levels of *CUL1, CUL7,* and *SKP1* were modestly (< 10%), albeit significantly, increased in NUMB-deficient *vs*. NUMB-proficient BCs. This difference seems too small to account for the vast differences in NUMB expression at the protein level. In further support of this notion, *CUL 1, CUL7,* and *SKP1* mRNA levels were not predictive of disease outcome in the IEO-cohort (shown in panel c). **C.** Kaplan-Meier survival analysis of patients from the IEO-cohort stratified by *CUL 1, CUL7,* and *SKP1* mRNA levels as described in b. Kaplan-Meier analyses, univariate and multivariable survival analyses were performed using the Survival platform and the Cox proportional hazards model, as appropriate, within JMP software, version 14.3 (SAS Institute Inc., Cary, NC, 1989–2023).

### REFERENCES TO SUPPORTING INFORMATION
